# MDA5 multimerization on LINE RNA drives pathogenic extracellular immune complexes in autoimmunity

**DOI:** 10.64898/2026.01.27.702129

**Authors:** Tiffany Y.-T. Hsu, Xi Wang, Jiayan Zhang, Yukari Isayama, Viktoria E. M. Schindler, Eveline van Gompel, Hadil Maadadi, Celia Torres-Arancivia, Akiko Shimazaki-Takahashi, Nina Kurihara, Mariko Kondo, Marina Galesic, Antonella Notarnicola, Sharmin Sultana, Anthony Maeda, Ruth Ann Vleugels, Paul F. Dellaripa, Takashi Yamashita, Yukiko Ito, Kentaro Awaji, Hirohito Kotani, Kazuki M. Matsuda, Begum Horuluoglu, Caroline Grönwall, Vijay Joshua, Yoshinori Ukai, Naoki Hosomi, Denisa D. Wagner, Ingrid E. Lundberg, Kazuki Kato, Sun Hur

**Affiliations:** Howard Hughes Medical Institute and Program in Cellular and Molecular Medicine, Boston Children’s Hospital, MA 02115, USA; Department of Biological Chemistry and Molecular Pharmacology, Harvard Medical School, Boston, MA 02115, USA; Division of Rheumatology, Inflammation, and Immunity, Brigham and Women’s Hospital, Harvard Medical School, Boston, MA 02115, USA; Mechanistic Immunology Research Unit, Institute of Integrated Research, Institute of Science Tokyo 113-8510, Japan; Program in Cellular and Molecular Medicine, Boston Children’s Hospital, MA 02115, USA; Division of Rheumatology, Department of Medicine, Solna, Karolinska Institutet, Stockholm, Sweden; Division of Gastroenterology, Dermatology and Rheumatology, Theme Inflammation and Aging, Karolinska University Hospital, Stockholm, Sweden; Center for Molecular Medicine, Karolinska University Hospital, Stockholm, Sweden; Perseus Proteomics Inc., Tokyo, Japan; Division of Pulmonary and Critical Care Medicine, Brigham and Women’s Hospital, Boston, MA 02115, USA; Department of Dermatology, Brigham and Women’s Hospital, Harvard Medical School, Boston, MA 02115, USA; Department of Dermatology, University of Tokyo Graduate School of Medicine, Tokyo, Japan; Department of Pediatrics, Harvard Medical School, 25 Shattuck Street, Boston, MA 02115, USA; Division of Hematology/Oncology, Boston Children’s Hospital, 1 Blackfan Circle, Boston, MA 02115, USA

## Abstract

Autoantibodies are hallmarks of many autoimmune diseases, but their potential pathogenic roles, particularly for those targeting intracellular proteins, remain unclear. Anti-MDA5-positive dermatomyositis (anti-MDA5 DM) is characterized by autoantibodies against the intracellular protein MDA5^1,2^, a conserved innate immune receptor that recognizes viral dsRNA by forming filaments^3^. Here, using four patient-derived monoclonal autoantibodies (mAbs), we reconstitute and define the molecular architecture, biogenesis, and immunological activity of pathogenic MDA5 immune complexes. Our cryo-EM analysis revealed that these mAbs bind dsRNA-scaffolded MDA5 filaments in at least two distinct binding modes, each exhibiting striking epitope convergence using germline-encoded residues. Extracellular immune complexes formed between mAbs and filamentous, but not monomeric, MDA5 potently activate multiple innate immune pathways, with the magnitude of activation determined by antibody binding mode and immune complex stoichiometry. Antibody bivalency further crosslinks MDA5 filaments into higher-order aggregates with heightened immunostimulatory activity, demonstrating an active role of autoantibodies in shaping immune complex architecture. Analysis of patient plasma reveals elevated levels of extracellular MDA5 filaments and identifies LINE retroelement-derived dsRNA as a structural scaffold. Notably, MDA5 immune complexes induce endogenous LINE dsRNA expression, likely promoting additional MDA5 filament formation and extracellular release through inflammatory cell death. These data thus support a self-amplifying inflammatory cycle as a pathogenic mechanism for anti-MDA5 DM. Collectively, our study defines a broadly applicable architectural principle, in which higher-order organization and binding modes of autoantibodies––beyond antibody affinity or nucleic acid presence alone––govern innate immune activation.

## Main text

Anti-MDA5 DM is a multisystem inflammatory disease associated with significant morbidity and mortality^4–6^. Diagnosis relies upon the presence of IgG anti-MDA5 autoantibodies and a distinctive constellation of symptoms. Patients typically present with rashes, an unusual predisposition to ulceration, and interstitial lung disease that can rapidly progress (RP-ILD) to respiratory failure^4–6^. In patients with RP-ILD, therapeutic options are limited, and disease courses are often fulminant and fatal^6,7^. At the molecular level, anti-MDA5 DM patients display strong features of systemic type I interferon (IFN) signaling, and evidence of broad dysregulation across both innate and adaptive immune cells^8–10^. Accumulating evidence suggests that anti-MDA5 antibodies contribute to disease pathogenesis^11–13^, yet the underlying mechanisms––and the precise molecular nature of the immune-stimulatory entity, including whether the antibody alone is sufficient––remain debated. Thus, anti-MDA5 DM provides an important framework for investigating how autoantibodies targeting intracellular proteins can drive immune-mediated disease.

The target antigen MDA5 belongs to the RIG-I–like receptor (RLR) family and recognizes long stretches of dsRNA that accumulate during viral replication^14^. Several viruses, including coronaviruses and picornaviruses, generate long dsRNA replication intermediates that can activate MDA5^14^. Under pathological conditions, however, shorter but abundant cellular dsRNAs, such as ∼300 bp hairpin structures formed by inverted-repeat Alu transcripts, have also been shown to activate MDA5^15–17^. This length-selective detection is mediated by MDA5 filament formation along dsRNA^18–20^. In the resting state, MDA5 is monomeric; upon engaging dsRNA, it assembles into helical filaments that recruit the dimeric E3 ligase TRIM65 via a bivalent binding mode^21^. TRIM65 then catalyzes K63-linked ubiquitination of MDA5, inducing formation of signaling-competent state of MDA5 that activates the adaptor MAVS^20^. MAVS then forms a signaling scaffold on the mitochondrial surface, activating IRF3 and NF-κB for induction of type I IFNs and pro-inflammatory cytokines^22^. While these intracellular signaling functions of MDA5 are well characterized, both in antiviral immunity and in monogenic interferonopathies^14,23^, whether MDA5 has any extracellular function has been unclear.

Here, we report an integrated biochemical, structural, and functional characterization of four monoclonal antibodies derived from anti-MDA5 DM patients and present evidence directly linking autoantibodies in complex with extracellular MDA5 filaments to disease pathogenesis in anti-MDA5 DM.

### Patient-derived mAbs bind MDA5 filaments in two distinct modes through germline-encoded residues

To investigate disease pathogenesis, we isolated antibody-secreting cells from patients with anti-MDA5 DM and cloned MDA5-reactive antibodies (Extended Data Fig. 1a). This identified two distinct MDA5-specific monoclonal antibodies (mAbs), NP6 and NP8, from one Japanese patient (Extended Data Table 1). Neither mAb cross-reacted with RIG-I or LGP2, other members of the RLR family (Extended Data Fig. 1b). Both antibodies were IgG1 isotype and utilized VH3 family-encoded heavy chains: NP6 paired IgHV3-74 with λ light chain IgLV3-1, while NP8 paired IgHV3-73 with IgLV1-47 (Extended Data Fig. 1c). Intriguingly, two additional anti-MDA5 mAbs (F01 and F12) were reported from a Swedish patient^24^, where F01 shares light chain gene usage with NP6 (hereafter class I, Fig. 1a) and F12 shares identical heavy and light chain gene usage with NP8 (class II).

**Figure 1.**
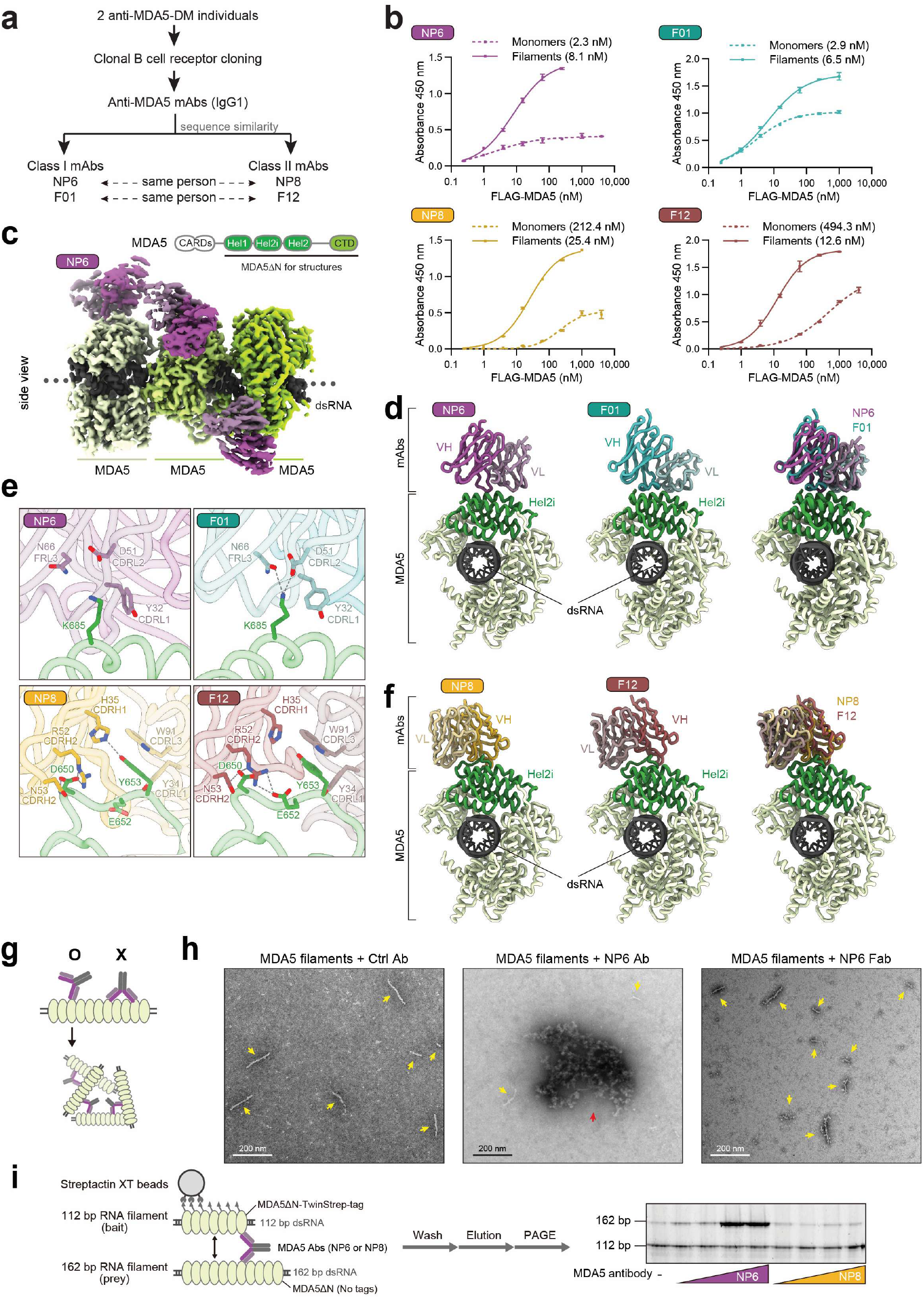
Patient-derived mAbs bind MDA5 filaments in two distinct modes. **a**) Schematic classifying monoclonal IgG1 antibodies derived from two anti-MDA5 DM individuals as class I (NP6, F01) or class II (NP8, F12) based on sequence similarity. **b**) ELISA binding curves showing class I or class II Fab reactivity to FLAG-MDA5 monomers or filaments, with apparent *K*_d_ shown in parentheses. Fab fragments were immobilized, titrated with FLAG-tagged MDA5 (full-length), followed by detection with anti-FLAG antibody. Filaments were formed by incubating FLAG-MDA5 (20 μM) with *in vitro* transcribed 512 bp dsRNA (640 nM). Data is shown as mean ± SD of 2 technical replicates. See Extended Data Figure 1d for summary table. **c**) Cryo-EM density map of the MDA5ΔN (G495R) filament formed on dsRNA in complex with NP6 scFv antibody from a side view. NP6, MDA5ΔN, and dsRNA are colored magenta, green, and gray, respectively. MDA5 filaments were computationally segmented into three monomers for reconstruction. (Upper right) Domain architecture of MDA5. **d**) Top view of the central monomeric complex of MDA5ΔN with NP6 (left), F01 (middle), and a superposition of the two mAbs (right). Hel2i subdomain is colored dark green. NP6 and F01 are colored magenta and cyan, respectively. **e**) Close-up view of MDA5 recognition by NP6 (top left), F01 (top right), NP8 (bottom left), and F12 (bottom right). Germline-encoded Ab residues interacting with MDA5 K685 (top), D650 and Y653 (bottom) are shown as stick model. The side chain of MDA5 D650 in the NP8 structure is shown at 50% transparency due to incomplete resolution. **f**) Top view of the central monomeric complex of MDA5ΔNΔ640–644Δ657–667 with NP8 (left), F12 (middle), and a superposition of the two mAbs (right). MDA5 coloring is the same as in (d). NP8 and F12 are colored yellow and red, respectively. **g**) Cartoon of the two different modes of intact antibody binding to MDA5. Structural analysis (Extended Data Figure 4a) indicates that only one Fab arm of each Ab can engage a given filament at a time (O); two Fab arms cannot simultaneously bind MDA5 molecules within the same filament (X). This geometry could enable antibodies to bridge distinct filaments, promoting inter-filament crosslinking and aggregation. **h**) Negative stain images of MDA5 filaments formed by wild-type MDA5(FL) (75 nM) ± 512 bp dsRNA (2.5 nM) bound by antibodies (18.8 nM), imaged at 30,000x. Yellow arrows indicate individual filaments, whereas red arrows indicate filament aggregates. **i**) (Left) Schematic of streptactin pull-down to examine antibody-mediated cross-bridging of MDA5 filaments. Filaments formed by MDA5ΔN tagged with TwinStrep-tag (500 nM) on 112 bp dsRNA (1 ng/μl) were incubated with untagged MDA5ΔN filaments (500 nM) assembled on 162 bp dsRNA (1 ng/μl) and mAbs (62.5-500 nM) prior to streptactin pull-down. The bound RNA was extracted and analyzed by native PAGE (right). Data are representative of independent experiments, *n* = 2 for (b), *n* = 3 for (h-i).

We next compared binding to monomeric and filamentous MDA5 using the antigen binding Fab fragments. Both NP6 and F01 Abs (class I) bound monomeric MDA5 and MDA5 filaments with comparable apparent affinity (*K*_d_ = 2-8 nM, Fig. 1b). In contrast, NP8 and F12 mAbs (class II) showed a strong preference for filamentous MDA5, binding the filament ∼8 to 38-fold more tightly than the monomeric form (apparent *K*_d_ = ∼10-20 nM for filament; apparent *K*_d_ = ∼200-400 nM for monomer, Fig. 1b).

To define the structural basis for these binding modes, we determined cryo-EM structures of MDA5 filaments on dsRNA in complex with Fab fragments or single-chain variable fragments (scFvs) of all four mAbs (Supplementary Fig. 1-6). We used the MDA5 construct lacking the N-terminal signaling domain (MDA5ΔN), which is sufficient to form filaments and displayed comparable affinity to mAbs as full-length MDA5 filament (Extended Data Fig. 1d). The gain-of-function mutation G495R that increases filament stability^25^ was used to improve filament structural homogeneity.

Structures of MDA5 bound to the class I mAbs NP6 and F01 (Supplementary Fig. 1, 2) showed that both antibodies recognized an identical epitope on the Hel2i domain of MDA5 (residues 670–690) and bound in nearly identical orientations, despite their divergent heavy chain usage and CDR3 sequences (Fig. 1c, 1d, Extended Data Fig. 1e, 2a). Mutational analysis identified Lys685 in MDA5 as the most critical residue (Extended Data Fig. 3a, 3b), which interacts with germline-encoded residues within the IgLV3-1 light chain of both NP6 and F01 (Tyr32 in CDR1, Asp51 in CDR2, and Asn66 in the framework, Fig. 1e). Similar germline-encoded lysine recognition by IgLV3-1 has been reported across multiple antibodies targeting viral epitopes^26^ (Extended Data Fig. 3c). Further supporting the importance of the germline-encoded residues, none of the somatically mutated residues directly contact MDA5 (Extended Data Fig. 2b).

In contrast, class II mAbs recognized a disordered Hel2i loop as their primary epitope, while also contacting the C-terminal domain (CTD) of an adjacent protomer (Extended Data Fig. 2c, 3a, Supplementary Fig. 3, 4), explaining the observed filament preference of class II Fabs (Fig. 1b). Because flexibility of the Hel2i loop limited local resolution, we shortened a dispensable portion of the loop (Δ640–644 and Δ657-667), enabling higher-resolution structure determination (Fig. 1f, Supplementary Fig. 5, 6). Both NP8 and F12 used germline encoded residues to recognize the Hel2i loop of MDA5, with Arg52 and Asn53 in heavy chain CDR2 interacting with MDA5 Asp650, and light chain residues Tyr34 in CDR1 and Trp91 in germline-encoded portion of CDR3 interacting with Tyr653 in MDA5 (Fig. 1e, Extended Data Fig. 2d). While some differences were also observed (e.g., MDA5 Glu652 conformation), NP8 and F12 shared key interacting residues, despite harboring distinct CDR3 insertions and somatic hypermutations (Extended Data Fig. 1e). None of the somatically mutated residues directly contact MDA5 (Extended Data Fig. 2e). Consistent with the structures, MDA5 mutations in the Hel2i loop abolished class II Ab binding (Extended Data Fig. 3a, 3d). Thus, as with class I Abs, class II Abs converge on a common MDA5 epitope through germline-encoded residues, demonstrating epitope convergence as a shared feature of both antibody classes.

### Anti-MDA5 mAbs bridge MDA5 filaments into higher-order aggregates

Having defined how antibody variable fragments recognize MDA5 filaments, we next asked how intact IgGs engage MDA5 filaments. Comparison of immobilized Fab and intact Ab binding to MDA5 filaments showed comparable affinities across all four mAbs (Extended Data Fig. 1d), indicating minimal contribution of antibody bivalency. Structural modeling further showed that two class I Fabs bound within a single filament adopt geometries and distances incompatible with an intact IgG molecule (Extended Data Fig. 4a). These observations suggest that a class I IgG bound to one filament could use its second Fab arm to engage another filament, thereby driving inter-filament crosslinking (Fig. 1g).

Consistent with this prediction, the class I mAbs NP6 and F01 induced extensive filament crosslinking, yielding large higher-order assemblies that can reach several microns in size, with few individual non-bridged filaments remaining (Fig. 1h, Extended Data Fig. 4b, 4c). In contrast, no such aggregation was observed when class I mAbs were incubated with monomeric MDA5 or when Fab fragments were incubated with MDA5 filaments (Extended Data Fig. 4b, 4c), demonstrating that both filament assembly and antibody bivalency are required for cross-bridging.

Class II Abs (NP8 and F12) also promoted filament bridging but generated substantially fewer higher-order assemblies, with many individual filaments remaining visible (Extended Data Fig. 4c), indicating less efficient crosslinking. This difference between class I and II Abs was independently confirmed by an orthogonal pull-down assay (Fig. 1i), in which Strep-tagged MDA5 filament formed on 112 bp dsRNA efficiently co-purified non-tagged MDA5 filament on 162 bp dsRNA in the presence of class I NP6 Ab, but much less so with class II NP8 Ab. One possible explanation for the superior filament-bridging efficiency of class I Abs is that they recognize a relatively rigid epitope, which may position the unoccupied Fab arm for productive engagement of another filament. By contrast, class II Abs recognize the flexible Hel2i loop, which may reduce accessibility of the second Fab arm or permit occasional bivalent engagement of a single filament.

Collectively, these results indicate that patient-derived anti-MDA5 mAbs recognize MDA5 filaments through two distinct, germline-encoded binding modes and crosslink them into higher-order assemblies, albeit with divergent efficiency.

### MDA5 filament immune complex activates type I interferon signaling in a binding mode-dependent manner

We next examined whether patient-derived anti-MDA5 mAbs can activate immune responses, and whether such activity requires immune complex (IC) formation with Ab bound to monomeric vs. filamentous MDA5. Both THP-1-derived macrophages and monocyte-derived primary macrophages were used as model systems. Using purified recombinant mAbs, MDA5 (either full-length or MDA5ΔN), and *in vitro* transcribed 512 bp dsRNA, we formed complexes in various combinations and stimulated macrophages for 4 hours. Of note, all IC stimulations were performed without the use of transfection reagents or other intracellular delivery systems.

RNA-seq analysis showed that THP-1 cells incubated with NP6 mAb + dsRNA, NP6 + monomeric MDA5, or isotype control antibody + MDA5 filament formed on dsRNA showed minimal immune signaling, compared to the mock control (Extended Data Fig. 5a, Supplementary Table 1). In contrast, NP6 + filamentous MDA5 (filament ICs) strongly induced immune-related genes, most prominently type I and II IFN signatures characteristic of anti-MDA5 DM^8–10^ (Fig. 2a, 2b). Interferon-stimulated genes such as *IFIH1* (encoding MDA5), *IFIT3*, *CXCL10*, and *MX2* were also induced (see Supplementary Table 1 for the full gene list). NP8 mAb also showed filament IC-specific immune response (Fig. 2b, Extended Data Fig. 5b, 5c), which was seen with F01 and F12 mAbs as well (Fig. 2c). Consistent with the importance of the filamentous state of MDA5 for antiviral innate immune activation, the gain-of-function mutation G495R that stabilizes filaments^25^ increased IFN-β induction (Extended Data Fig. 5d). Similarly, monocyte-derived macrophages in either M0 (M-CSF) or M1 (LPS/IFN–γ) differentiated state also showed filament-specific interferon signaling (Fig. 2d, 2e, Extended Data Fig. 5e). Both full-length MDA5 and MDA5ΔN showed comparable, filament-specific immune activation (Extended Data Fig. 5f), suggesting that MDA5 filament within the ICs do not directly activate antiviral signaling.

**Figure 2.**
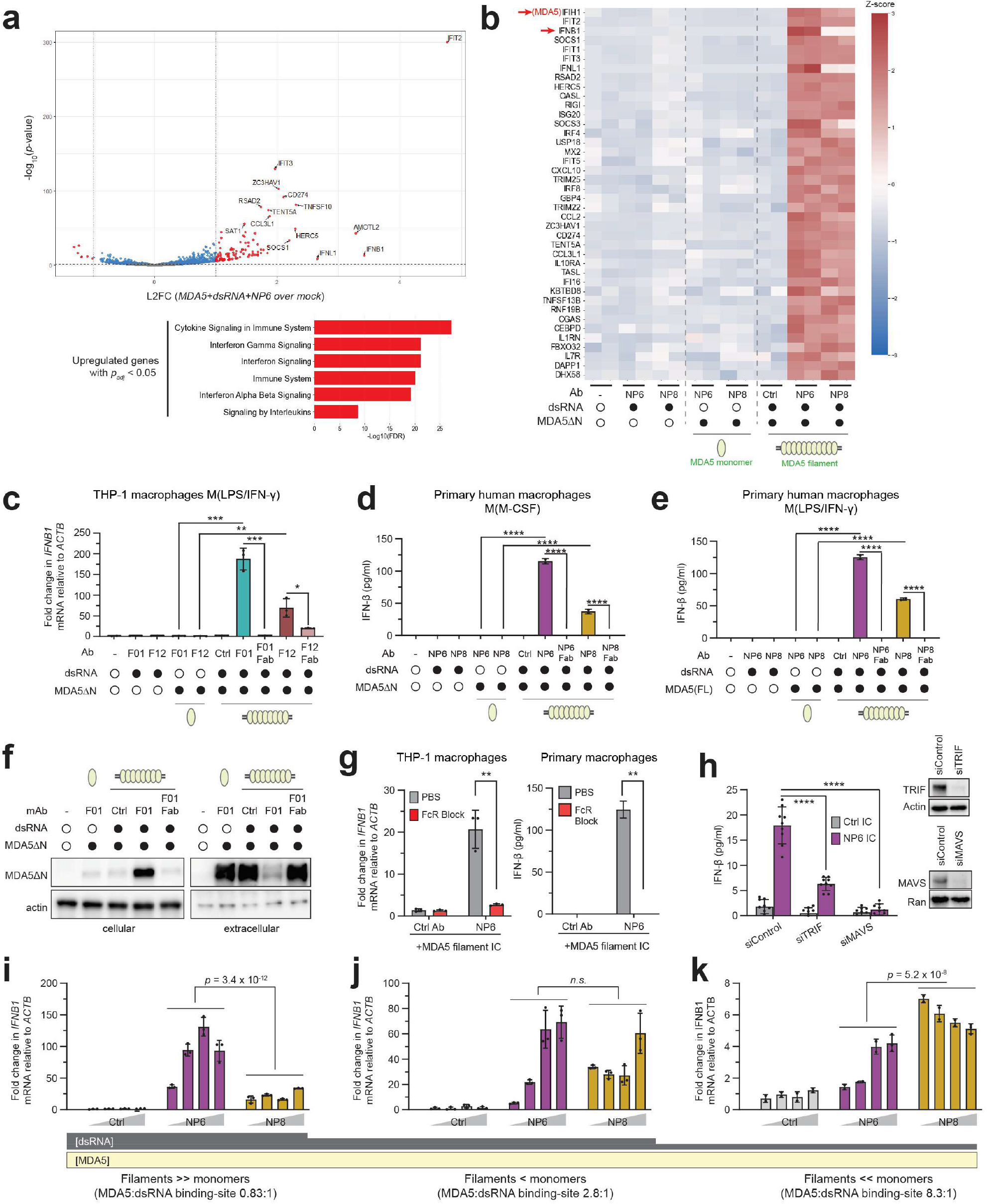
MDA5 filament immune complex activates type I interferon signaling in a binding mode-dependent manner. a) (Upper) Volcano plot of differentially expressed genes in THP-1-derived macrophages (PMA + IFN-γ/LPS) stimulated with MDA5ΔN filament-NP6 ICs over mock. All IC stimulations were performed without the use of intracellular delivery systems. Genes with log_2_-fold change |L2FC| >1 with *p* <0.05 (red) are shown; all other genes shown have *p* > 0.05 (gray) or *p* < 0.05 (blue). (Lower) Reactome pathways of significantly upregulated genes (*p_adj_* < 0.05, L2FC > 0). **b**) Heatmap of *Z*-scored expression of interferon, interferon-stimulated genes, and other immune-related genes in THP-1-derived macrophages stimulated with combinations of MDA5ΔN (90 nM monomer), 512 bp dsRNA (3 nM), mAb (NP6/NP8, 90 nM). Ctrl indicates isotype control Ab. *IFIH1* and *IFNB1* are highlighted. **c**) *IFNB1* mRNA levels in THP-1-derived macrophages stimulated with combinations of MDA5ΔN, dsRNA, mAb (F01/F12, either full-length or Fab) as in (b). Data was normalized to mock treatment. *P*-values were calculated by two-tailed Student’s *t* test. **d**) Secreted IFN-β levels measured by ELISA from healthy donor human monocyte-derived macrophages M(M-CSF) stimulated with combinations of MDA5ΔN, 512 bp dsRNA, and/or mAb as in (b). *P*-values were calculated by two-tailed Student’s *t* test. This experiment was repeated using monocytes from three different healthy donors, which showed similar patterns. **e**) Secreted IFN-β levels from monocyte-derived macrophages M(LPS/IFN-γ) stimulated with combinations of MDA5(FL), 512 bp dsRNA, and/or mAb with the same concentrations as in (b). *P*-values were calculated by two-tailed Student’s *t* test. As in (d), this experiment was repeated using monocytes from three different healthy donors. **f**) Cellular and extracellular MDA5ΔN protein levels in THP-1-derived macrophages stimulated with MDA5ΔN monomer or filament ICs as in (b). 4 hours post-stimulation, supernatant and washed cell pellet were analyzed for MDA5ΔN, which was added extracellularly. **g**) (Left) *IFNB1* mRNA levels in THP-1 macrophages pre-treated with PBS or Fc block for 1.5 hours, then stimulated with dsRNA-bound MDA5ΔN filaments (13.5 nM; 512 bp dsRNA, 0.45 nM) and NP6 Ab (13.5 nM). (Right) Secreted IFN-β levels from monocyte-derived macrophages M(M-CSF) pre-treated with PBS or Fc block for 1 hour, then stimulated with dsRNA-bound MDA5ΔN filaments (98.2 nM; 512 bp dsRNA, 3.3 nM) and NP6 Ab (98.2 nM). *P*-values were calculated by two-tailed Student’s *t* test. **h**) (Left) IFN-β ELISA using THP-1 macrophages electroporated with siRNAs prior to LPS/IFN-γ priming and stimulated with dsRNA-bound MDA5ΔN filaments (90 nM; 512 bp dsRNA, 3 nM) and NP6 Ab (90 nM). (Right) Western blot showing knockdown of TRIF and MAVS proteins prior to IC stimulation. *P*-values were calculated by two-tailed Student’s *t* test. **i-k)** *IFNB1* mRNA levels in THP-1 macrophages stimulated with ICs composed of NP6/NP8 Abs (90 nM), MDA5ΔN (90 nM), and 512 bp dsRNA [3 nM (g), 0.9 nM (h), 0.3 nM (i)]. Data was normalized to mock treatment. *P*-values were calculated by two-way ANOVA to compare NP6 and NP8 IC values across 4 mAb concentrations. Data are presented as mean ± SD of 3 biological replicates. RNA-seq results contain 2 biological repeats. All other data are representative of at least 3 independent experiments. \**P* < 0.05, \*\**P* < 0.01, \*\*\**P* < 0.001, \*\*\*\**P* < 0.0001. *n.s*, not significant.

We next investigated how MDA5 filament ICs activate the type I IFN pathway. Previous studies based on crudely purified ICs demonstrated complex uptake by Fc gamma receptors (FcγRs), delivering bound nucleic acids to endosomal TLRs for innate immune activation^27–30^. We first asked whether MDA5ΔN filament ICs are selectively internalized. Indeed, within 4 hours of addition to THP-1 culture medium, filament ICs–– but not monomeric ICs––were efficiently cleared from the extracellular compartment and instead appeared in the cellular fraction, using both F01 (Fig. 2f) and NP6 mAbs (Extended Data Fig. 5g). In contrast, F01 Fab fragments failed to induce intracellular uptake (Fig. 2f) or IFN-β production (Fig. 2c), consistent with a requirement for Fc-mediated uptake and immune activation. Similarly, fluorescently labeled MDA5, dsRNA, and Ab were internalized only when all three components were present (Extended Data Fig. 5h). FcγR blockade also inhibited filament IC-mediated interferon induction in both THP-1 and monocyte-derived macrophages (Fig. 2g), with the strongest effect observed with blockade of high affinity FcγRI (CD64) (Extended Data Fig. 5i). To determine which dsRNA sensors are involved in innate immune activation, we used CRISPR/Cas9 and siRNAs to perturb known dsRNA-sensing pathways. Loss of TLR3 (with the adaptor TRIF) and RLR (with MAVS) suppressed IC-mediated signaling (Fig. 2h, Extended Data Fig. 5j), suggesting that dsRNA within ICs can access both the endosome and cytoplasm. Interestingly, depletion of IFNAR1 also restrained *IFNB1* induction by class I and II ICs (Extended Data Fig. 5j), signifying a positive feedback loop in IFN signaling.

Because macrophage activation required all three components (MDA5, dsRNA, and Ab), we next tested how stoichiometry influences signaling output. We also compared NP6 (class I) and NP8 (class II) mAbs, which share similar filament affinities but differ in filament aggregation efficiencies (Fig. 1). The comparison showed that their signaling activities are strongly dependent on the precise filament assembly condition and antibody binding modes. When MDA5 was predominantly filamentous with a slight excess amount of dsRNA (MDA5:dsRNA binding-site ratio 0.83:1), NP6 outperformed NP8 despite similar filament-binding affinities (Fig. 2i). Similarly, F01 (class I) was significantly more potent than F12 (class II) under the same condition (Fig. 2c). These differences correlate with the enhanced filament-bridging capacity of class I mAbs, which promote the formation of higher-order crosslinked aggregates (Fig. 1h, Extended Data Fig. 4b, 4c). Consistent with the importance of filament bridging by Abs, NP6 titration revealed a bell-shaped signaling curve (Fig. 2i), likely reflecting excess antibody saturating filaments and preventing crosslinking.

In contrast, when MDA5 was largely monomeric with excess of MDA5 over dsRNA (MDA5:dsRNA binding-site ratio of 8.3:1), NP8 was more potent than NP6 (Fig. 2k), with intermediate effects observed at an intermediate ratio (2.8:1, Fig. 2j). This likely reflects sequestration of NP6 by excess monomeric MDA5, effectively diluting its ability to engage filaments. Such Ab sequestration occurs less for NP8 due to its inefficient binding to monomeric MDA5, explaining its relative dominance under monomer-rich conditions. Consistent with the notion that NP6 antibody is limiting under these conditions, NP6 titration did not result in bell-shaped dose response (Fig. 2k), in contrast to the filament-dominant condition (Fig. 2i).

Together, these data demonstrate that anti-MDA5 autoantibodies activate type I and II interferon signaling only when MDA5 autoantibodies are bound to filamentous MDA5. This activity is amplified when antibodies crosslink filaments into higher-order aggregates, but it is diminished when excess RNA-free MDA5 sequesters antibodies. This illustrates that antibody binding mode, immune complex stoichiometry and filament crosslinking potential, rather than affinity alone, determines immunological outcome.

### MDA5 filament immune complexes activate inflammatory cell death pathways

We next asked whether exogenously added MDA5 filament ICs can also activate immunological cell death pathways, given that uptake of large particulate or aggregated stimuli is known to activate inflammasomes and induce pyroptosis in macrophages^31–34^. Interestingly, genes involved in pyroptosis (including *CASP1* and *GSDMD*) as well as other cell death pathways were upregulated in THP-1 macrophages upon MDA5 filament IC stimulation (Fig. 3a), suggesting priming for inflammatory death. Immunofluorescence analyses showed increased fraction of cells with ASC specks upon stimulation of THP-1 macrophage with MDA5 filament ICs, but not with monomeric ICs (Fig. 3b, 3c). Some cells displayed one bright ASC speck, as reported with commonly used inflammasome stimuli, while others showed multiple ASC specks within individual cells (Fig. 3c). Consistent with enhanced pyroptosis, we also observed an increase in IL-1β release in THP-1 macrophages that is dependent on the NLRP3 inflammasome (Fig. 3d, 3e).

**Figure 3.**
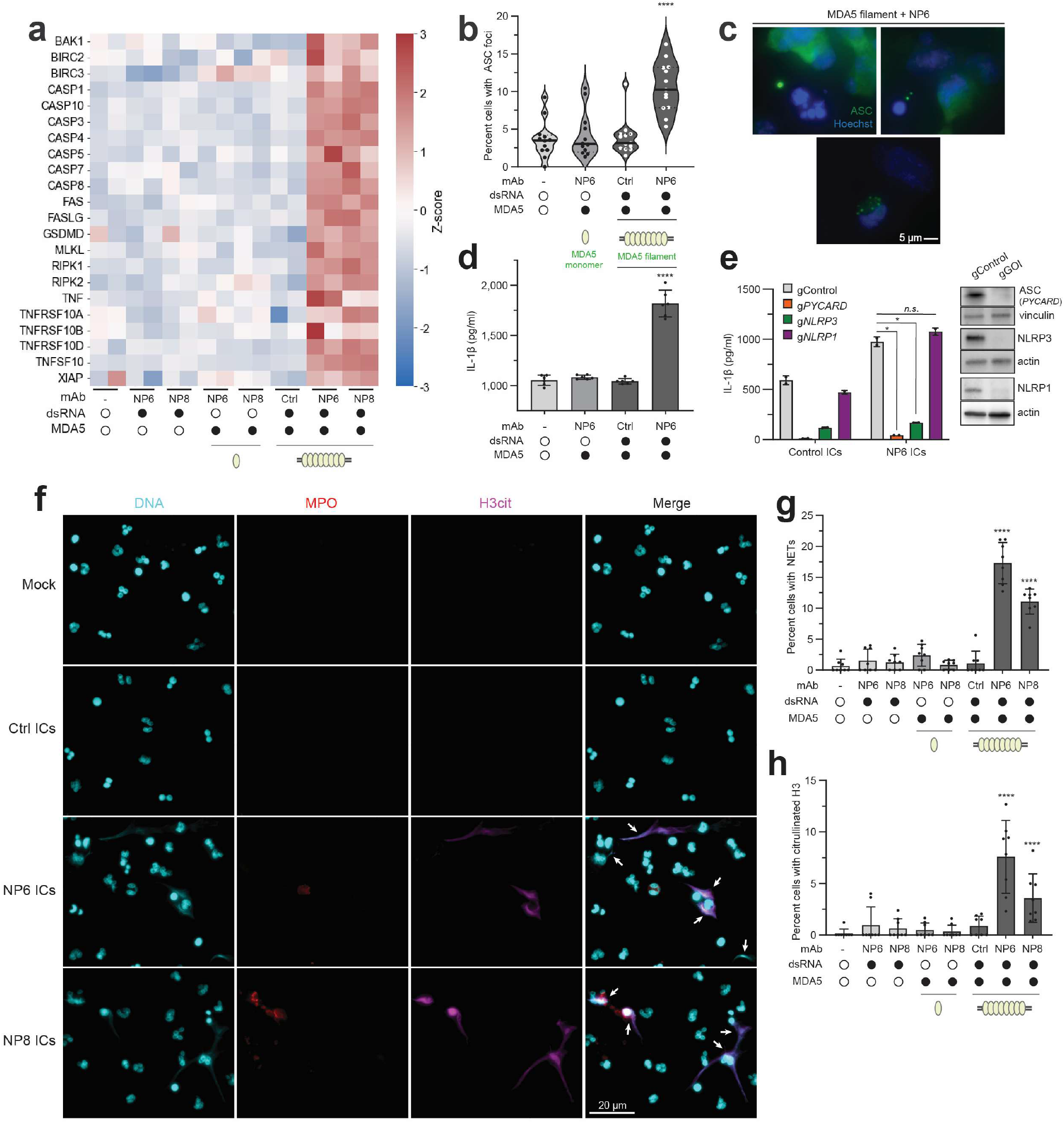
MDA5 filament immune complexes activate inflammatory cell death programs. **a**) Heatmap of *Z*-scored expression of cell death-related genes in THP-1 macrophages stimulated with combinations of NP6/NP8 Abs (90 nM), 512 bp dsRNA (3 nM), and MDA5ΔN (90 nM). **b**) ASC foci frequency in THP-1 macrophages stimulated with MDA5ΔN monomer or filament ICs as in (a). Median (solid line) and quartiles (dotted lines) are shown. *P*-values were calculated by one-way ANOVA. **c**) Immunofluorescence of ASC foci in THP-1 macrophages stimulated with MDA5ΔN filament ICs as in (a). Nuclei were stained with Hoechst 3342. **d**) Secreted IL-1β levels measured by ELISA from THP-1 macrophages stimulated with MDA5ΔN monomer or filament ICs as in (a). *P*-values were calculated by one-way ANOVA. **e**) Secreted IL-1β levels from THP-1 macrophages transduced with gRNAs targeting the indicated genes of interest (gGOI) or a non-targeting control were stimulated with MDA5(FL) filament ICs with concentrations as in (a). (Right) Western blot showing knockout of ASC, NLRP3, and NLRP1 proteins. *P*-values were calculated by two-tailed Student’s *t* test. **f**) Immunofluorescence analysis of DNA (Hoechst 3342, cyan), myeloperoxidase (MPO, red), citrullinated histone 3 (H3cit, magenta) in healthy human neutrophils stimulated with MDA5ΔN filament ICs (MDA5ΔN, 90 nM; 512 bp dsRNA, 3 nM; mAb, 20nM). This experiment was repeated using neutrophils from three different healthy donors, which showed similar patterns (see Extended Data Fig. 6a, 6b). **g**) NETosis frequency in neutrophils stimulated as in (f). NETs were defined by extracellular DNA strands or smears co-localizing with MPO and/or H3cit. Four fields of view per well were analyzed. *P*-values were calculated by one-way ANOVA comparing relevant conditions for NP6 or NP8 ICs. **h**) Quantitation of citrullinated H3-positive NETs in neutrophils stimulated as in (f). *P*-values were calculated as in (g). Data are presented as mean ± SD of at least 3 biological replicates. RNA-seq results and neutrophil assays contain 2 biological repeats. All other data are representative of at least 3 independent experiments. \**P* < 0.05, \*\**P* < 0.01, \*\*\**P* < 0.001, \*\*\*\**P* < 0.0001.

Previous studies have implicated neutrophil extracellular traps (NETs) and the associated cell death pathway NETosis in the pathogenesis of anti-MDA5 DM^11,35,36^. We thus asked whether MDA5 ICs, in particular filament ICs, can also activate neutrophils and induce NETosis. Using neutrophils freshly isolated from healthy individuals, we measured NETosis by staining extracellular DNA, myeloperoxidase (MPO), and citrullinated histone H3 (H3Cit), which are found in NETs released from neutrophils. Although basal levels of cells with NETs varied among donors (<1–8%), in all cases, filamentous MDA5 ICs robustly induced NETosis with both NP6 and NP8, whereas monomeric ICs had no measurable effect (Fig. 3f-h, Extended Data Fig. 6a, 6b). Importantly, removal of any single component of the filament IC (MDA5, RNA, or antibody) abolished NETosis induction (Fig. 3f-h, Extended Data Fig. 6a, 6b).

Together, these results demonstrate that MDA5 filament ICs also activate inflammatory cell death pathways, including pyroptosis and NETosis, highlighting their potential role in amplifying immune pathology.

### MDA5 filament ICs are enriched in anti-MDA5 DM plasma

Since MDA5 filament ICs robustly activate interferon signaling and NET release, both of which are key features of anti-MDA5 DM, we hypothesized that these filament ICs circulate in the peripheral blood of anti-MDA5 DM patients. To detect MDA5 filaments in human samples, we developed a sandwich ELISA that discriminates between monomeric and filamentous MDA5 by leveraging TRIM65’s selective recognition of filamentous MDA5^21^ (Fig. 4a). Recombinant TRIM65 was immobilized on plates, incubated with monomeric or filamentous MDA5, and then probed with anti-MDA5 antibodies (1° Ab, NP8 or F01) and anti-human secondary IgG (2° Ab, Fig. 4b). Proof-of-principle testing revealed that, regardless of the filament specificity of the 1° Ab used, TRIM65 ELISA detects MDA5 filaments with 60-80-fold higher sensitivity than monomers (Fig. 4b, Extended Data Fig. 6c), suggesting that TRIM65 binding is sufficient for filament specificity.

**Figure 4.**
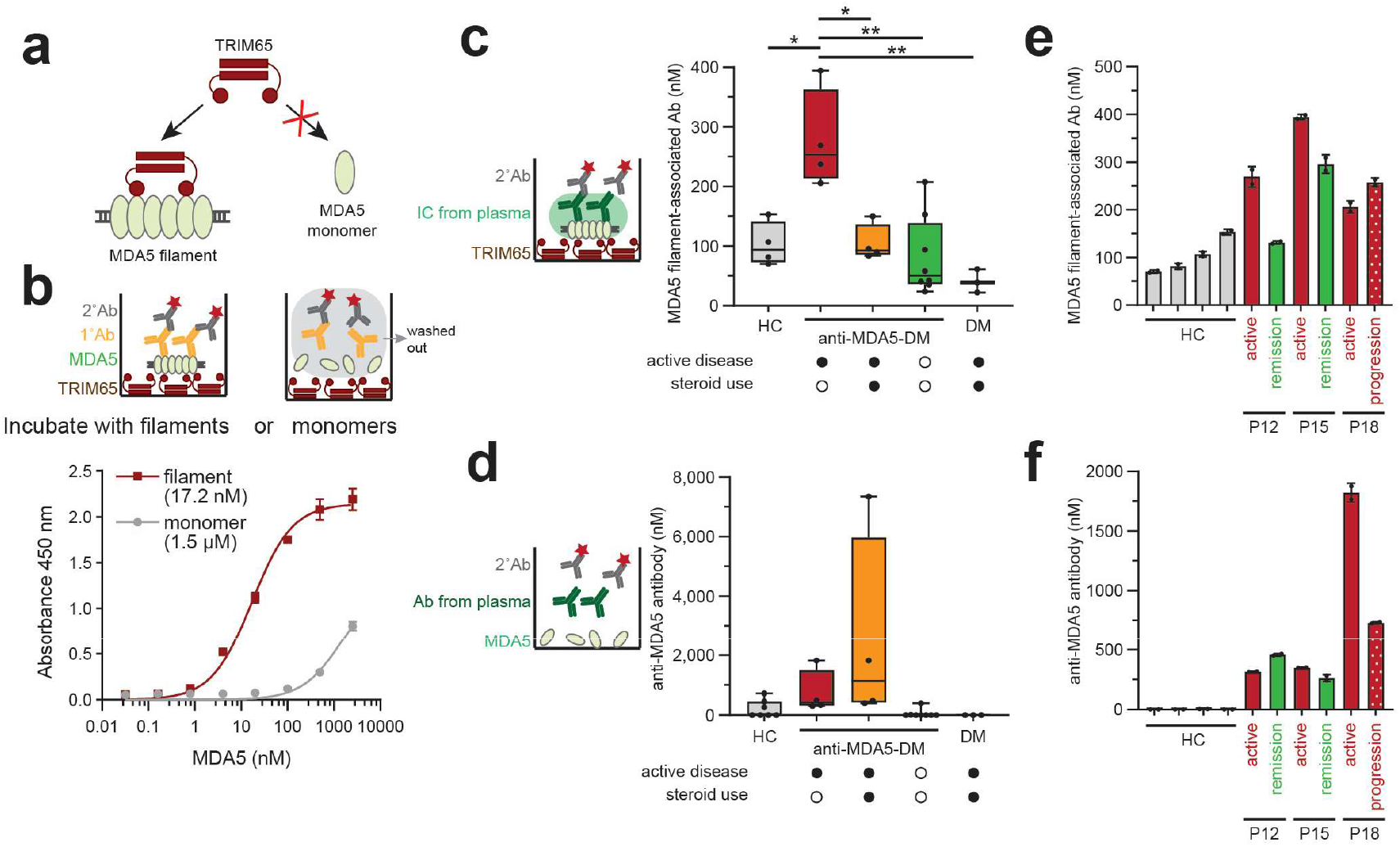
MDA5 filament immune complexes are enriched in anti-MDA5 DM plasma. a) Schematic illustrating TRIM65-mediated selective recognition of MDA5 filaments. **b**) Binding curves of TRIM65 to MDA5ΔN monomers or filaments, measured by sandwich ELISA, with apparent *K_d_* shown in parentheses (bottom). Filaments were assembled from MDA5ΔN (5 μM) and 512 bp dsRNA (160 nM). Bound filaments were detected with NP8 mAb (1°) and anti-human IgG-HRP (2°) (top). **c**) Plasma concentrations of endogenous antibodies bound to MDA5 filaments from individuals diagnosed with anti-MDA5-positive DM with active disease not on glucocorticoid therapy (*n* = 4), on glucocorticoid therapy (*n* = 4), anti-MDA5-positive DM with inactive disease (*n* = 8), anti-MDA5-negative DM (*n* = 3), or healthy controls (HC) (*n* = 4). Each data point represents an individual sample with measurements determined by TRIM65 sandwich ELISA, calibrated with a standard curve using recombinant MDA5ΔN filaments. Levels of TRIM65-captured ICs were measured directly with IgG-HRP (2°) without NP8. See Extended Data Figure 6f for ± NP8 comparison. Disease activity at the time of blood draw was defined by the treating physician and confirmed by a reviewing rheumatologist. *P*-values were calculated by two-tailed Student’s *t* test. **d**) Plasma anti-MDA5 antibody concentrations determined by direct ELISA using MDA5(FL)-coated plates, calibrated using an NP6 standard curve. See Extended Data Figure 6g. **e-f)** Plasma concentrations of MDA5 filament-bound antibody (e) and anti-MDA5 antibody (f) from healthy and anti-MDA5 DM individuals during active disease and subsequent remission (P12, P15) or progression (P18). Data are presented as mean ± SD (n *=* 2 technical replicates) from at least 3 independent experiments. \**P* < 0.05, \*\**P* < 0.01.

We examined plasma from a panel of 4 healthy controls (HCs) and 19 individuals diagnosed with dermatomyositis^37^, 16 of whom were anti-MDA5-positive by clinical testing (Extended Data Table 3). Most patients were actively treated, most commonly with intravenous immunoglobulin (IVIG)^38^ (which contains a mixture of immunoglobulins from healthy donors thought to modulate immunity by neutralizing autoantibodies, blocking FcRs, inhibiting complement, and modifying T/B lymphocytes), and/or the immunosuppressant mycophenolate mofetil (Extended Data Table 4). TRIM65 pull-down of plasma confirmed IgGs co-precipitating with MDA5 filaments, which included both full-length and truncated MDA5 species (Extended Data Fig. 6d). Since CARD domains are cleaved during cell death^39^, these truncated species may have emerged during their extracellular release.

Because circulating MDA5 filament ICs may already be bound by native anti-MDA5 antibodies, thus limiting 1° Ab access to the filaments, we performed the TRIM65 ELISA with and without 1° Ab incubation (Extended Data Fig. 6e). Surprisingly, the absorbance values were similar with or without 1° Ab (Extended Data Fig. 6f), suggesting that filament ICs in plasma were likely saturated with native anti-MDA5 antibodies. We thus omitted 1° Ab to measure the MDA5 filament-associated Ab titer as a proxy for circulating MDA5 filament level. The results showed that anti-MDA5 DM patients with active disease at blood draw had elevated levels of MDA5 filament ICs that was attenuated by glucocorticoid use (Fig. 4c). Anti-MDA5 DM patients with quiescent disease and anti-MDA5-negative individuals generally had less peripheral MDA5 filament ICs. We also determined the levels of anti-MDA5 antibody using a direct ELISA against MDA5 (Fig. 4d, Extended Data Fig. 6g) and found that these titers were also elevated in active disease, consistent with prior reports^40,41^, but did not correlate with glucocorticoid use. Notably, MDA5 filament IC levels in this group were uniformly low with minimal variability despite substantial fluctuations in autoantibody levels, raising the possibility that MDA5 filament biogenesis and/or extracellular release is steroid-responsive.

To further examine the relationship between disease course and circulating MDA5 filament ICs and autoantibodies, three anti-MDA5 DM subjects with active disease (P12, P15, P18) were followed longitudinally through disease remission or progression. In P12 and P15, clinical remission coincided with declining circulating MDA5 filament ICs despite stable autoantibody titers, whereas in P18, ICs rose despite falling titers during disease progression (Fig. 4e, 4f). These data support a close relationship between circulating MDA5 filament ICs and disease activity, consistent with their pathogenic role in anti-MDA5 DM.

### LINE-derived RNAs are enriched in anti-MDA5 DM patient plasma and associate with extracellular MDA5 filaments

Given that MDA5 filament formation requires dsRNA, the presence of MDA5 filament in patient plasma raised the question of which RNA species might drive their assembly. We performed stranded RNA-seq of total cell-free and TRIM65-purified plasma RNA, as well as total PBMC RNA, from healthy controls (HC), anti-MDA5-negative DM (with active disease), anti-MDA5 DM (active), and anti-MDA5 DM (inactive) individuals (Fig. 5a).

**Figure 5.**
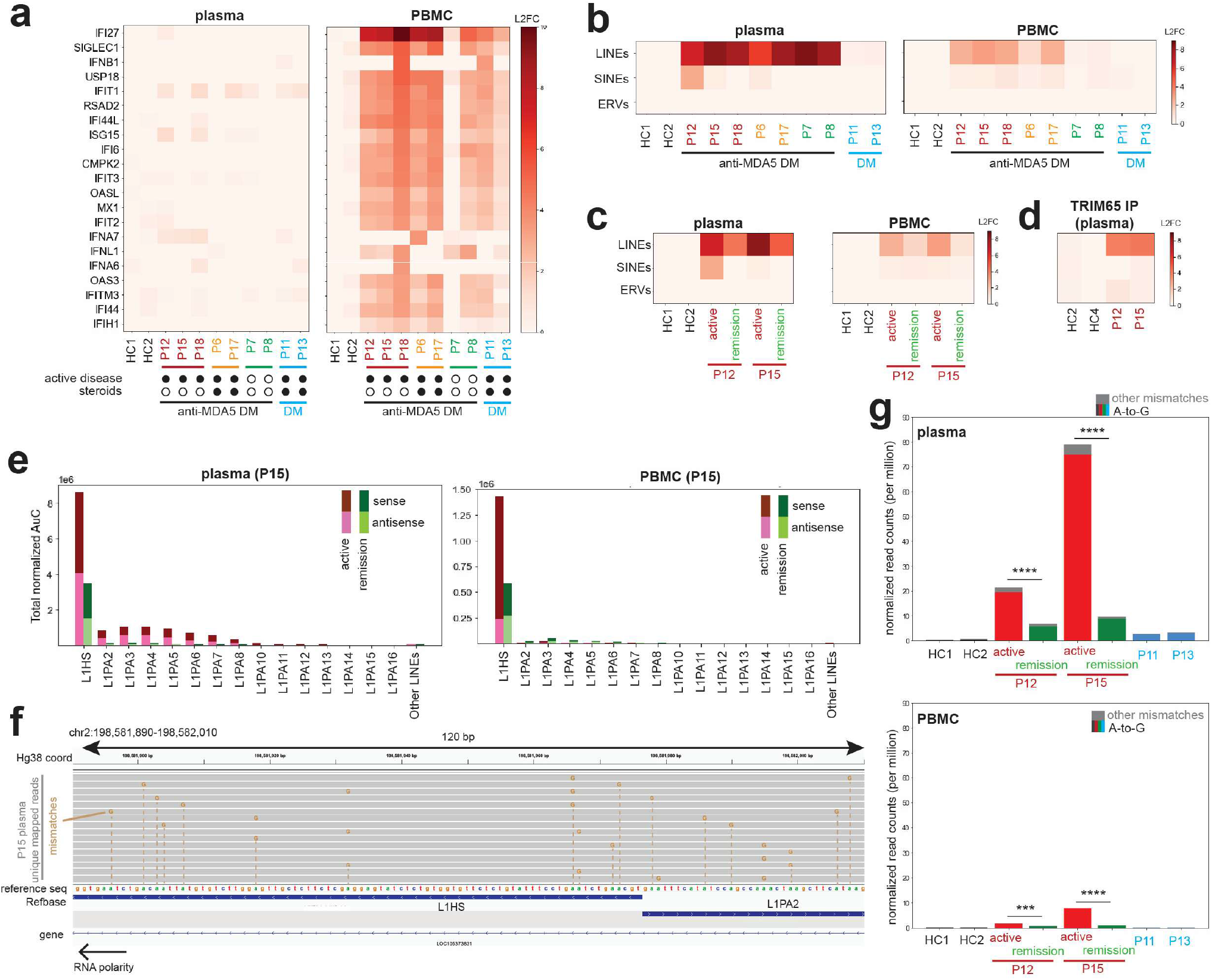
LINE-derived dsRNAs accumulate in anti-MDA5 DM during active disease. **a-b)** Heatmaps of the expression of top interferon and interferon-stimulated genes (a) and repeat elements (b) in plasma and PBMC samples. HC, healthy control. All read counts were normalized to the spike-in control. Primary mapping (random assignment among best loci) was used for multimappers. **c**) Levels of repeat elements in plasma and PBMC from individuals P12 and P15 during active disease and subsequent disease quiescence. **d**) Levels of repeat elements in plasma RNA immunoprecipitated by TRIM65 from healthy control and anti-MDA5 DM individuals with active disease. **e**) Levels of LINE-1 (L1) and all other LINE elements in plasma (left) and PBMCs (right) from individual P15 during active (red) and quiescent (green) disease. Sense (dark) and antisense (light) sequences are shown. **f**) IGV track view of uniquely mappable reads at a representative L1 locus from P15 plasma. Orange nucleotides indicate A-to-G mismatches. No other mismatches were found in these reads. **g**) Frequency of reads with A-to-G mismatches and all other mismatches within L1 regions in plasma (upper) and PBMC (lower) samples from healthy controls, anti-MDA5 DM patients during active disease and remission (P12 and P15, red and green), and anti-MDA5-negative DM patients (P11 and P13, blue). Other mismatches are shown in gray. *P*-values were calculated by two-way ANOVA with disease state and patient as factors. RNA-seq results contain 2 technical repeats except for TRIM65 RIP-seq results. \**P* < 0.05, \*\**P* < 0.01, \*\*\**P* < 0.001, \*\*\*\**P* < 0.0001.

Differential gene expression analysis of PBMC RNA confirmed a strong type I IFN signature in anti-MDA5 DM (active) patients relative to HC, consistent with previous reports^8–10^. This IFN signature was somewhat attenuated, with substantial inter-individual variability, in anti-MDA5 DM (inactive) patients and anti-MDA5-negative DM (active) patients. In contrast to the PBMC RNA, plasma RNA showed no broad IFN signature in any patient group, likely reflecting degradation of most mRNAs in the nuclease-rich plasma environment, consistent with its markedly shorter RNA size distribution (Extended Data Fig. 7a).

We next expanded our analysis beyond annotated genes to all genomic loci. This revealed long interspersed nuclear elements (LINEs), common retroelements constituting ∼17% of the genome that are normally transcriptionally silenced^42,43^, as the most enriched and abundant species in plasma from anti-MDA5 DM patients P15 and P12 relative to HCs, but not from anti-MDA5-negative DM P11 or P13 (Extended Data Fig. 7c). Unlike LINEs, short interspersed nuclear elements (SINEs), endogenous retroviruses (ERVs) (Fig. 5b), and mitochondrial genes (Extended Data Fig. 7b) showed no upregulation in patient plasma. Selective LINE enrichment was observed in plasma from all anti-MDA5 DM patients, but in PBMCs this was restricted to anti-MDA5 DM patients with active disease (Fig. 5b). Longitudinal analysis of P12 and P15 showed that LINE levels in both plasma and PBMC declined with clinical improvement (Fig. 5c). These findings were independently confirmed by TEtranscripts analysis (Extended Data Fig. 7e) and by an orthogonal library preparation method (Extended Data Fig. 7e, right).

To determine whether LINE RNAs are bound by MDA5 filaments, we performed TRIM65 pull-down followed by RNA-seq. LINE RNAs were strongly and selectively enriched in TRIM65-purified plasma fractions from P12 and P15 relative to HCs (Fig. 5d, Extended Data Fig. 7f), linking these RNAs to filamentous MDA5 complexes.

### LINE-derived RNAs form dsRNA in plasma

The enrichment of LINE RNAs in both plasma and TRIM65-purified fraction raised the question of whether they form dsRNA. Strand-specific mapping analysis of LINE subfamilies (Extended Data Fig. 7d) identified L1HS, the youngest and most transcriptionally active L1 subfamily^42,43^, as the predominant species in both PBMCs and plasma RNA from patient P15 and P12 (Fig. 5e, Extended Data Fig. 7e, 8a). In plasma, L1HS reads were present at nearly equivalent levels (sense-to-antisense ratio ∼1.1), including across regions with limited secondary structural potential (Extended Data Fig. 8b), whereas sense transcripts predominated in PBMCs (ratio ∼5.2) (Fig. 5e, Extended Data Fig. 8a). A similarly balanced strand distribution was observed in TRIM65 pull-down RNA (Extended Data Fig. 8b). This pattern is consistent with selective stabilization of sense:antisense duplexes protected by MDA5 multimerization and degradation of excess sense strands in the nuclease-rich plasma environment. Both sense and antisense LINE RNAs markedly declined after remission in P15 and P12 (Fig. 5e, Extended Data Fig. 8a), further linking LINE RNAs to disease pathogenesis.

To independently assess whether plasma LINE RNAs form duplexes *in vivo*, we quantified adenosine-to-inosine (A-to-I) editing, a hallmark of dsRNA introduced by dsRNA-specific enzyme ADAR1^3^. Because inosines are reverse transcribed as guanosines, A-to-I editing can be detected as A-to-G mismatches between sequencing reads and the reference genome (Fig. 5f). A-to-I editing was significantly elevated in active anti-MDA5 DM patients (P12 and P15) than HCs or anti-MDA5-negative DM patients (P11 and P13), declined upon clinical remission, and was higher in plasma than in PBMCs, (Fig. 5g). These are bona fide A-to-I edits rather than mapping errors or germline variants as they were present at a vastly higher rate than other types of mismatches (Fig. 5g) and were distributed stochastically across reads (Fig. 5f). Although extensive A-to-I editing can destabilize dsRNA and disrupt MDA5 filament formation^15^, the observed editing frequency on LINE RNAs (approximately 2 edits per 100 bp on average) is unlikely to preclude MDA5 recognition, given that MDA5 tolerates ∼4 bp mismatches within 112 bp dsRNA^44^. This is in contrast to IR-Alus, which contain ∼20% mismatches within the duplex region even prior to A-to-I editing^15^, which would be further destabilized by A-to-I editing.

Altogether, these data suggest that LINE-derived dsRNAs, likely formed by sense:antisense hybrids, are enriched in patients with anti-MDA5 DM, associate with MDA5 filaments in patient plasma, and decline with disease remission.

### LINE-derived RNAs are induced by immune stimulation with MDA5 filament ICs

We next examined the genomic origins of LINE-derived RNAs in anti-MDA5 DM patients. Patient-enriched, uniquely mapped PBMC LINE RNAs were categorized into 6 groups (intergenic, intron, CDR, 5’ UTR, 3’ UTR, <3kb upstream and downstream of genes), weighted by their read abundance (see Methods). Although L1HS elements in the human genome are primarily located in intergenic regions and introns (Fig. 6a, left), patient-enriched LINE reads showed a marked shift towards gene-proximal regions. In P15 and P12, 3’ UTR and upstream region together accounted for nearly 50% of both sense and antisense LINE reads and were even more prominent among highly expressed LINE loci (Fig. 6a; Extended Data Fig. 9a).

**Figure 6.**
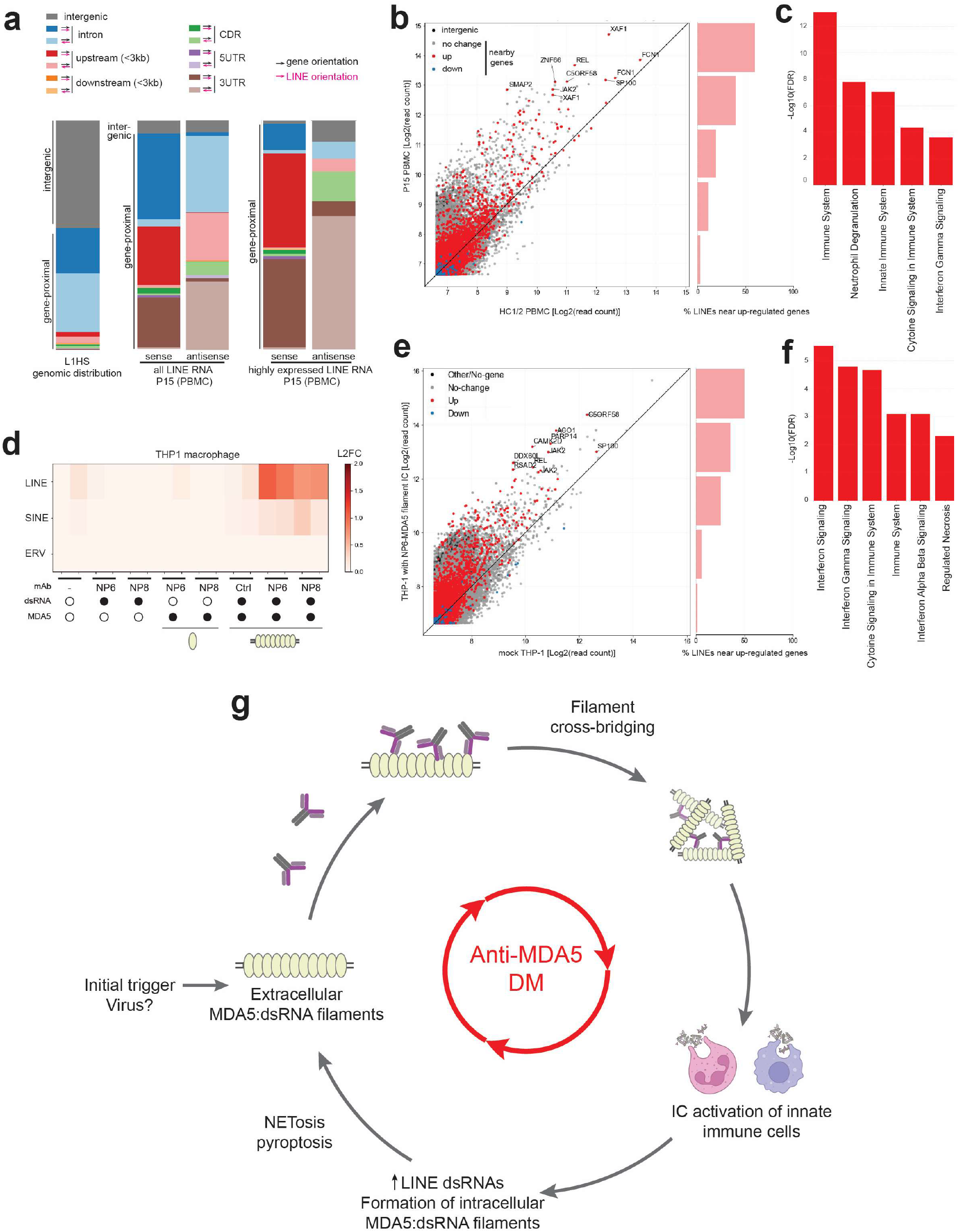
LINE-derived RNAs are induced by MDA5 immune complex stimulation. **a**) Annotation of all L1HS regions in the genome (left), all LINE regions with uniquely mappable reads weighted with read counts (middle), and all LINE regions with over 200 uniquely mappable reads, weighted with read counts (right) in P15 PBMCs. Concordant orientation of LINE and gene (darker shading) generates sense LINE RNAs upon gene transcription, whereas discordant orientation (lighter shading) generates antisense LINE RNAs. **b**) (Left) Scatterplot of spike-in-normalized read counts of individual LINE loci in P15 PBMC versus averaged read counts in HC1/2 PBMCs. Each point represents a LINE locus, grouped into those within 3 kb of an upregulated gene (L2FC > 0.3, red), a downregulated gene (L2FC < −0.3, blue), a gene with no change (|L2FC| ≤ 0.3, gray), or not within 3 kb of any gene (black). Only uniquely mappable reads were counted. (Right) Percentage of LINEs near upregulated genes in five bins. **c**) Reactome pathway of genes nearby top 100 highly expressed and upregulated LINEs in P15 PBMCs relative to HC1/2. **d**) Heatmap of L2FC in expression of repeat elements in THP-1-derived macrophages stimulated with MDA5 monomer or filament ICs as in Figure 2a. **e**) (Left) Scatterplot of reads-per-million (RPM) of individual LINE loci in THP-1 macrophages stimulated with NP6-MDA5 filament ICs versus mock control. Only uniquely mappable reads were counted. Each point is colored as in (b). (Right) Percentage of LINEs near upregulated genes in five bins. **f**) Reactome pathway of genes nearby top 100 highly expressed and upregulated LINEs in NP6-MDA5 filament IC-stimulated THP-1 relative to mock. **g**) Model of MDA5 filament IC-mediated pathogenesis of anti-MDA5 DM. Extracellular MDA5 filaments released from cells undergo antibody-mediated aggregation. These complexes induce type I interferon and pro-inflammatory signaling in innate immune cells, which upregulate expression of LINE RNA and MDA5 protein. Upregulated sense and antisense LINE RNAs form duplexes that scaffold cytoplasmic MDA5 filaments. These filaments are subsequently released extracellularly as innate immune cells activate inflammatory cell death programs like NETosis and pyroptosis. These events drive a self-amplifying feedforward loop of innate immune activation and inflammation, providing a mechanistic basis for the severe and often treatment-refractory inflammation in fulminant anti-MDA5 DM. RNA-seq results contain 2 technical repeats for patient samples and 2 biological repeats for THP-1 data. L2FC, log_2_-fold change.

Interestingly, LINE read orientation was strongly coupled to that of the host or nearby gene: LINE reads were predominantly co-directional with the host transcript, such that antisense LINE reads were enriched when the LINE orientation opposed the gene direction, and vice versa for sense LINEs (Fig. 6a, Extended Data Fig. 9a). Furthermore, the most strongly induced LINE loci were preferentially located near genes that were themselves upregulated in anti-MDA5 DM (Fig. 6b). Pathway analysis of genes near the top 100 upregulated LINEs revealed enrichment for immune-related pathways, including interferon signaling (Fig. 6c; Extended Data Fig. 9b)—a hallmark of anti-MDA5 DM^8–10^ and a signature also observed in our THP-1 model (Fig. 2b). These results suggest that LINE transcription is largely driven by host or nearby gene transcription rather than autonomous LINE promoters.

To determine whether innate immune stimulation with MDA5 filament ICs alone is sufficient to upregulate LINEs, we reanalyzed THP-1 RNA-seq data from Fig. 2a with a focus on repetitive elements. Notably, THP-1 macrophages stimulated with filamentous MDA5 ICs, but not with monomeric MDA5 ICs, showed robust upregulation of LINEs, whereas upregulation of SINEs or ERVs was minimal (Fig. 6d). As in patient PBMCs, LINE induction was strongly biased toward elements located within or near genes that were transcriptionally upregulated (Fig. 6e), many of which are again involved in innate immune signaling (Fig. 6f, Extended Data Fig. 9c).

Together, these findings indicate that LINE upregulation in anti-MDA5 DM and upon MDA5 filament IC activation is largely driven by host gene transcription, particularly innate immune programs, providing a mechanistic link between inflammation and biogenesis of endogenous dsRNA ligands.

## Discussion

Previous studies of IFN- and autoantibody-associated autoimmune diseases, such as SLE^45^, led to the prevailing model that nucleic acid-binding autoantigens in complex with autoantibodies engage FcγRs and deliver immunostimulatory nucleic acids to endosomal TLRs^27–30,46^, thus activating endogenous IFN pathways^47,48^. Yet the identity and molecular architecture of pathogenic immune complexes (ICs), and features that determine their immunological activity, remained incompletely defined.

Using four monoclonal autoantibodies isolated from anti-MDA5 DM patients, we reconstitute pathogenic ICs from fully purified components—MDA5, RNA, and antibody—enabling direct biochemical, structural, and cellular analyses. This approach reveals that RNA functions not only as a ligand for innate immune receptors but also as a structural scaffold for antigen multimerization, and that autoantibodies actively reshape antigen oligomeric state through crosslinking. As a result, antibody binding mode, stoichiometry and assembly architecture of the ICs, beyond antibody affinity, govern immune-stimulatory activity. This architectural principle is likely relevant to other autoimmune diseases involving antibodies against intracellular nucleic acid-binding proteins^49,50^. The resulting ICs are highly pathogenic in that they activate a broad range of innate immune pathways. After endocytosis of ICs via FcγRs, the dsRNA payload then activates type I and II interferon signaling via dsRNA sensors, pyroptosis via the NLRP3 inflammasome, and NETosis via PAD4. Released proinflammatory cytokines would drive leukocyte recruitment and local tissue inflammation, while NETs may impair wound healing. These observations together provide a mechanistic explanation for the inflammatory lung disease and cutaneous ulceration^8–10^ frequently seen in anti-MDA5 DM.

Analysis of RNA in patient plasma and TRIM65-purified fraction identifies LINE-derived dsRNA as a key extracellular component of pathogenic MDA5 ICs. Unlike other disease contexts in which autonomous LINE activation reflects epigenetic dysregulation or promotes genomic instability and cGAS–STING signaling through LINE-encoded enzymatic activities^51,52^, LINE RNAs in anti-MDA5 DM arise largely through host or nearby gene transcription and function through stable dsRNA formation. Because many induced LINE loci are associated with inflammatory genes, MDA5 ICs can further amplify LINE dsRNA expression. Together with IFN-driven upregulation of MDA5, this LINE induction would likely promote additional filament formation and their extracellular release through inflammatory cell death, establishing a self-amplifying feed-forward loop that drives disease propagation (Fig. 6g). However, it is important to note that LINE induction is not a universal consequence of inflammation and was not observed in other IFN-driven diseases, including anti-MDA5-negative DM in this study and SLE^53–55^. What factors distinguish anti-MDA5 DM from other diseases remains an important question.

The initiating trigger of the pathogenic vicious cycle––the source of initiating dsRNA–– also remains uncertain. Viral dsRNA represents one plausible trigger and may help explain the pulmonary symptoms, seasonal disease flares, and the detection of anti-MDA5 antibodies in subsets of COVID-19 patients^6,10^. However, we did not detect enrichment of viral RNA from anti-MDA5 DM patient plasma, which may indicate that persistence of the initiating stimulus may not be required once the inflammatory circuit is established.

Finally, our structural analyses reveal striking epitope convergence across unrelated patients. While class I and II antibodies recognize distinct epitopes, two antibodies within each class converged on the same respective epitope through germline-encoded antibody residues despite divergent heavy chain CDR3 sequences and somatic hypermutations. These two independent examples suggest that MDA5 harbors epitopes with an intrinsic predisposition to breach B cell tolerance, reminiscent of germline-driven immunodominance described in antiviral immunity^26^.

Altogether, our findings define anti-MDA5 DM as a “perfect storm” in which both innate and adaptive immune tolerance are breached, resulting in a self-sustaining pathogenic inflammatory loop. More broadly, this work provides a molecular architectural framework for understanding how intracellular nucleic acid-binding proteins can become potent extracellular immunogens.

## Methods

### Material Preparation

#### Guide RNA design and virus production

For experiments using knockout cell lines, guide RNA (gRNA) sequences were designed either manually or using the online tool “CRISPR-Cas9 guide RNA design checker” from IDT. gRNA sequences are listed in Supplementary Table 2. Synthetic gRNA oligonucleotides were cloned into LentiCRISPRv2 at BsmBI restriction sites^1^. Lentivirus was produced by transiently transfecting constructs using Mirus Bio TransIT transfection protocols into 293T cells and collecting viral supernatants 48 hours after transfection.

#### Cell lines and primary cells

THP-1 cells (gift from Dr. J. Kagan, Harvard Medical School) were maintained in RPMI-1640 (GE Healthcare) supplemented with 10% heat-inactivated fetal bovine serum (FBS). Monocyte-derived macrophages were maintained in RPMI-1640 supplemented with 10% FBS. Cells were grown at 37°C in a 5% CO_2_ humidified atmosphere. Cells were routinely tested for mycoplasma contamination. Stable knockout THP-1 cell lines were generated by lentiviral transduction in the presence of 8 μg/ml polybrene followed by selection with 2 μg/ml puromycin. To generate THP-1 *MAVS* KO, cells were sequentially transduced with lentivirus encoding *MAVS* gRNA1 and gRNA2 (Supplementary Table 2), then selected with puromycin.

#### Protein expression and purification

MDA5 full-length (FL), MDA5Δ1-286 (MDA5ΔN), and LGP2 proteins were expressed in Sf9 cells using the Bac-to-Bac baculovirus expression systems (Thermo Fischer Scientific). Sf9 cells were infected in Sf-900 culture media (Thermo Fischer Scientific) for 48-60 h at 27°C. Cells were harvested, lysed by sonication, and clarified by centrifugation. The supernatant was applied to Ni-NTA agarose (QIAGEN) and the eluted protein was treated with HRV 3C protease at 4°C overnight to cleave the N-terminal tag. The proteins were further purified by affinity chromatography using a HiTrap Heparin column (Cytiva), followed by size-exclusion chromatography on a Superdex 200 Increase 10/300 column (Cytiva).

FLAG-MDA5(FL), FLAG-MDA5ΔN, MDA5ΔN, RIG-I(FL), RIG-IΔN, and their variant proteins were expressed in *Escherichia coli* Rosetta2 (DE3) (Novagen) by induction with 0.25 mM isopropyl β-D-thiogalactopyranoside (IPTG) at 18°C overnight. Cells were lysed by Emulsiflex or sonication and clarified by centrifugation. N-terminally His_6_-SUMO-tagged proteins were purified using Ni–NTA agarose chromatography. The eluted proteins were incubated with Ulp1 protease at 4°C for 4 h to cleave the His_6_-SUMO-tag. The proteins were further purified by chromatography on HiTrap Heparin and Superdex 200 Increase 10/300 columns.

MDA5, SNAP-MDA5ΔN, and their variant proteins were expressed in *E. coli* BL21(DE3) (Novagen) with groEL chaperones by induction with 0.5 mM IPTG at 20°C overnight. Cells were lysed by Emulsiflex and clarified by centrifugation. N-terminally His_6_-NusA-tagged MDA5 were purified using Ni-NTA agarose chromatography, and the eluted proteins were purified by chromatography on Superdex 200 Increase 10/300 column. Appropriate fractions were treated with HRV 3C protease at 4°C overnight to cleave the His_6_-NusA-tag. The proteins were further purified by chromatography on HiTrap Heparin and Superdex 200 Increase 10/300 columns.

Strep-HA-TRIM65^CC-PSpry^ (residues 125-517) protein expression and lysis was performed in the same manner as MDA5, MDA5ΔN, and LGP2 in Sf9 cells, while protein purification was done by chromatography on Strep-Tactin Superflow (IBA). The eluted protein was treated with HRV 3C protease and further purified by size-exclusion chromatography on Superdex 200 Increase 10/300 columns.

All MDA5 antibodies were expressed in Expi293 cells (Thermo Fischer Scientific). Culture supernatants were collected by centrifugation 5 days post-transfection. IgG antibodies were purified using a rProteinA column (Cytiva), followed by size-exclusion chromatography on Superdex 200 Increase 10/300 column. scFv of MDA5 antibodies were purified using Ni-Sepharose Excel column (Cytiva) and Superdex 200 Increase 10/300 column. Fab fragments of MDA5 antibodies were prepared with papain digestion at 1 mg/ml IgG and IgG:papain mass ratio of 20:1 in PBS with 10 mM EDTA and 10 mM L-Cysteine at 37°C for 1 hour. Digestion was quenched with 50 mM iodoacetamide. After buffer exchange with 50 mM Tris-HCl (pH 7.0), Fabs were purified with anion exchange chromatography (1 ml HiTrap SP FF, GE Healthcare) at 1 ml/min flow rate in Tris-HCl (pH 7) with a gradient in the same buffer to 1 M NaCl in 30 ml.

For fluorescein labeling of NP6, the light chain sequence was C-terminally tagged with LPETGG and labeled with a 0.5 mM fluorescein-conjugated peptide AZDye 647 Gly-Gly-Gly (Vector labs) using 0.3 uM *Staphylococcus aureus* sortase A 5M (a gift from Dr. H.L. Ploegh, Boston Children’s Hospital). Subsequently the reaction mixture was passed through a Ni-NTA column and further purified using Superdex 200 Increase 10/300 column.

All purified proteins were flash-frozen in liquid nitrogen and stored at −80°C until use. All proteins used in cell-based experiments were tested for endotoxins at < 0.1 EU/ml using a chromogenic endotoxin quant kit (Pierce).

#### RNA preparation

dsRNAs were prepared by *in vitro* T7 transcription as previously described^2^. The sequences of 112, 162, 512, and 1012 bp dsRNAs were taken from the first 100, 150, 500, and 1000 bp of the MDA5 gene flanked by 5’-gggaga and 5’-tctccc, respectively. The two complementary strands were co-transcribed, and the duplex was separated from unannealed ssRNAs by 10% acrylamide gel electrophoresis in TBE buffer. RNA was gel-extracted using the Elutrap electroelution kit (Whatman), ethanol precipitated, and stored in 20 mM HEPES, pH 7.0

For 3’-fluroescent labeling of RNA, the 3’ end of RNA was oxidized with 0.1 M sodium periodate in 0.1 M NaOAc, pH 5.5, at room temperature (RT) for 2 h. The reaction was subsequently quenched with 0.2 M KCl, and the reaction buffer was exchanged with 0.1 M NaOAc, pH 5.5, using Zeba desalting column (Pierce). Oxidized RNA was incubated with 1 mM Cy3-monohydrazide (Amersham) at RT for 4 h. The labeled RNA was desalted twice to remove unincorporated dye and stored in 20 mM HEPES, pH 7.0, at −20°C.

### Collection of patient samples

For anti-MDA5 antibody cloning to generate NP6 and NP8, as outlined in Extended Data Figure 1a, peripheral blood mononuclear cells (PBMCs) and plasma were collected from a Japanese patient with positive anti-MDA5 antibody (by clinical ELISA testing) and rapidly progressive interstitial lung disease (RP-ILD) prior to treatment. The patient met the Sontheimer criteria for clinically amyopathic dermatomyositis (CADM)^3^. The study was conducted in accordance with the Declaration of Helsinki and was approved by the Ethics Committee of the University of Tokyo Graduate School of Medicine. Written informed consent was obtained from all participants.

For anti-MDA5 antibody cloning to generate F01 and F12, PBMCs and sera were collected from 3 anti-MDA5-positive patients diagnosed between 1999 and 2021 at Karolinska University Hospital (Stockholm, Sweden), as reported separately^4^.

PBMCs and plasma samples in Figures 4-6 were collected between March 2022 and December 2025 from patients diagnosed with dermatomyositis (DM) at Brigham and Women’s Hospital (Boston, USA). Dermatomyositis diagnosis was made in accordance with EULAR/ACR criteria^5^ and clinical designation of anti-MDA5-positivity was determined by clinical testing by the Oklahoma Medical Research Foundation Myositis profile. The study was performed with informed consent in accordance with protocols approved by the institutional review boards at Mass General Brigham and Boston Children’s Hospital. Clinical variables of interest regarding the DM diagnosis were extracted from the electronic health record by manual chart review. These included the subject’s age, sex, race, ethnicity, DM duration, autoantibody status, presence of muscle weakness, DM skin findings, ILD severity, inflammatory arthritis, and DM immunomodulatory medications (including the specific dose of any glucocorticoid when applicable). CK and aldolase were measured using commercial assays in the clinical laboratory at Brigham and Women’s Hospital. Disease activity level was determined by the global assessment from the last rheumatology provider note documented in the electronic health record. ILD severity was graded based upon fulfillment of the following parameters: subclinical, undiagnosed or asymptomatic with <5% involvement on CT; mild, minimally symptomatic but not requiring treatment, may have seen a pulmonologist but does not require frequent management with <10% involvement on CT; moderate, symptomatic and on treatment requiring regular pulmonology follow up with >10% involvement on CT; and severe, symptomatic and on treatment, with regular pulmonology follow up, on supplemental oxygen or with flares requiring hospitalization or treatment escalation.

PBMC samples from healthy donors in Figure 2 were obtained from discarded specimens and received ethical approval from the institutional review board at Brigham and Women’s Hospital.

### Isolation of human plasma, PBMCs, and neutrophils

Plasma and PBMCs were isolated from the same anticoagulated whole-blood sample collected in EDTA tubes. Whole blood was centrifuged at 1,500 x *g* for 10 minutes at room temperature, and the plasma supernatant was carefully collected and stored at −80°C until further analysis. The remaining cellular fraction was diluted 1:1 with phosphate-buffered saline (PBS) and layered onto a Lymphoprep Tube (Abbott Diagnostics Technologies AS, Oslo, Norway) or Ficoll-Paque (Cytiva). Density-gradient centrifugation was performed at 800 x *g* for 30 minutes at RT with the brake disengaged. The PBMC layer at the plasma–gradient interface was collected, washed twice with PBS (250 x *g* for 10 minutes), treated with 1x RBC lysis buffer (Invitrogen), and resuspended for downstream immunological assays. Cell number and viability were assessed using trypan blue exclusion.

Neutrophils were collected from 3 ml of peripheral blood collected from healthy controls in EDTA tubes. Neutrophils were isolated using Histopaque-1119 (Sigma-Aldrich) and Percoll Plus (GE Healthcare) gradients as described previously^6^, which is a method that causes minimal activation of neutrophils during isolation. Purity of cells was >95% as determined by Wright-Giemsa staining.

### Generation of patient-derived NP6 and NP8 mAbs

#### Single-cell isolation of anti-MDA5 antibody secreting cells

PBMC (1 x 10^6^ cells) obtained from an anti-MDA5 antibody-positive CADM patient were loaded onto microchambers^7^ (20 μm in diameter; 196,000 microwell array on a chip) that had been pre-coated with 5 μg/ml MDA5 antigen. Single-cell analysis was performed using an automated single-cell analysis and isolation system, which enables high-throughput screening with a microchamber array-based single-cell picking platform (AS ONE, Osaka, Japan). Antibody secretion from individual cells was assessed using an on-chip fluorescence immunoassay with an Alexa Fluor 488-conjugated goat anti-human IgG antibody (Invitrogen, A11013). Cells exhibiting high fluorescence intensity were identified as anti-MDA5 antibody-secreting cells. These cells were individually collected using a glass capillary attached to an automated micromanipulator and transferred into tubes containing cell lysis buffer for subsequent analyses.

#### VH and VL sequencing

Complementary DNA (cDNA) from each single anti-MDA5 antibody-producing cell was synthesized and amplified using SuperScript IV Single Cell/Low-Input cDNA PreAmp Kit (Thermo Fisher Scientific), according to the manufacturer’s instructions. Primary and secondary polymerase chain reaction (PCR) amplifications of the immunoglobulin heavy-chain variable region (VH) and light-chain variable region (VL) were performed using a universal forward primer and gene-specific reverse primers, as previously described^8,9^. Products from the secondary PCR were separated on a 1% agarose gel prepared in TAE buffer. Amplified VH and VL DNA fragments of approximately 600-750 base pairs were excised from the gel and purified using the PCR Clean-Up and Gel Extraction Kit (Takara, 740609). The purified VH and VL DNA fragments were cloned into the pGEM-T Easy vector (Promega, A1360) and subjected to Sanger sequencing.

### Generation of patient-derived F01 and F12 mAbs

For generation of monoclonal anti-MDA5 antibodies F01 and F12, PBMCs were isolated from 3 anti-MDA5-positive patients diagnosed between 1999 and 2021 at Karolinska University Hospital (Stockholm, Sweden), and isolation of MDA5-specific B cells and sequencing of monoclonal BCRs are described separately4.

### Enzyme-linked immunosorbent assay (ELISA)

#### Anti-MDA5 sandwich ELISA for determination of Ab/Fab affinity for MDA5 monomers and filaments

96-well polystyrene high bind microplate for ELISA (Corning) was incubated with or without anti-MDA5 antibodies (1 μg/ml in PBS) overnight at 4°C. The wells were then blocked with blocking buffer (5% BSA in PBS with 0.1% Tween 20, or PBS-T) for 2 h at RT. After blocking, the plates were incubated with FLAG-MDA5(FL):512 bp dsRNA or FLAG-MDA5ΔN:512 bp dsRNA (0.3 nM to 4.0 μM) in blocking buffer for 1.5 h at RT. Plates were then washed with PBS-T three times, then incubated with HRP-conjugated mouse anti-FLAG M2-peroxidase (Sigma-Aldrich) at a dilution of 1:10,000 in blocking buffer for 1 h. Plates were washed with PBS-T three times then developed using TMB (Thermo Fisher Scientific) for 2-3 min. The reaction was stopped by adding 1 M hydrochloric acid. Absorbance at 450 nM was measured using a Biotek M1 Synergy microplate reader within 30 min of stopping the reaction. Signal arising from uncoated wells was subtracted from antibody-coated wells.

#### Direct MDA5 ELISA for validating cryo-EM structures

MDA5ΔN and its variants (2-5 μg/ml) were coated onto 96-well Nunc MaxiSorp ELISA plates (Thermo Fisher Scientific) and incubated for 2 h at RT. The wells were then blocked with blocking buffer (5% BSA in TBS-T) for 1 h at RT. After blocking, the plates were washed three times with TBS-T. The scFv-formatted MDA5 antibodies were added and incubated at RT for 1 h, followed by three washes with TBS-T. For detection, a secondary antibody (HRP-conjugated mouse anti-His-tag; MBL) was diluted 1:5,000 in blocking buffer, added to each well, and incubated for 30 min. The plates were washed three times with TBS-T and then incubated with ELISA POD Substrate Solution (Nacalai Tesque) for 2-3 min. The reaction was stopped by adding 1 N sulfuric acid Stop Solution (Nacalai Tesque). Absorbance at 450 nm was measured using a SpectraMax iD5e plate reader (Molecular Devices) within 30 min of stopping the reaction.

Similarly, MDA5(FL) and its variants (2 μg/ml) were coated onto high bind ELISA microplates (Corning) overnight at 4°C and blocked with blocking buffer (5% BSA in PBS-T) for 1 h at RT. Anti-MDA5 Abs or Fabs were added and incubated at RT for 1 h. Plates were washed with PBS-T three times, then incubated with anti-human IgG-HRP (Promega) at 1:5,000 in blocking buffer for 1 h. After washing plates with PBS-T three times, plates were developed as described above for anti-MDA5 sandwich ELISAs.

#### Direct MDA5 ELISA for patient anti-MDA5 autoantibody detection

High bind ELISA microplates (Corning) were coated with or without MDA5(FL) (2 μg/ml in PBS with 2 mM DTT) overnight at 4°C. The wells were blocked with blocking buffer (5% BSA in PBS-T) for 2 h at RT, then washed with PBS-T three times. The plate was incubated with NP6 mAb 0.128 ng/ml to 10 μg/ml for standard curve) or human plasma diluted at 1:10 to 1:32,000 in plasma dilution buffer [1 mg/ml nuclease-free BSA (Sigma-Aldrich, 126609) in PBS-T] for 1 h. The plate was washed three times, then incubated with anti-human IgG-HRP (Promega) at 1:5,000 in blocking buffer for 1 h. Plates were washed with PBS-T three times and developed as described above for anti-MDA5 sandwich ELISAs.

#### IFN-β and IL-1β ELISAs

For cellular supernatant used in IFN-β and IL-1β ELISAs, cell culture supernatant was first centrifuged at 16,000 x *g* for 2 min at 4°C to exclude cellular debris. For IFN-β ELISA, the LumiKine Xpress hIFN-b 2.0 kit (InvivoGen) was used according to the manufacturer’s instructions; similarly, IL-1β ELISA was performed with the Human Il-1 beta/IL-1F2 ELISA kit Quantikine kit (R&D Systems) per manufacturer’s instructions.

#### TRIM65 sandwich ELISA

High bind ELISA microplates (Corning) were incubated with or without TRIM65^CC-PSpry^ (1 μg/ml in PBS with 2 mM DTT) overnight at 4°C. The wells were blocked with blocking buffer (5% BSA in PBS-T) for 3 h at RT. After blocking, the plates were incubated with MDA5ΔN:512 bp dsRNA (4 to 100 nM for standard curve) or human plasma diluted at 1:50, 1:75, 1:100, 1:150, 1:200 in plasma dilution buffer for 1.5 h at RT. Plates were then washed with PBS-T three times, then incubated with anti-human IgG-HRP (Promega) at 1:5,000 in blocking buffer for 1 h. Plates were washed with PBS-T three times and developed as described above for anti-MDA5 sandwich ELISAs.

### Native gel-shift assay

Assays were performed as previously reported^2^. For demonstration of mAb binding to RLR proteins lacking CARD domains, purified MDA5ΔN, RIG-IΔN, or LGP2 proteins were incubated with Cy5-labeled 112-bp dsRNA for 20 min at RT in binding buffer containing 20 mM HEPES, pH 7.5, 100 mM NaCl, 1.5 mM MgCl₂ and 2 mM ATP. Subsequently, scFv MDA5 antibodies were added, and the mixtures were further incubated at RT for 10 min. Samples were then analyzed on 4–16% Bis-Tris native polyacrylamide gel electrophoresis (PAGE) (Life Technologies). Gels were imaged by detecting Cy5 fluorescence using an iBright FL1500 imaging system (Thermo Fisher Scientific).

For demonstration of mAb binding to full-length MDA5 and RIG-I, purified MDA5(FL) or RIG-I(FL) proteins were incubated with unlabeled 112-bp dsRNA in binding buffer (20 mM HEPES, pH 7.5, 150 mM NaCl, 1.5 mM MgCl_2,_ 2 mM DTT) for 15 min at RT, then with anti-MDA5 Fabs for 5 min at RT. For RIG-I assays, 2 mM ATP was included in the binding buffer. Samples were analyzed on Bis-Tris native PAGE stained with SYBR Gold (Life Technologies). SYBR Gold fluorescence was detected using the ChemiDoc imager (Bio Rad).

For demonstration of differential dsRNA binding by MDA5(FL) WT or gain-of-function mutant, purified MDA5ΔN-WT or -G495R proteins (200-500 nM) were incubated with 112-bp dsRNA (5 ng/μl) for 15 min at RT in the same binding buffer as before. Complexes were analyzed on Bis-Tris native PAGE after staining with SybrGold stain as well.

### Negative-stain electron microscopy

MDA5ΔN-G495R (111 nM) was incubated with 512 bp dsRNA (3.7 nM) in 20 mM HEPES pH 7.5, 100 mM NaCl, 1.5 mM MgCl2, and incubated at RT for 15 min, followed by addition of 1 mM ADP·AlF_4_. ADP·AlF_4_ was prepared by mixing ADP, AlCl_3_, and NaF in a molar ratio of 1:1:3. Antibodies were added (27.8 nM) and incubated at RT for an additional 5 min. For full-length MDA5 filaments, MDA5(FL) (75 nM) was incubated with 512 bp dsRNA (2.5 nM) in the same buffer and conditions as above, followed by ADP·AlF_4_. Antibodies (18.8 nM) were included for an additional 5 min.

Samples were then adsorbed to carbon-coated grids (Electron Microscopy Sciences) and stained with 0.75% uranyl formate as described^10^. Images were collected using the JEM-1400 transmission electron microscope (JEOL) at a magnification of 800x to 30,000x.

### Strep-Tactin pull-down assay

Twin-Strep-tagged or untagged MDA5ΔN (0.5 μM) was mixed with either 112 bp (bait) or 162 bp (prey) double-stranded RNA (dsRNA) at a concentration of 1 ng/μl and incubated at RT for 20 min in binding buffer containing 20 mM HEPES pH 7.5, 150 mM NaCl, and 1.5 mM MgCl₂. The resulting bait filaments were captured using MagStrep Strep-Tactin XT beads (IBA Lifesciences). Beads were washed three times with the same buffer to remove unbound bait filaments, followed by the addition of prey filaments together with either NP6 or NP8 antibodies. Bound dsRNAs were eluted by treatment with Proteinase K (160 U/ml) and analyzed by Bis-Tris native PAGE (Life Technologies). Gels were stained with SybrGold and imaged using an iBright FL1500 imaging system.

### Cryo-EM sample preparation and data collection

For the MDA5ΔN complexes with NP6 and NP8, scFv format of anti-MDA5 antibodies was used. Purified MDA5ΔN-G495R (1 mg/ml) was incubated with the 1012 bp dsRNA (0.050 mg/ml) in 50 mM HEPES, pH 7.5, 100 mM NaCl, 10 mM MgCl_2_ at RT for 20 min. The mixture was then incubated with anti-MDA5 scFv at 1:1.2 molar ratio relative to MDA5ΔN at 4°C until cryo-EM grid preparation. The complex was further incubated in the presence of 2 mM ADP at 4°C for 10 min just before grid preparation. The samples (3 μL) were applied to a freshly glow-discharged Au 300 mesh R1.2/1.3 grid (Quantifoil), in a Vitrobot Mark IV (FEI) at 4°C with a blotting time of 4 seconds under 100% humidity conditions. The grid was plunge-frozen in liquid ethane cooled at liquid nitrogen temperature. The cryo-EM data were collected using a Titan Krios microscope (Thermo Fisher Scientific) installed at the University of Tokyo, running at 300 kV and equipped with a Gatan Quantum-LS Energy Filter (GIF) and a Gatan K3 Summit direct electron detector, operated in the electron counting mode. Movies were recorded at a nominal magnification of 105,000x, corresponding to a calibrated pixel size of 0.83 Å. The electron flux was set to 7.31 e−/pix/sec for 4.83 seconds, resulting in accumulated exposures of 50.0 e−/Å^2^. The data were automatically acquired using the EPU software (Thermo Fisher Scientific), with a defocus range of −0.8 to −2.0 μm, and 3,843 and 2,500 movies were obtained for NP6 and NP8, respectively.

For the MDA5ΔN complexes with F01 and F12, Fab fragments were used. Purified MDA5ΔN-G495R (9 μM) was incubated with 512 bp dsRNA (0.050 mg/ml) in 20 mM HEPES, pH 7.5, 100 mM NaCl, 1.5 mM MgCl_2_ on ice for 1 h. The mixture was then incubated with anti-MDA5 Fabs at 1:1.3 molar ratio relative to MDA5ΔN on ice until cryo-EM grid preparation. F01 complex was then incubated in the presence of 2 mM ADP for a minimum of 20 min and 0.1% glutaraldehyde for a maximum of 10 min on ice. F12 complex was incubated with 2 mM ADP·AlF_x_ for a minimum of 20 min on ice just before grid preparation. The samples were applied to a freshly glow-discharged Au 200 mesh R1.2/1.3 grid in a Vitrobot Mark IV at 4°C with a blotting time of 7 seconds under 100% humidity conditions at the Harvard Cryo-EM Center for Structural Biology. The grid was plunge-frozen in liquid ethane cooled at liquid nitrogen temperature. The cryo-EM data was collected using a Titan Krios microscope (Thermo Fisher Scientific), running at 300 kV and equipped with a Gatan Quantum-LS Energy Filter (GIF) and a Gatan K3 Summit direct electron detector, operated in the electron counting mode. F01 complex dataset was collected at the UMass Chan Medical School Cryo-EM core, and F12 complex dataset at the Cryo-EM Facility at Janelia Research Campus at HHMI Institute. Movies were recorded at a nominal magnification of 105,000x, corresponding to a calibrated pixel size of 0.83 Å. The data were automatically acquired using the EPU software (Thermo Fisher Scientific), with a defocus range of −1.0 to −2.5 μm for F01 and −0.8 to −2.5 for F12. 15,252 and 12,845 movies were obtained for F01 and F12, respectively.

For the MDA5-G495RΔNΔ640-644Δ657-667:dsRNA complexes with NP8 and F12, scFv format of anti-MDA5 antibodies was used. Purified MDA5-G495RΔNΔ640-644Δ657-667 complex (1 mg/ml) was incubated with 1012 bp dsRNA (0.050 mg/ml) in 50 mM HEPES, pH 7.5, 100 mM NaCl, 10 mM MgCl_2_ at RT for 20 min. The mixture was then incubated with anti-MDA5 scFv at 1:1.2 molar ratio relative to MDA5-G495RΔNΔ640-644Δ657-667:dsRNA complex at 4°C until cryo-EM grid preparation. The complex was further incubated in the presence of 2 mM ADP at 4°C for 5 min just before grid preparation. The samples (3 μL) were applied to freshly glow-discharged Au 300-mesh R1.2/1.3 Quantifoil grids and vitrified using a Vitrobot Mark IV (FEI) at 4°C and 100% humidity. The grids were plunge-frozen in liquid ethane cooled at liquid nitrogen temperature. The cryo-EM data were collected using a Titan Krios microscope (Thermo Fisher Scientific) installed at the University of Tokyo, running at 300 kV and equipped with a Gatan Quantum-LS Energy Filter (GIF) and a Gatan K3 Summit direct electron detector, operated in the electron counting mode. Movies were recorded at a nominal magnification of 105,000x, corresponding to a calibrated pixel size of 0.83 Å. The accumulated exposures were 50.0 e−/Å2. The data were automatically acquired using the EPU software (Thermo Fisher Scientific), with a defocus range of −0.8 to −2.0 μm, and 8,375 and 9,104 movies were obtained for NP8 and F12, respectively.

### Cryo-EM data processing and structure refinement

Data processing was performed using the cryoSPARC v4.7.1 software package^11^. The dose-fractionated movies were aligned by Patch motion correction, and the contrast transfer function (CTF) parameters were approximated using Patch-based CTF estimation with the default settings. Particles were automatically picked using Filament Tracer or pre-trained Topaz^12^. The picked particles were curated by 2D classification.

For the NP6 dataset, a selected class containing 826,701 particles after 2D classification was subject into Non-uniform refinement^13^, followed by global and local refinement and reference based motion correction. To deal with the structural heterogeneity of MDA5 filament and scFv NP6, no-align 3D classification was performed using a mask covering the central monomer of MDA5ΔN:NP6:dsRNA duplex. The best class contained 415,377 particles and the resulting 3D map was subjected to Local Refinement, yielding a map with a global resolution of 2.49 Å resolution.

For the filamentous NP6:MDA5:RNA complex, the particle set after 2D classification was further curated by Heterogeneous Refinement using the density of mouse MDA5:dsRNA filament (EMDB 11937) as a reference model. The best class containing 293,099 particles was subjected to Helical Refinement by applying helical symmetry with 76.3 degree twist and 43.5 Å rise, yielding a map at 2.96 Å resolution.

For the F01 dataset, selected classes after 2D classification were curated by ab initio reconstruction with four classes. The best class was subjected to non-uniform refinement^13^, followed by local refinement and 3D classification. The class showing clear density for F01 was selected and subjected to CTF refinement, reference-based motion correction, and local refinement, yielding a map at 3.20 Å resolution.

For the NP8 complex with MDA5ΔN-G495R:dsRNA, selected classes were curated by heterogeneous refinement with six classes. Classes showing the MDA5:RNA filament structure were subjected to non-uniform refinement. A mask was generated for the central MDA5 dimer, and the particles were classified into 10 classes using 3D classification. The best class, containing 257,613 particles was refined using local refinement, yielding a map at 3.07 Å.

For the F12 complex with MDA5ΔN-G495R:dsRNA, selected classes were curated by ab initio reconstruction and heterogeneous refinement. A mask was generated for the central MDA5 dimer, and the particles were classified into 10 classes. The best class, containing 25,448 particles, was refined using local refinement, yielding an improved map at 3.67 Å.

For the NP8 complex with MDA5-G495RΔNΔ640-644Δ657-667:dsRNA, selected particles from the two datasets were curated by heterogeneous refinement and subsequently combined, yielding 1,095,810 particles. A mask was generated for the central MDA5 monomer, and signal subtraction was performed followed by the particles were classified into 12 classes using 3D classification. The best class containing 96,943 particles was refined using local refinement, yielding a map at 2.47 Å.

For the F12 complex with MDA5-G495RΔNΔ640-644Δ657-667:dsRNA, selected classes from one dataset were curated by ab initio reconstruction and heterogeneous refinement, and 2,148,462 particles were subjected to non-uniform refinement. A mask was generated for the central MDA5 monomer, followed by signal subtraction and local refinement, and the particles were classified into 12 classes using 3D classification. The best class containing 195,842 particles was further refined using local refinement, yielding an improved map at 2.09 Å.

### Model building and validation

Initial models were automatically built with ModelAngelo^14^. The manual model building was performed using COOT^15^, with the aid of a density map calculated by DeepEMhancer^16^. Models were refined using real-space refinement in PHENIX^17^, and validated using MolProbity^18^ from the PHENIX package. The cryo-EM density maps were calculated, and molecular graphics figures were prepared using UCSF ChimeraX^19^.

### Differentiation of cells into macrophages

THP-1 cells were differentiated into macrophage-like cells^20^ by seeding at a density of 0.5 x 10^6^/ml in Phorbol 12-Myristate 13-Acetate (PMA, Sigma-Aldrich) 62.5 ng/ml. After 24 h, cells were washed with PBS once and rested in complete media for 24 h. Afterwards, cells were seeded into LPS (1 μg/ml) (Invivogen, tlrl-b5lps) and IFN-γ (20 ng/ml) (GenScript, Z02915) for 24 h, followed by stimulation with ICs.

Primary monocytes were differentiated into macrophages^21^ by first seeding 300 million human PBMCs into a T75 flask overnight. Unattached lymphocytes were aspirated, and cells were washed twice with PBS, then stimulated with 100 ng/ml M-CSF (Proteintech, HZ-1192) for 4 days, with media and supplement change after 2 days. Cells were then treated with 25 ng/ml M-CSF for 2 days for M(M-CSF) cells. For M(LPS/IFN-γ) macrophages, cells were additionally treated with 50 ng/ml LPS and 25 ng/ml IFN-γ for 24 hours.

### Immune complex assembly for cellular stimulation

ICs were formed by incubating 1.8 μM MDA5 protein with 0.06 μM 512 bp dsRNA in RNA binding buffer (20 mM HEPES, pH 7.5, 150 mM NaCl, 1.5 mM MgCl_2_) at RT for 15 min. 1.8 μM antibodies [anti-MDA5 antibodies or human IgG1 lambda isotype control (SouthernBiotech, 0151L-14)] were added and incubated at RT for an additional 5 min. The complexes were kept on ice until they were added directly to culture media for a final concentration of 90 nM MDA5, 3 nM dsRNA, and/or 90 nM mAb, unless stated otherwise.

### RT-qPCR

Total RNAs were extracted using TRI Reagent and Directzol RNA miniprep kit (Zymo Research). cDNA was synthesized using High-Capacity cDNA reverse transcription kit (Applied Biosystems) according to the manufacturer’s instruction. Real-time PCR was performed using a set of gene-specific primers or random primers, a SYBR Green Master Mix, and the StepOne Real-Time PCR Systems (Applied Biosystems). Primer sequences are shown in Supplementary Table 3.

### Cellular FcγR blockade assays

After differentiating THP-1 cells into macrophages, cells were incubated with 10 μg/ml of Fcγ receptor antibodies [Fc Block (BD, 564219), CD64 (Biorad, MCA756G), CD32A (BioXCell, BE00224), CD32B (Cell Signaling, 20312), CD32C (MyBioSource, MBS820241), CD16A (Proteintech, 98293-1-RR), or CD16B (Proteintech, 98192-1-RR)] for 1.5 h. Then cells were stimulated with ICs (13.5 nM MDA5, 0.45 nM dsRNA, and/or 13.5 nM mAb) added directly to the media for 4 hours, followed by harvesting cells for RNA or culture supernatant for cytokine testing. For primary monocyte-derived macrophages, cells were incubated with 91 μg/ml of Fc Block (BD, 564219) for 1 h, then stimulated with ICs (90 nM MDA5, 3 nM dsRNA, and/or 90 nM mAb).

### Immunoblotting

Cells were seeded at 80% confluency in 24- or 48-well plates and stimulated with ICs as described previously. At indicated timepoints, cells were lysed with 1% SDS lysis buffer (1% SDS, 50 mM Tris pH 7.5, 150 mM NaCl, 2 mM DTT) supplemented with 1x Mammalian ProteaseArrest Protease Inhibitor Cocktail (G Biosciences, 786-433), then boiled for 10 min. Supernatant was collected and spun down at 500 x *g* for 5 min, and collected in 1x SDS sample buffer (62.5 mM Tris-HCl pH 6.8, 10% glycerol, 2% SDS, 2.5% β-mercaptoethanol), followed by boiling for 10 min. The following antibodies were used for Western blotting: β-actin (Cell Signaling, 4967), ASC (Cell Signaling, 78000), IFNAR1 (Proteintech, 83002-4), MAVS (Bethyl, A300-782A), MDA5 (NP6, in-house antibody), NALP1/NLRP1 (Cell Signaling, 56719), NLRP3 (Cell Signaling, 15101), Ran (BD Biosciences, 610340), Streptactin-HRP (Bio Rad, 1610381), TLR3 (Proteintech, 83136-3), TRIF (Cell Signaling, 16809), vinculin (Cell Signaling, 4650).

### siRNA electroporation

To knock down transcripts encoding MAVS and TRIF, THP-1 cells were differentiated using PMA 62.5 ng/ml for 24 hours, then electroporated with siRNAs using Neon NxT electroporation system (Thermo Fischer Scientific) according to manufacturer’s instructions. siGENOME SMARTPool siRNAs for *MAVS* and *TICAM1* (Horizon Discovery) were electroporated into cells. Cells recovered for 24 h before being polarized with LPS (1 μg/ml) and IFN-γ (20 ng/ml) for 24 h prior to IC stimulation.

### ASC foci immunofluorescence imaging

Differentiated THP-1 cells were seeded on coverslips and stimulated with ICs as indicated. Cells were fixed with 4% paraformaldehyde (PFA) at RT for 10 min and permeabilized with 0.1% Triton-X at RT for 10 min. Cells were blocked for 1 h at RT with 1% BSA in PBS-T, and stained using ASC (1:100, Adiopogen AL177) overnight. After washing with PBS-T three times, coverslips were incubated with secondary antibody for 1 h. Hoechst 33342 (1:3,000) was used to stain the nuclei. Coverslips were mounted using Fluoromount-G and imaged with a Nikon Ti2 motorized inverted microscope with 60x oil-immersion lens and Hamamatsu Flash4.0 LT camera at the Core for Imaging Technology & Education at Harvard Medical School. Images were taken from 12 fields of view from z-stack images (0.11 μm step size). All images were processed and analyzed with Fiji.

### Fluorescently labeled IC stimulation and IC foci immunofluorescence imaging

Immediately prior to IC assembly, SNAP-MDA5ΔN (5 μM) was labeled with SNAP-Vista Green (NEB) according to the manufacturer’s instructions. ICs were then assembled by incubating 0.9 μM labeled MDA5 protein with 0.03 μM Cy3-labeled 512 bp dsRNA in RNA binding buffer (20 mM HEPES pH 7.5, 150 mM NaCl, 1.5 mM MgCl_2_) at RT for 15 min, then with 0.9 μM Alexa Fluor 647-conjugated NP6 at RT for 5 min. The complexes were added directly to culture media of differentiated THP-1 cells seeded onto coverslips. After 2 hours, cells were fixed with 4% PFA at RT for 10 min, washed with PBS, then stained with Hoeschst 33342 (1:3,000). As above, coverslips were mounted and imaged with a Nikon Ti2 inverted microscope with 60x oil-immersion lens. Images were taken from 15-20 fields of view, and images were processed and analyzed with Fiji.

### *In vitro* neutrophil extracellular trap (NET) formation assay

Neutrophils were seeded at a density of 0.5 x 10⁵ cells per well in HEPES-buffered, phenol red-free RPMI 1640 medium in poly-L-Lysine coated chambered coverslips (μ-Slide 18 Well-Glass Bottom, Ibidi). Cells were allowed to adhere for 30 minutes at 37°C and 5% CO₂. Following adherence, neutrophils were stimulated with either vehicle control, 4 μM ionomycin (Invivogen), or 100nM PMA (Invivogen) for 4 hours at 37°C and 5% CO₂. After stimulation, cells were fixed with 4% paraformaldehyde in PBS for 10 minutes at RT, followed by three washes with PBS.

For immunofluorescence staining, fixed cells were permeabilized with 0.1% Triton X-100 in PBS for 10 minutes and subsequently blocked with 3% BSA in PBS containing 0.5% Tween-20 for 1 h at RT. Primary antibodies against myeloperoxidase (MPO; R&D Systems, AF3667, 1:200) and citrullinated histone H3 (H3Cit; Abcam, ab281584, 1:500) were applied and incubated overnight at 4°C. Following primary antibody incubation, cells were washed three times with PBS and incubated with fluorophore-conjugated secondary antibodies (Abcam, 1:2000) for 2 h at RT in the dark. Nuclear DNA was counterstained with Hoechst 33342 (1:10,000) for 20 minutes at RT. After final washes with PBS, samples were stored in PBS at 4°C prior to imaging. Fluorescence microscopy was performed using a Keyence BZ-X800 all-in-one fluorescence microscope with a 20X objective. Images were acquired from two randomized, non-overlapping fields of view per well. For NETosis analysis, cells were counted based on Hoechst staining, and NETs were considered extracellular DNA strands or smears that can co-localize with MPO or H3cit. H3cit-positive NETs were counted only if they overlaid with Hoechst staining.

### Plasma and PBMC RNA-seq library preparation

Cell-free RNAs (cfRNAs) were extracted from plasma using the Plasma/Serum RNA Purification kit (Norgen). To account for fluctuations in RNA isolation and variable RNA content across different patient samples, 4 μl of ERCC (at 1:10,000 dilution, Invitrogen) was added to 2 ml plasma prior to RNA isolation. Total RNA was extracted from human PBMCs using RNeasy mini kit (Qiagen). 4 ul of ERCC (at 1:10,000) was added to 20 ng of PBMC RNA. The concentration of extracted RNAs was assessed using the Qubit RNA assay (Life Technologies) and High Sensitivity RNA ScreenTape (Agilent Technologies). For both plasma- and PBMC-derived RNAs mixed with ERCC, 2 ng of input was used for cDNA library preparation using the SMARTer Stranded Total RNA-Seq Kit v3-Pico Input Mammalian (Takara Bio) according to the manufacturer’s instructions. cDNA libraries were sequenced using the Illumina partial lane NovaSeq platform.

For preparation of cDNA libraries from PBMC RNA using the Collibri Stranded RNA Library Prep Kit (Thermo Fisher Scientific), 250 ng of total RNA was combined with 1 μl of ERCC (at 1:200) for rRNA depletion using depletion beads and further purified using Zymo-Spin IC columns (Zymo Research). RNA was fragmented by heat at 95°C for 6 min, and libraries were prepared according to the manufacturer’s instructions. cDNA libraries were sequenced using the Illumina partial lane NovaSeq platform.

### TRIM65-pulldown RNA-seq

Pull-downs were performed at 4°C unless specified. Strep-HA-TRIM65^CC-PSpry^ was captured onto anti-HA magnetic beads (Pierce) in SEC buffer (50 mM HEPES, pH 7.5, 150 mM NaCl, 0.1% Tween 20) for 45 min. TRIM65-bound beads were washed twice with SEC buffer and then blocked with 0.1% BSA in SEC buffer for 1 h until ready to mix with diluted plasma. Plasma was diluted 1:5 in plasma dilution buffer (1 mg/ml nuclease-free BSA in PBS-T) with 0.8 U/μL RNase inhibitor (NEB, M0307L) and incubated with TRIM65-bound beads rotating overnight. Beads were then washed three times with PBS-T with 40 U/ml RNase inhibitor. RNA was eluted with TRI Reagent rotating at RT for 1 h. RNA was extracted using Zymo-Spin II columns (Zymo Research), followed by DNase digestion (NEB, M0303) at 37°C for 15 min and further clean up using Zymo-Spin IC columns. RNA was directly used for cDNA library preparation using the SMARTer Stranded Total RNA-Seq Kit v3-Pico Input Mammalian according to the manufacturer’s instructions. cDNA libraries were sequenced using the Illumina partial lane NovaSeq platform.

### RNA-seq analysis

#### Mapping and differential gene expression analysis

Pair-ended reads were trimmed by Trimmomatic v0.39^22^, and mapped to Grch38.p14 by STAR aligner v2.7.9a^23^ in splice-aware mode. Resultant bam files were sorted with Samtools v1.21^24^ by read coordinates. Sorted bam files were processed by Rsubread^25^, and reads were counted for human gene transcripts by Gencode v43 (based on CDR and UTR counts)^26^. Multi-mapping reads were counted by fraction. Read count files were processed in Deseq2^27^, and Volcano plots were generated by EnhancedVolcano v1.20.0 with p-values and Log2FC values from Deseq2. For pathway analysis, genes were selected based on indicated criteria (see figure legends) and analyzed using EnrichR^28^ with Reactome 2024 database^29^. The -log10 of pathway FDR values were used for ranking and plotting pathway enrichment graphs. For patient-derived plasma and PBMC RNA-seq, ERCC spike-in control was used for normalization. ERCC was mapped to Invitrogen ERCC92 reference, and gene expression and area under the curve (AuC) values were normalized by a loess function^30^.

To generate differential gene expression heatmaps for THP-1 cells, sample stimulated with MDA5 filament ICs (with either NP6 or NP8) were combined as one group, and mock groups were used as controls for Deseq2 inputs. Genes with *p-adj*<0.5 and Log2FC>0 were selected as inputs for Reactome pathways. Genes identified by top Reactome pathways were separated into 3 groups: Cell death/necrosis pathways (shown in Figure 3a), IFN/ISG pathways, and other immune-related genes (shown in Figure 2b). *Z*-score heatmaps were calculated by the number of standard deviations above or below the mean value of genes for different groups.

#### LINE/SINE/ERV analysis

Normalized AuC values of all LINE, SINE and ERV regions in Hg38 by Repbase was calculated for patient and HC samples. For each independent LINE/SINE/ERV region, the AuC value of a given patient and an averaged value of HC1/2 were compared, and only the repeat regions with patient over HC Log2FC < −1 or >1 were considered (to be referred as “differential regions”). The sum of AuC values from all differential regions were calculated for each repeat class, and the overall Log2FC of all samples over the mean of HC were used to generate the heatmap.

For gene feature analysis of LINE RNA in Figure 6a and Extended Data Figure 9a, only uniquely mappable reads (MAPQ>10)^31^ were used by filtering Bam files. Each read uniquely mappable to a LINE locus was assigned to one of the 7 groups (CDR, 5’UTR, 3’UTR, intron, <3 kb upstream, <3 kb downstream, intergenic). To account for the LINE orientation relative to genes, each LINE region within 3 kb of genes was further divided into “sense potential” or “antisense potential”, and was separately counted in gene feature analysis.

For scatter plots of LINE RNA in Figure 6b and related figures, mean normalized read counts of each LINE region was calculated for each patient or HC1/2 combined. As with above, only uniquely mappable reads (MAPQ>10)^31^ were analyzed. LINE regions within 3 kb of a gene were labelled as gene-associated LINEs, and further sub-grouped by whether the associated gene is up-regulated, down-regulated or without change in patients over HC TPM value, with a cutoff of Log2FC>0.3, Log2FC<-0.3, and |Log2FC|≤0.3. The top 100 up-regulated genes associated with the most abundant LINE RNA were used as inputs for Reactome pathway discovery by EnrichR.

#### Quantification of A-to-I editing

Bam files were filtered for MAPQ>10 for unique mapping. Reads containing mismatches relative to Hg38 were selected using bcftools and filtered for regions within Repbase-annotated LINEs. The number of uniquely mapped reads with A-to-G mismatches and other types of mismatches were then counted and normalized to per-million total primarily mappable reads by sample. To eliminate the reads containing true genomic single-nucleotide polymorphisms, mismatches that are consistently present in all reads were eliminated from the analysis.

#### Analysis of LINE by mapping to Repbase data

We first compared LINE sequences among family members in Extended Data Figure 7d. The Repbase consensus of all L1 repeats were multiple aligned by MAFFT algorithm^32^ and sorted by percentage identity values to L1HS. The sorted L1 sequences were used to generate a similarity matrix heatmap by percentage identity values. To analyze sense and antisense LINE transcripts categorized into individual LINE subfamily in Figure 5e and Extended Data Figure 8a, reads were mapped to Repbase in a strand-specific manner and counted and normalized against ERCC.

#### TEtranscripts

TEtranscripts (v2.2.4)^33^ and GRCh38_GENCODE_rmsk_TE.gtf annotation file (April 2026) were used to quantify transposable element (TE) expression from BAM files generated by allowing up to 100 alignments per read to retain multimapping reads. TE counts were extracted from the resulting *gene_TE_analysis.txt files by selecting features containing “:” in the first column, as recommended in the TEtranscripts GitHub repository. For heatmap visualization, TEs with baseMean > 1000 in PBMC RNA-seq were retained to exclude lowly expressed elements, and to use the consistent TE set across all heatmaps. Applying the same expression filter to plasma yielded a similar TE set. Because fold-change estimates can be inherently unstable and prone to artificial inflation at low abundance, TEtranscripts output was further analyzed using negative-binomial generalized linear modeling (GLM) and fold-change shrinkage estimation, as in previous studies^34–38^.

#### Virome analysis

A virome reference was generated from all curated RefSeq viral genomes (n = 18,670; February 2025). Reads were first mapped to the human genome (GRCh38), and unmapped reads were re-mapped to the virome reference with multi-mapping allowed. Mapping read counts did not support enhanced viral presence in anti-MDA5-DM patients over HC1/2. Different mappers (star/v2.7.9a and bowtie2/v2.5.4) were also tested for this process with similar conclusions.

## Data availability

The atomic coordinates have been deposited in the Protein Data Bank with accession codes PDB: 21SJ, 22HM, 28BD, and 28BE. The cryo-EM maps included in this study have been deposited in the Electron Microscopy Data Bank under the accession codes EMD-67964, EMD-68271, EMD-81614, EMD-81615, EMD-68272, and EMD68273. The raw sequencing and read count data were deposited in GSE315387. The scripts used in this manuscript were documented in this Github link: (https://github.com/DylannnWX/Mda5_dermatomyositis_manuscript)

## Extended Data Figures 1-9

**Extended Data Figure 1.**
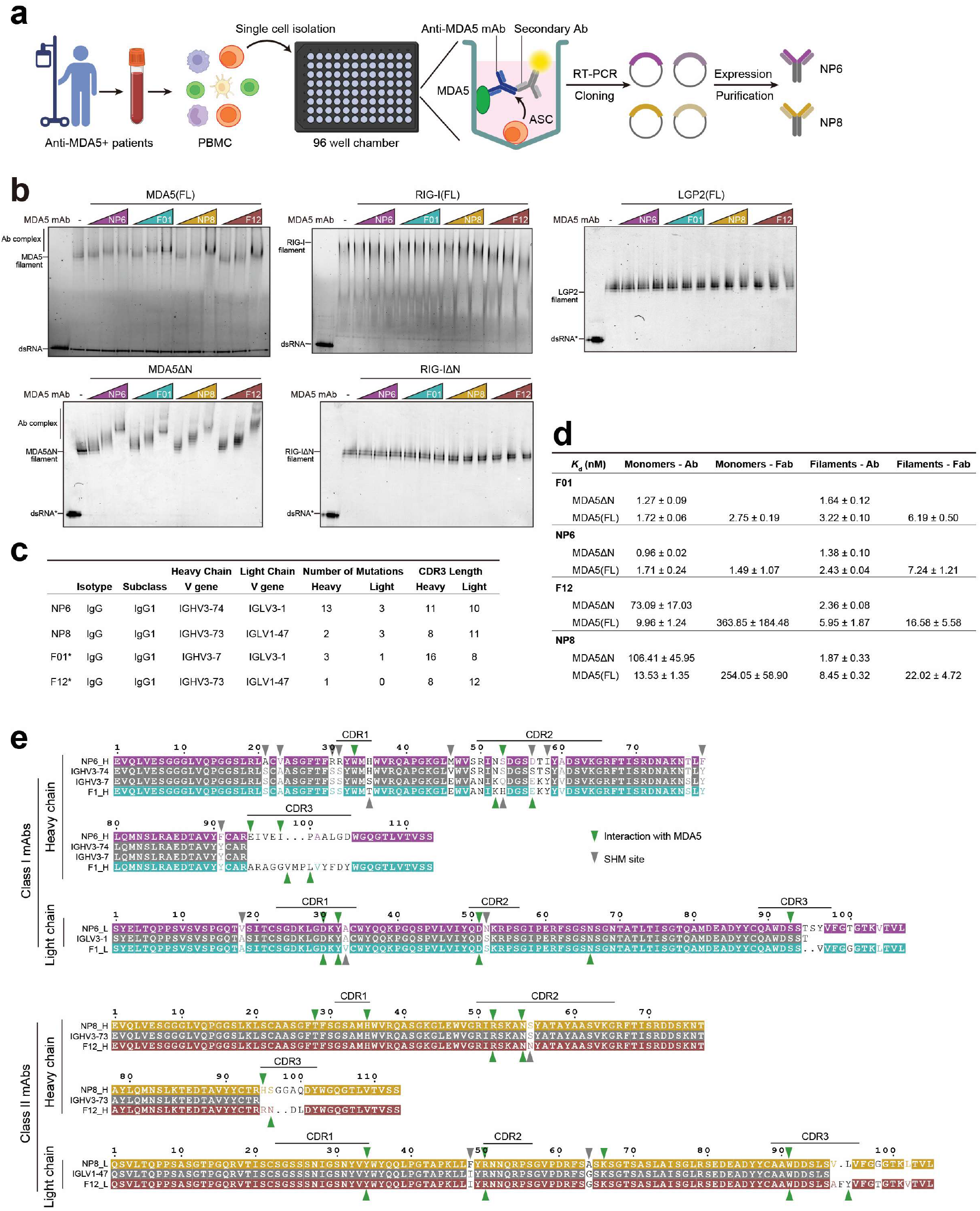
B-cell cloning and sequences of monoclonal MDA5 antibodies. **a**) Schematic overview of the antibody cloning process of NP6 and NP8 from an anti-MDA5 DM patient. See Methods. **b**) Native gel mobility shift assay showing specificity of anti-MDA5 mAbs. RLR filaments were formed by mixing MDA5, RIG-I, or LGP2 proteins (full-length, FL; or CARD-deletion, ΔN) with 112 bp dsRNA and incubating with anti-MDA5 scFv (used with MDA5ΔN, RIG-IΔN, LGP2), or anti-MDA5 Fabs [used with MDA5(FL), RIG-I(FL)]. Anti-MDA5 scFv and Fabs bind specifically to MDA5ΔN and MDA5(FL) filaments but not RIG-I or LGP2. **c**) Summary of mAb gene usage. *F01 and F12 (from Ref 24) are displayed for ease of comparison. **d**) Summary of FLAG-MDA5 affinity for monoclonal Abs (IgG) and Fabs. MDA5 (ΔN or FL) in the monomeric and filamentous forms were compared. Apparent *K*_d_s were determined by ELISA binding curves. See Figure 1b for representative curves of FLAG-MDA5 monomer and filament affinities for Fabs. Data is shown as mean ± SD of 2 biological replicates. **e**) Sequence alignment of class I and class II anti-MDA5 antibodies. Residues interacting with MDA5 are indicated by green triangles. Gray triangles indicate somatic hypermutation (SHM) sites. Sequences were aligned using Clustal Omega, and alignment figures were generated using ESPript3.

**Extended Data Figure 2.**
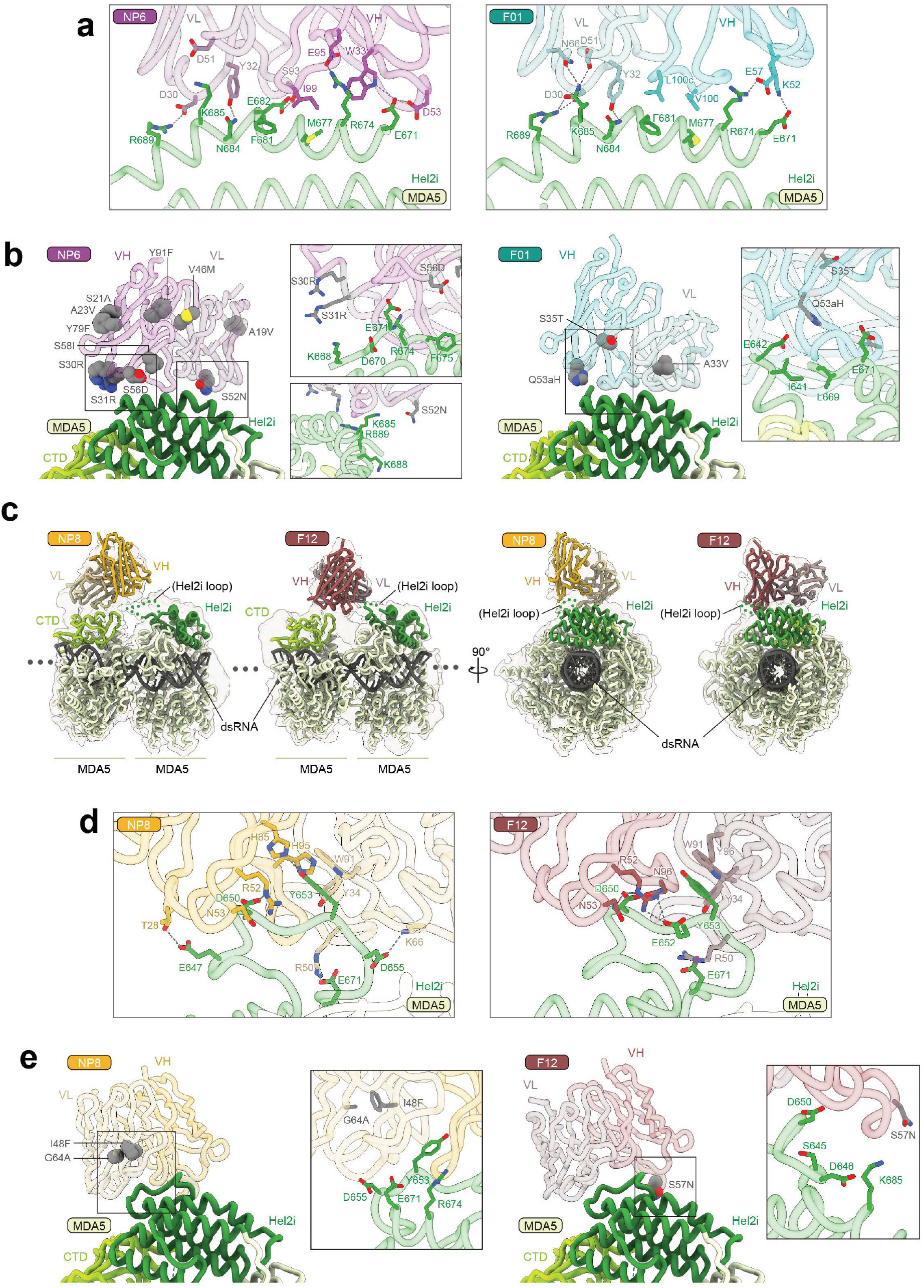
Structural characterization of MDA5-mAb interactions. **a)** Close-up view of the interaction between Hel2i domain of MDA5 and NP6 (left) or F01 (right). The residues involved in the interactions are shown as stick model. **b** Mapping of somatic hypermutations (gray spheres) on NP6 (left) and F01 (right). Close-up views of boxed areas show that these mutated residues do not directly contact MDA5. **c** Cryo-EM structures of MDA5ΔN in complex with class II mAbs. The cryo-EM map was low-pass filtered to 5 Å resolution and is shown at 10% transparency. Hel2i and CTD subdomains are colored dark green and yellow green, respectively. NP8 and F12 are colored yellow and red, respectively. Because of the low local resolution for the mAbs, we docked AlphaFold-predicted antibody models together with known MDA5 structure (PDB ID 7JL0)^21^ into the density. Class II mAbs appeared to contact two adjacent protomers within the filament, explaining their preference for filaments. One interaction, associated with stronger density, involves the MDA5 C-terminal domain (CTD), whereas the second interaction involves Hel2i of the neighboring protomer, near a likely disordered loop (dashed green line, residue 645-662). Deletion of the Hel2i loop abolished both NP8 and F12 binding to monomeric MDA5, whereas CTD deletion had minimal impact (see Extended Data Figure 3a), indicating that Hel2i provides the primary binding interface. The intrinsically disordered nature of this Hel2i loop likely resulted in positional heterogeneity of class II mAbs, accounting for their weaker density and lower local resolution. **d** Cryo-EM structures of MDA5ΔNΔ640–644Δ657–667 in complex with NP8 (left) and F12 (right), with close-up view of the interaction between Hel2i loop of MDA5 and mAbs. The residues involved in the interactions are shown as stick model. **e** Mapping of somatic hypermutations (gray spheres) on NP8 (left) and F12 (right). Close-up views of boxed areas show that these mutated residues do not directly contact MDA5.

**Extended Data Figure 3.**
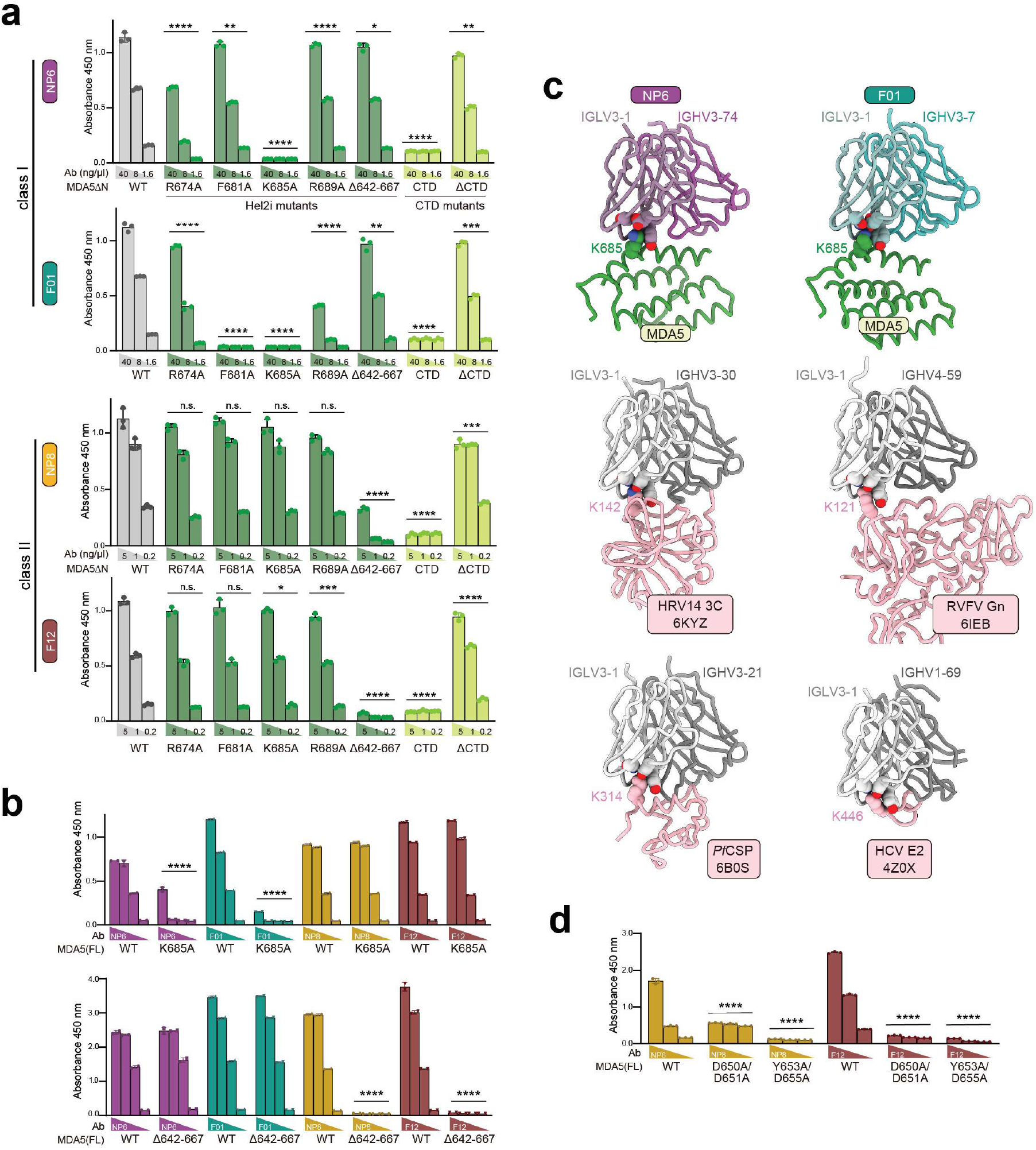
Biochemical validation of MDA5–mAb interactions. **a**) Mutational analyses of MDA5-mAb interactions by ELISA. Monomeric MDA5ΔN (WT or variants) was immobilized on ELISA plates and incubated with His-tagged scFv antibodies at three different concentrations. Antibodies bound to MDA5 were detected using an HRP-conjugated anti-His antibody. Data are presented as mean ± SD of 3 technical replicates. *P*-values were calculated by two-way ANOVA to compare antibody binding to WT versus mutant MDA5 across 3 Ab concentrations. **b, d)** Mutational analyses of MDA5-mAb interactions by ELISA. Monomeric MDA5(FL) (WT or variants) was immobilized and incubated with (b) mAbs (0.6 ng/ml - 10 μg/ml) or (d) Fabs (0.2 - 5 μg/ml), then detected using HRP-conjugated anti-human IgG. Data are represented as mean ± SD of 2 technical replicates. *P*-values were calculated by two-way ANOVA. **c** Conserved antigen-antibody interactions involving the IGLV3-1 light chain. Published structures are aligned to the light chain of NP6. The GRAB motif (Y32, D51, and N66) recognizes a conserved lysine residue. Data are representative of 2 independent experiments. \**P* < 0.05, \*\**P* < 0.01, \*\*\**P* < 0.001, \*\*\*\**P* < 0.0001. *n.s*, not significant.

**Extended Data Figure 4.**
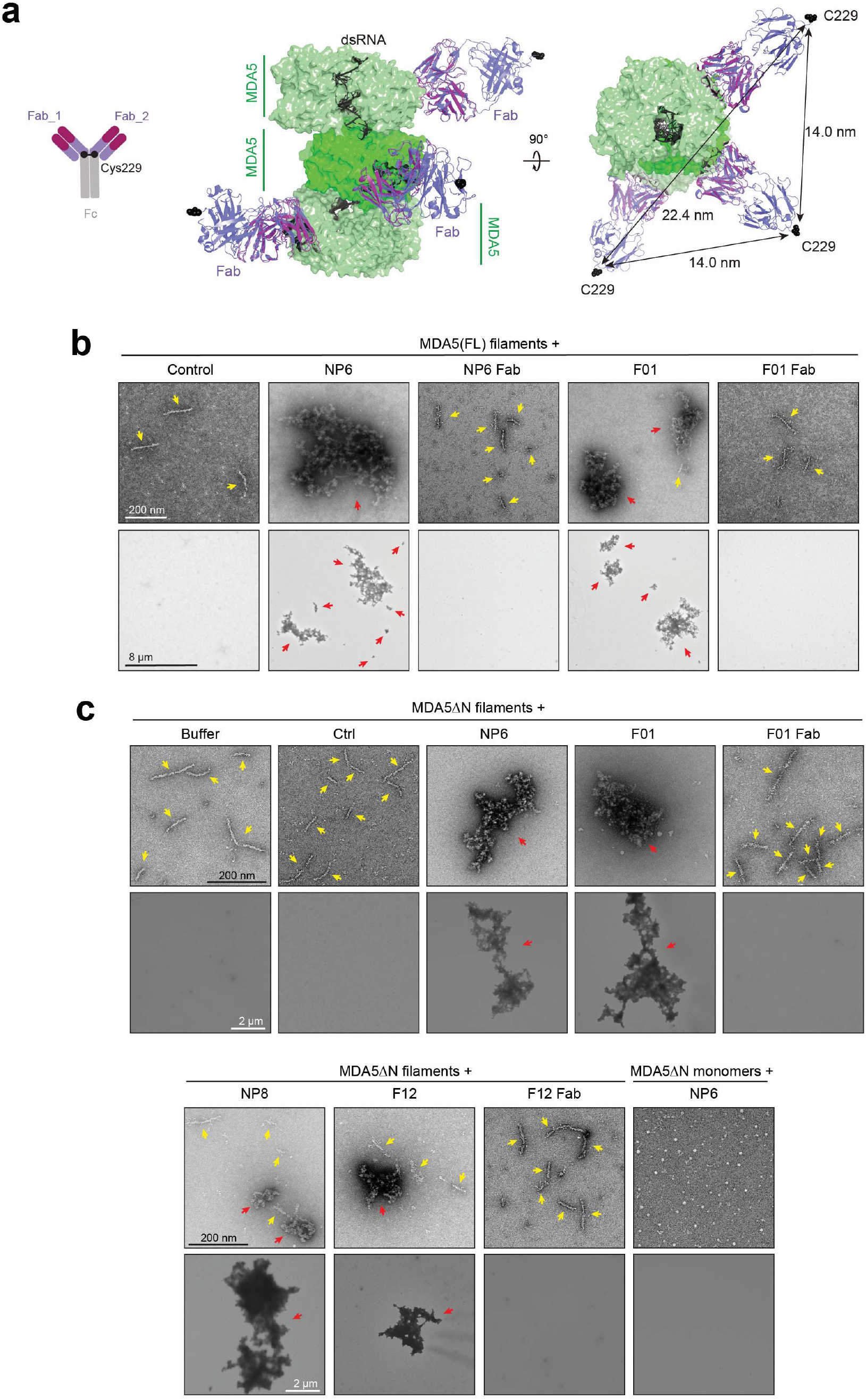
Antibody-mediated cross-bridging of MDA5 filaments. **a**) Individual Fabs from the crystal structure of IgG1 (PDB: 1IGY, slate color) were modeled into the MDA5-NP6 complex structure (MDA5 in green, NP6 in magenta) to predict the approximate positions of the CH1/hinge junction, specifically Cys229 (black dots), which forms the interchain disulfide bond that tethers the two heavy chains together in an intact IgG1. Between two Fabs bound to adjacent MDA5 subunits or subunits two positions apart in the filament, the equivalent Cys229 residues are ∼14 nm and ∼22 nm apart, respectively. However, if both Fabs belonged to the same antibody, their Cys229 residues would need to be within disulfide bond distance (∼2 Å) of each other. This shows that bivalent engagement of two MDA5 subunits within the same filament by a single antibody is geometrically impossible. **b** Negative stain images of MDA5 filaments formed by wild-type MDA5(FL) (75 nM) ± 512 bp dsRNA (2.5 nM) bound by antibodies (18.8 nM), imaged at 30,000x (top row) and 800x magnification (bottom row). Yellow arrows indicate individual filaments, whereas red arrows indicate filament aggregates. Upper left 2 images are also seen in Figure 1h. **c** (Top row) Negative stain images of MDA5 filaments formed by MDA5ΔN (G495R) (111 nM) ± 512 bp dsRNA (3.7 nM) bound by antibodies (27.8 nM), imaged at 25,000x magnification. (Bottom row) Filaments were formed with 2.7-fold higher protein and RNA concentrations and imaged at 2,000x. Arrows are colored as in (b).

**Extended Data Figure 5.**
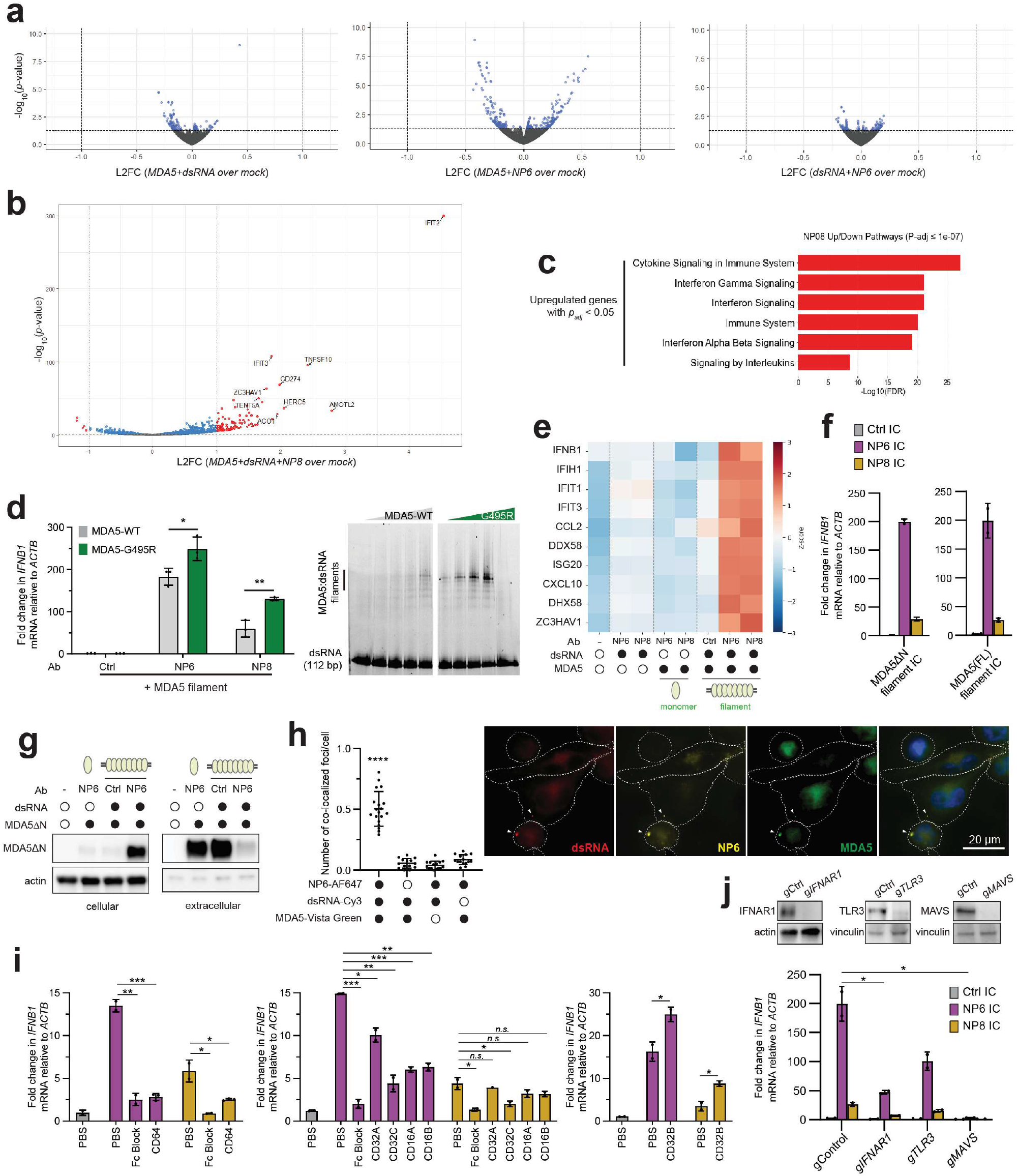
MDA5 filament ICs activate inflammatory pathways in macrophages. **a**) Volcano plots of THP-1-derived macrophages stimulated with MDA5ΔN filaments (90 nM; dsRNA 3 nM; left), MDA5 monomers (90 nM) with NP6 (90 nM) (middle), or dsRNA (3 nM) with NP6 (90 nM), relative to mock. Genes with log_2_-fold change |L2FC| >1 and *p* <0.05 (red) are shown; all other genes shown have *p* > 0.05 (gray) or *p* < 0.05 (blue). **b** Volcano plot of differentially expressed genes in THP-1-derived macrophages stimulated with MDA5ΔN filament-NP8 ICs over mock. Each point is colored as in Figure 2a. **c** Reactome pathways of significantly upregulated genes in THP-1-derived macrophages stimulated with NP8-MDA5ΔN filament ICs over mock (*p_adj_* < 0.05, L2FC > 0). **d** (Left) *IFNB1* mRNA levels in THP-1-derived macrophages stimulated with MDA5ΔN filaments formed by WT or gain-of-function G495R mutant (90 nM) and 512 bp dsRNA (3 nM), in complex with isotype control, NP6, or NP8 antibody (90 nM). Data was normalized to mock treatment. *P*-values were calculated by two-tailed Student’s *t* test. (Right) Native gel mobility shift assay to monitor dsRNA binding capacity by MDA5. Filaments of MDA5ΔN-WT or -G495R (200-500 nM) were formed on 112 bp dsRNA (69 nM). **e** Heatmap of *Z*-scored expression of interferon and interferon-stimulated genes in healthy human monocyte-derived macrophages stimulated with combinations of MDA5ΔN (98.2 nM monomer), 512 bp dsRNA (3.3 nM), mAb (NP6/NP8, 98.2 nM). **f** *IFNB1* mRNA levels in THP-1-derived macrophages stimulated with (left) dsRNA-bound MDA5ΔN filaments (90 nM, dsRNA 3 nM) or (right) dsRNA-bound MDA5(FL) filaments (90 nM, dsRNA 3 nM) with isotype control, NP6, or NP8 antibodies (90 nM). **g** Cellular and extracellular MDA5ΔN protein levels in THP-1-derived macrophages stimulated with MDA5 monomer or filament ICs as in (a). 8 hours post-stimulation, supernatant and washed cell pellet were analyzed for MDA5ΔN, which was added extracellularly. **h** Immunofluorescence of THP-1-derived macrophages stimulated with combinations of Alexa Fluor 647-conjugated NP6, Cy3-labeled 512 bp dsRNA, and/or SNAP-Vista Green-labeled SNAP-MDA5ΔN. Nuclei were stained with Hoechst 3342. (Left) Quantitation of the number of co-localizing intracellular foci per cell within a field of view. (Right) Representative image with colocalizing intracellular foci indicated by white arrowheads. Cell boundaries were determined by brightfield images. **i** *IFNB1* mRNA levels in THP-1-derived macrophages pre-treated with PBS or FcγR antibodies (10 μg/ml) for 1.5 hours, then stimulated with dsRNA-bound MDA5ΔN filaments (13.5 nM; 512 bp dsRNA, 0.45 nM) and isotype control, NP6, or NP8 antibodies (13.5 nM). Data was normalized to mock treatment. *P*-values were calculated by two-tailed Student’s *t* test. **j** *IFNB1* mRNA levels in THP-1-derived macrophages stably transduced with gRNAs targeting the indicated genes or a non-targeting control stimulated with dsRNA-bound MDA5(FL) filaments (90 nM, dsRNA 3 nM) with isotype control, NP6, or NP8 antibodies (90 nM) (bottom). (Top) Western blot showing knockdown of IFNAR1, TLR3, and MAVS expression. *P*-values were calculated by two-tailed Student’s *t* test. Data are presented as mean ± SD of at least 2 biological replicates. RNA-seq results contain 2 biological repeats. All other data are representative of at least 3 independent experiments. \**P* < 0.05, \*\**P* < 0.01, \*\*\**P* < 0.001, \*\*\*\**P* < 0.0001, *n.s*., not significant.

**Extended Data Figure 6.**
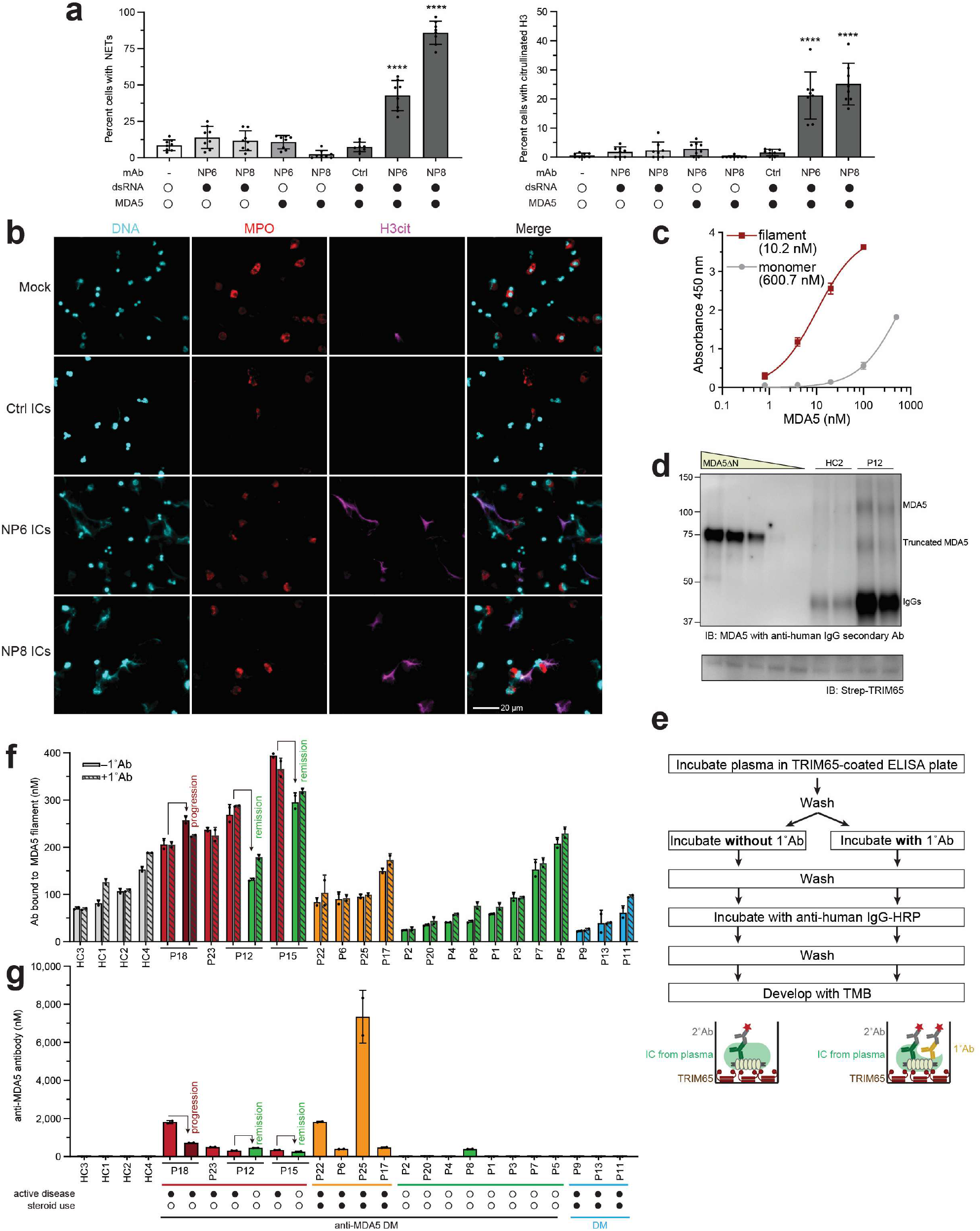
Extracellular MDA5 filament ICs are present in anti-MDA5 DM and activate human neutrophils. **a**) Quantitation of NETosis (left) and histone 3 citrullination (right) frequency in healthy human neutrophils stimulated with combinations of 512 bp dsRNA (3 nM), MDA5ΔN (90 nM), and/or isotype control or NP6/8 mAbs (20 nM). Four fields of view per well were analyzed from two biological replicates. Data are presented as mean ± SD. *P*-values were calculated by one-way ANOVA comparing relevant conditions for NP6 or NP8 ICs. \*\*\*\**P* < 0.0001. **b** Immunofluorescence of DNA (Hoechst 3342, cyan), myeloperoxidase (MPO, red), citrullinated histone 3 (H3cit, magenta) in healthy human neutrophils stimulated with MDA5ΔN filament ICs (MDA5ΔN, 90 nM; 512 bp dsRNA, 3 nM; mAb, 20nM). This experiment was repeated using neutrophils from three different healthy donors, which showed similar patterns (see Figure 3e). **c** Binding curves of TRIM65 to MDA5ΔN monomers or filaments, measured by sandwich ELISA, with apparent *K_d_* shown in parentheses. Filaments were assembled from MDA5ΔN (5 μM) and 512 bp dsRNA (160 nM). Bound filaments were detected with F01 mAb (1°) and anti-human IgG-HRP (2°). **d** Immunoblot of TRIM65 sandwich ELISA, showing TRIM65-captured MDA5 isolated from HC2 and P12 plasma replicate samples. As a control, TRIM65-captured, *in vitro*-reconstituted MDA5ΔN filaments are shown in lanes 1-5. Top blot was probed with NP6 primary antibody followed by anti-human IgG secondary antibody, showing two upper bands with sizes that correspond to full-length MDA5 (∼120 kDa) and MDA5ΔN (∼85 kDa). The lower band corresponds to the heavy chain of denatured IgGs (∼50 kDa), which is detected by the secondary antibody. Bottom blot was probed with HRP-conjugated streptactin. **e** Workflow of TRIM65 sandwich ELISA in which TRIM65-captured MDA5ΔN filaments were incubated without NP8 to measure endogenous antibodies co-purified with MDA5 filament. This assay was repeated with the addition of 1° Ab (NP8), which would increase signal if additional antibody-binding sites on TRIM65-captured ICs remained available. **f** Plasma concentrations of antibodies bound to MDA5 filaments, determined by TRIM65 sandwich ELISA without or with NP8 probing. Addition of NP8 had minimal impact on signal, indicating that MDA5 filaments are largely saturated with endogenous antibodies. Antibody concentrations were interpolated from a standard curve generated using TRIM65-captured MDA5ΔN filaments (20 μM) assembled on 512 bp dsRNA (640 nM) and probed with NP8 (1°Ab) followed by anti-human IgG-HRP (2°Ab). Data are presented as mean ± SD of 2 technical replicates for 4 dilutions. **g** Plasma anti-MDA5 antibody concentrations determined by direct ELISA using MDA5(FL)-coated plates, calibrated using an NP6 standard curve. Data are presented as mean ± SD of 2 technical replicates for 4 dilutions. Active disease at the time of blood draw was defined by the treating physician and confirmed by a reviewing rheumatologist.

**Extended Data Figure 7.**
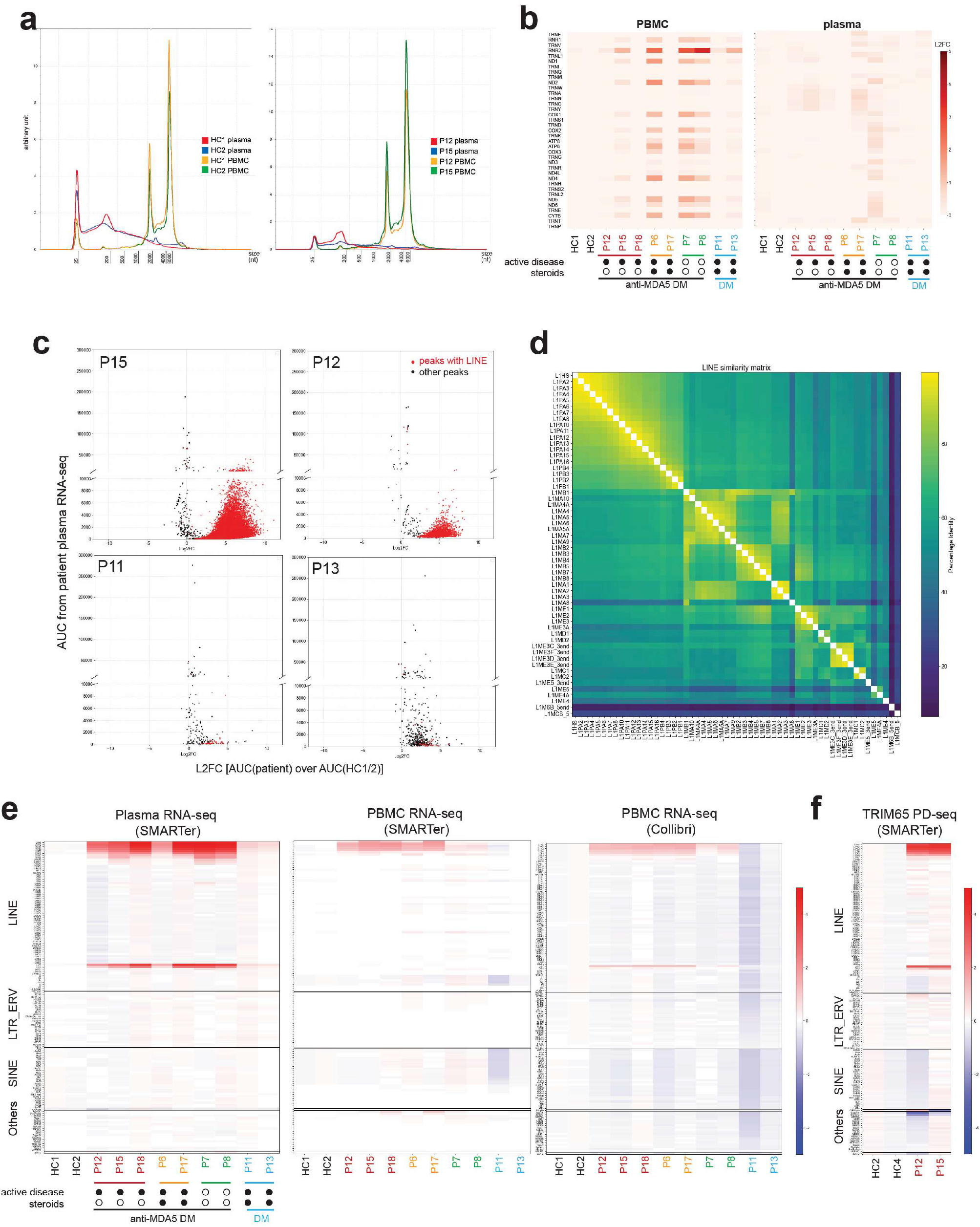
LINE retroelements are enriched in anti-MDA5 DM plasma and PBMC RNA. **a**) Electropherogram of RNA extracted from plasma and PBMCs of healthy controls (left) and anti-MDA5 DM patients (right). **b** Heatmap of log_2_-fold change (L2FC) in mitochondrial RNAs in PBMCs (left) and plasma (right) of healthy controls and DM patients. **c** Whole genome peak calling and annotation of plasma RNA-seq from anti-MDA5 DM patients (P15, P12; top) and non-MDA5 DM patients (P11, P13; bottom). Peaks were called using HC1/2 plasma RNA-seq as the control. Each dot represents an individual peak, plotted by peak area under the curve (AUC) in patient samples (y-axis) versus L2FC in patients relative to healthy controls (x-axis). Peaks containing one or more LINE elements were colored red to distinguish from other peaks (black). **d** Similarity matrix of all LINE sequences based on Repbase, sorted by percentage identity value to L1HS. **e** TEtranscripts heatmaps of plasma (left) and PBMC (center and right) samples. PBMC RNA-seq was performed using two independent library preparation methods (SMARTer and Collibri) to control for potential methodological bias. Only transposable elements with moderate-to-high expression in SMARTer PBMC RNA-seq are shown and are grouped by repeat family. L2FC values were calculated by comparing each patient sample with average of HC1 and HC2. **f** TEtranscripts heatmap of TRIM65 pull-down (PD) RNA-seq, as in (e), with L2FC relative to average of HC2 and HC4.

**Extended Data Figure 8.**
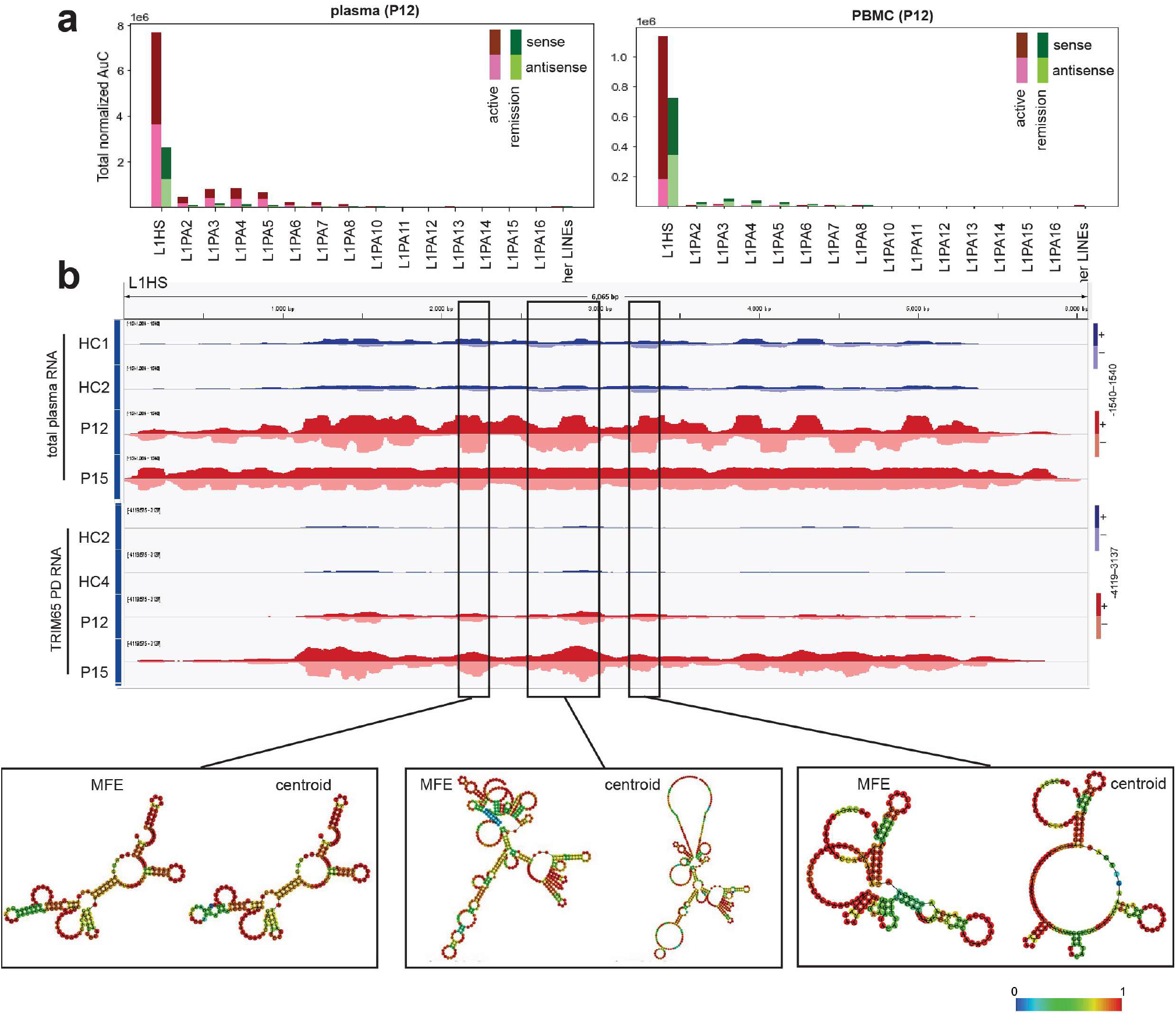
Extracellular LINE-1 RNAs in anti-MDA5 DM may exist as sense:antisense duplexes. **a**) Levels of LINE-1 (L1) and all other LINE elements in plasma (left) and PBMCs (right) from individual P12 during active (red) and quiescent (green) disease. Sense (dark) and antisense (light) sequences are shown. Both sense and antisense LINE-1 RNAs are highly enriched in plasma relative to PBMC, but antisense RNA showed preferential enrichment, resulting in a balanced sense-to-antisense ratio in plasma. This pattern is consistent with the selective stabilization of sense:antisense duplexes and the preferential degradation of unpaired and excess sense RNA in nuclease-rich plasma. **b** (Top) Genome-browser views of total plasma RNA-seq and TRIM65 pull-down (PD) RNA-seq, with positive- and negative-strand reads shown as mirrored tracks. Sense and antisense LINE-1 RNAs exhibit balanced representation in both plasma total RNA and TRIM65 pulled-down RNA. This is consistent with sense:antisense duplex formation and association with MDA5 filaments in plasma. (Bottom) RNAfold secondary structure prediction in representative L1HS regions enriched by TRIM65 PD. (Bottom) Minimum-free-energy (MFE) and centroid structure predictions using sense RNA sequences, suggesting little intrinsic potential to form >100 bp duplex required for MDA5 binding. Therefore, MDA5 association is unlikely to arise from intramolecular secondary structures. Nucleotide colors indicate base-pairing probabilities derived from the thermodynamic ensemble.

**Extended Data Figure 9.**
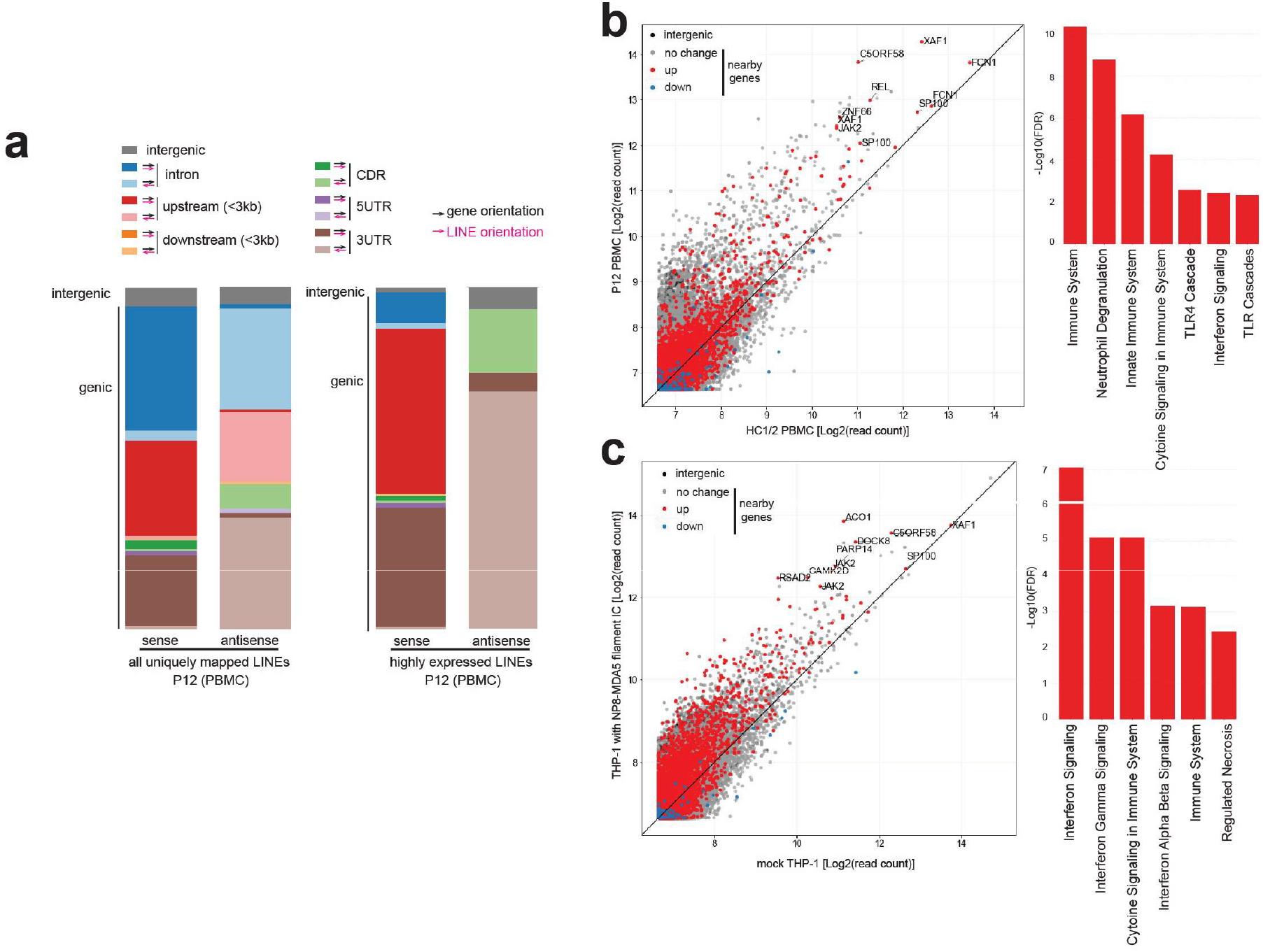
Characterization of LINE RNAs in PBMCs from anti-MDA5 DM patient and macrophages upon IC stimulation. **a**) Annotation of all LINE regions with uniquely mappable reads weighted with read counts (left), and all LINE regions with over 200 uniquely mappable reads in P12 PBMCs (right). Concordant orientation of LINE and gene (darker shading) generates sense LINE RNAs upon gene transcription, whereas discordant orientation (lighter shading) generates antisense LINE RNAs. **b** (Left) Scatterplot of spike-in-normalized read counts of individual LINE loci in P12 PBMCs versus averaged read counts in HC1/2 PBMCs. Each point represents a LINE locus, grouped into those within 3 kb of an upregulated gene (L2FC > 0.3, red), a downregulated gene (L2FC < −0.3, blue), a gene with no change (|L2FC| ≤ 0.3, gray), or not within 3 kb of any gene (black). (Right) Reactome pathway of genes nearby top 100 highly expressed and upregulated LINEs in P12 PBMCs relative to HC1/2. **c** (Left) Scatterplot of reads-per-million (RPM) of individual LINE loci in THP-1 macrophage stimulated with NP8-MDA5 filament ICs versus mock control. Only uniquely mappable reads were counted. Each point is colored as in (b). (Right) Reactome pathway of genes nearby top 100 highly expressed and upregulated LINEs in THP-1 stimulated with NP8-MDA5 filament ICs relative to mock control.

## Extended Data Tables 1-4

**Extended Data Table 1.** Characteristics of anti-MDA5 DM patient from whom antibody-secreting cells were isolated.

|  |  |
| --- | --- |
| **Basic demographics** |  |
| Age | 55 |
| Sex | Male |
| **Cutaneous involvement** |  |
| Gottron's papules | Yes |
| Inverse Gottron's papules | Yes |
| Mechanic's hands | Yes |
| Ulcers | No |
| Facial and truncal erythema | Yes |
| **Other involvement** |  |
| RP-ILD | Yes |
| Myositis | Yes |
| **Laboratory findings** |  |
| Anti-MDA5 autoantibody, Index, (0-31) | 4500 |
| Anti-SS-A autoantibody, U/mL, (0-10) | 25.6 |
| Lymphocyte count, /μL | 645 |
| KL-6, U/mL, (0-464) | 575 |
| SP-D, ng/mL, (0-110) | 26 |
| Creatine kinase, U/L, (59-248) | 1199 |
| Aldolase, U/L, (2.7-7.5) | 13.8 |
| AST, U/L, (13-30) | 142 |
| ALT, U/L, (10-42) | 90 |
| CRP, mg/dL, (0-0.3) | 0.94 |
| ESR, mm/h, (2-10) | 67 |
| Ferritin, ng/mL, (14-304) | 808 |
| **Disease activity score** |  |
| Physician's global disease activity (0-100) | 78 |
| **Sampling** |  |
| Treatment at time of sampling | none |
| Sample type | Fresh PBMC |
| RP-ILD: Rapidly progressive interstitial lung disease |  |
| KL-6: Krebs von den Lungen-6 |  |
| SP-D: surfactant protein-D |  |
| CRP: C-reactive protein |  |

**Extended Data Table 2.** Cryo-EM data collection, refinement, and validation statistics.

|  | MDA5-NP6<br>complex<br>(EMDB- 67964)<br>(PDB 21SJ) | MDA5-NP8<br>complex<br>(EMDB-81614)<br>(PDB 28BD) | MDA5-F1<br>complex<br>(EMDB-68273)<br>(PDB 22HM) | MDA5-F12<br>complex<br>(EMDB- 81615)<br>(PDB 28BE) |
| --- | --- | --- | --- | --- |
| **Data collection and processing** |  |  |  |  |
| Magnification | 105,000 | 105,000 | 105,000 | 105,000 |
| Voltage (kV) | 300 | 300 | 300 | 300 |
| Electron exposure (e<sup>-</sup>/Å<sup>2</sup>) | 50.0 | 50.0 | 59.6 | 50.0 |
| Defocus range (μm) | -0.8 to -2.0 | -0.8 to -2.0 | -1.0 to -2.5 | -0.8 to -2.0 |
| Pixel size (Å) | 0.83 | 0.83 | 0.83 | 0.83 |
| Symmetry imposed | C1 | C1 | C1 | C1 |
| Initial particle images (no.) | 2,597,334 | 3,750,024 | 3,105,681 | 6,481,600 |
| Final particle images (no.) | 415,377 | 96,943 | 257,495 | 195,842 |
| Map resolution (Å) | 2.49 | 2.47 | 3.20 | 2.09 |
| FSC threshold | 0.143 | 0.143 | 0.143 | 0.143 |
| **Refinement** |  |  |  |  |
| Model resolution (Å) | 2.7 | 3.00 | 3.5 | 2.93 |
| FSC threshold | 0.5 | 0.5 | 0.5 | 0.5 |
| Map sharpening *B* factor (Å<sup>2</sup>) | 86.1 | 50.3 | 127.6 | 35.3 |
| Model composition |  |  |  |  |
| Non-hydrogen atoms | 7691 | 7732 | 7530 | 7751 |
| Protein residues | 883 | 894 | 862 | 894 |
| Nucleotide residues | 30 | 30 | 30 | 30 |
| Ligands | 2 | 2 | 2 | 2 |
| *B* factors (Å<sup>2</sup>) |  |  |  |  |
| Protein | 44.52 | 60.59 | 86.37 | 52.49 |
| Ligand | 50.90 | 69.08 | 119.19 | 61.15 |
| R.m.s. deviations |  |  |  |  |
| Bond lengths (Å) | 0.002 | 0.002 | 0.003 | 0.002 |
| Bond angles (°) | 0.559 | 0.499 | 0.490 | 0.489 |
| Validation |  |  |  |  |
| MolProbity score | 1.32 | 1.27 | 1.45 | 1.31 |
| Clashscore | 5.88 | 5.07 | 8.25 | 5.72 |
| Poor rotamers (%) | 0 | 0 | 0.13 | 0 |
| Ramachandran plot |  |  |  |  |
| Favored (%) | 98.04 | 98.64 | 98.81 | 98.53 |
| Allowed (%) | 1.96 | 1.36 | 1.19 | 1.47 |
| Disallowed (%) | 0 | 0 | 0 | 0 |

**Extended Data Table 3.** Demographic and clinical characteristics of dermatomyositis cohort used to characterize extracellular MDA5 filament immune complexes. NA, not applicable.

|  | Dermatomyositis<br>(n = 19) | Healthy<br>(n = 4) |
| --- | --- | --- |
| **Demographics** |  |  |
| Mean age, years (SD) | 54.7 (15.0) | 27.8 (6.6) |
| Female, n (%) | 15 (78.9) | 3 (75.0) |
| **Race, n (%)** |  |  |
| White | 14 (73.7) | 0 (0) |
| Non-white | 5 (26.3) | 4 (100) |
| **Ethnicity, n (%)** |  |  |
| Hispanic | 1 (5.3) | 1 (25) |
| **Disease history** |  |  |
| Disease duration (months), median (IQR) | 21 (36.5) | NA |
| Immunosuppressive treatment, n (%) | 19 (100) | NA |
| Anti-Jo-1, n (%) | 0 (0) | NA |
| Anti-Mi-2, n (%) | 2 (10.5) | NA |
| Anti-PL-7, n (%) | 0 (0) | NA |
| Anti-PL-12, n (%) | 0 (0) | NA |
| Anti-EJ, n (%) | 0 (0) | NA |
| Anti-OJ, n (%) | 0 (0) | NA |
| Anti-SRP, n (%) | 0 (0) | NA |
| Anti-SAE, n (%) | 0 (0) | NA |
| Anti-MDA5, n (%) | 16 (84.2) | NA |
| Anti-NXP2, n (%) | 0 (0) | NA |
| Anti-TIF1y, n (%) | 3 (15.8) | NA |
| Anti-PM/Scl, n (%) | 0 (0) | NA |
| Anti-SS-A, n (%) | 5 (26.3) | NA |
| Anti-Ku, n (%) | 0 (0) | NA |
| Anti-U1RNP, n (%) | 0 (0) | NA |

**Extended Data Table 4.**
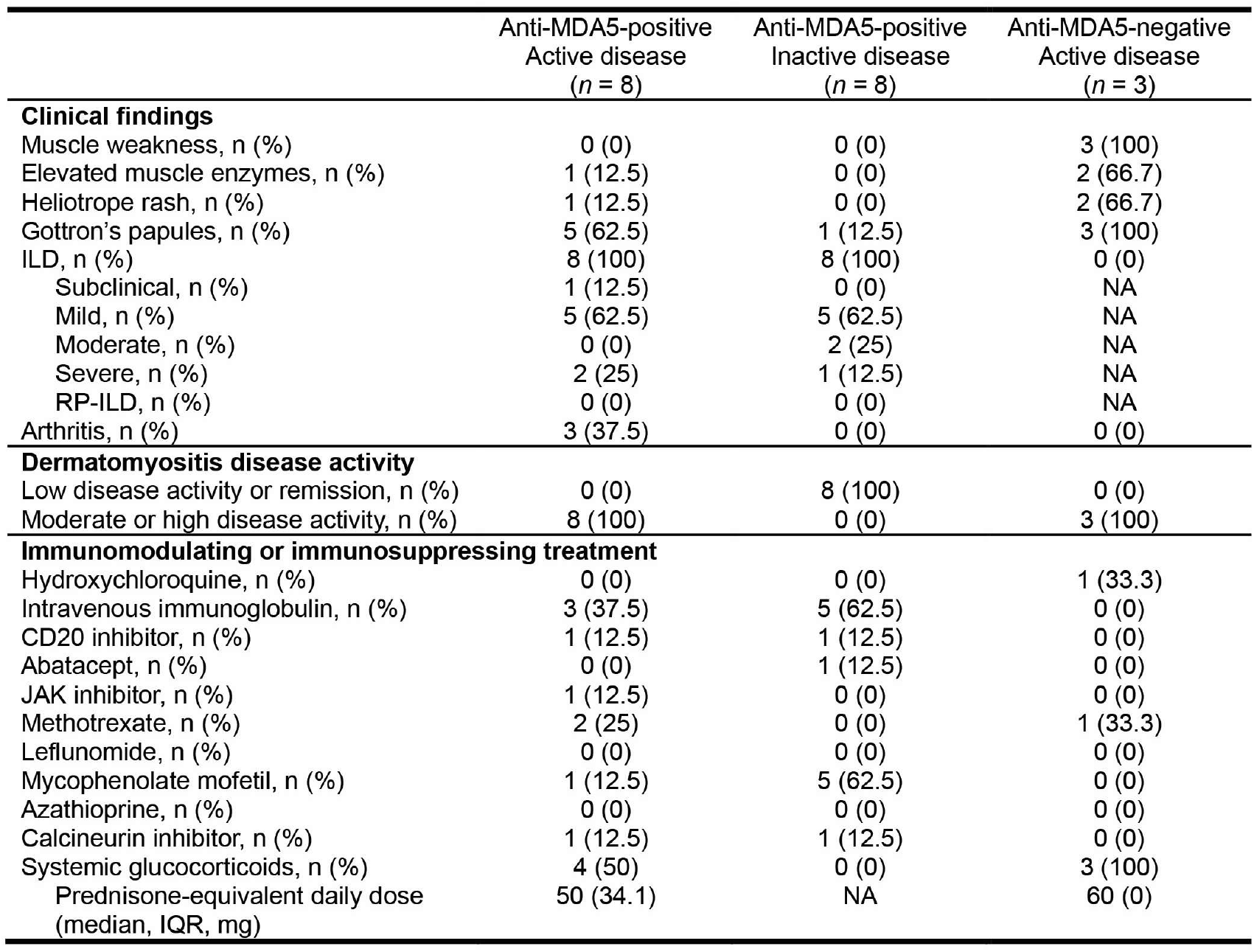
Clinical characteristics of participants based on anti-MDA5 positivity and disease activity at time of sample collection. ILD, interstitial lung disease; RP-ILD, rapidly progressive interstitial lung disease; NA, not applicable.

## References

1 Sato, S. et al. Autoantibodies to a 140-kd polypeptide, CADM-140, in Japanese patients with clinically amyopathic dermatomyositis. Arthritis Rheum 52, 1571–1576 (2005). 10.1002/art.21023

2 Sato, S. et al. RNA helicase encoded by melanoma differentiation-associated gene 5 is a major autoantigen in patients with clinically amyopathic dermatomyositis: Association with rapidly progressive interstitial lung disease. Arthritis Rheum 60, 2193–2200 (2009). 10.1002/art.24621

3 Hur, S. Double-Stranded RNA Sensors and Modulators in Innate Immunity. Annu Rev Immunol 37, 349–375 (2019). 10.1146/annurev-immunol-042718-041356

4 Kurtzman, D. J. B. & Vleugels, R. A. Anti-melanoma differentiation-associated gene 5 (MDA5) dermatomyositis: A concise review with an emphasis on distinctive clinical features. J Am Acad Dermatol 78, 776–785 (2018). 10.1016/j.jaad.2017.12.010

5 Lundberg, I. E. et al. Idiopathic inflammatory myopathies. Nat Rev Dis Primers 7, 86 (2021). 10.1038/s41572-021-00321-x

6 Lu, X., Peng, Q. & Wang, G. Anti-MDA5 antibody-positive dermatomyositis: pathogenesis and clinical progress. Nat Rev Rheumatol 20, 48–62 (2024). 10.1038/s41584-023-01054-9

7 Selva-O’Callaghan, A. et al. Pharmacologic Treatment of Anti-MDA5 Rapidly Progressive Interstitial Lung Disease. Curr Treatm Opt Rheumatol 7, 319–333 (2021). 10.1007/s40674-021-00186-x

8 Ye, Y. et al. Single-cell profiling reveals distinct adaptive immune hallmarks in MDA5+ dermatomyositis with therapeutic implications. Nat Commun 13, 6458 (2022). 10.1038/s41467-022-34145-4

9 Yoshida, A. et al. Dysregulated type I/III interferon system in circulation from patients with anti-MDA5-positive dermatomyositis. Sci Rep 15, 25537 (2025). 10.1038/s41598-025-10895-1

10 Yoshida, T. & Nakashima, R. Anti-Melanoma Differentiation-Associated Gene 5 Antibody Positive Dermatomyositis: Recent Progress in Pathophysiology and Treatment. Curr Rheumatol Rep 27, 23 (2025). 10.1007/s11926-025-01188-7

11 Seto, N. et al. Neutrophil dysregulation is pathogenic in idiopathic inflammatory myopathies. JCI Insight 5 (2020). 10.1172/jci.insight.134189

12 Ichimura, Y. et al. Autoimmunity against melanoma differentiation-associated gene 5 induces interstitial lung disease mimicking dermatomyositis in mice. Proc Natl Acad Sci U S A 121, e2313070121 (2024). 10.1073/pnas.2313070121

13 Pinal-Fernandez, I. et al. Pathogenic autoantibody internalization in myositis. medRxiv (2024). 10.1101/2024.01.15.24301339

14 Ablasser, A. & Hur, S. Regulation of cGAS- and RLR-mediated immunity to nucleic acids. Nat Immunol 21, 17–29 (2020). 10.1038/s41590-019-0556-1

15 Ahmad, S. et al. Breaching self-tolerance to Alu duplex RNA underlies MDA5-mediated inflammation Cell 172, 797–810 (2018).

16 Mehdipour, P. et al. Epigenetic therapy induces transcription of inverted SINEs and ADAR1 dependency. Nature 588, 169–173 (2020). 10.1038/s41586-020-2844-1

17 Sun, T. et al. ADAR1 editing is necessary for only a small subset of cytosolic dsRNAs to evade MDA5-mediated autoimmunity. Nat Genet 57, 3101–3111 (2025). 10.1038/s41588-025-02430-9

18 Peisley, A. et al. Cooperative assembly and dynamic disassembly of MDA5 filaments for viral dsRNA recognition. Proc Natl Acad Sci U S A 108, 21010–21015 (2011). 10.1073/pnas.1113651108

19 Berke, I. C., Yu, X., Modis, Y. & Egelman, E. H. MDA5 assembles into a polar helical filament on dsRNA. Proc Natl Acad Sci U S A 109, 18437–18441 (2012).

20 Wu, B. et al. Structural Basis for dsRNA Recognition, Filament Formation, and Antiviral Signal Activation by MDA5. Cell 152, 276–289 (2013).

21 Kato, K. et al. Structural analysis of RIG-I-like receptors reveals ancient rules of engagement between diverse RNA helicases and TRIM ubiquitin ligases. Mol Cell 81, 599–613 e598 (2021). 10.1016/j.molcel.2020.11.047

22 Hou, F. et al. MAVS forms functional prion-like aggregates to activate and propagate antiviral innate immune response. Cell 146, 448–461 (2011). 10.1016/j.cell.2011.06.041

23 Crow, Y. J. & Stetson, D. B. The type I interferonopathies: 10 years on. Nat Rev Immunol 22, 471–483 (2022). 10.1038/s41577-021-00633-9

24 Van Gompel, E. et al. Anti-MDA5 monoclonal antibodies from patients with dermatomyositis - B cell characteristics and differential targeting of the helicase domains. J Autoimmun 159, 103536 (2026). 10.1016/j.jaut.2026.103536

25 Rice, G. I. et al. Gain-of-function mutations in IFIH1 cause a spectrum of human disease phenotypes associated with upregulated type I interferon signaling. Nat Genet 46, 503–509 (2014). 10.1038/ng.2933

26 Shrock, E. L. et al. Germline-encoded amino acid-binding motifs drive immunodominant public antibody responses. Science 380, eadc9498 (2023). 10.1126/science.adc9498

27 Wu, Y. et al. TLR7/8 Activation in Immune Cells and Muscle by RNA-Containing Immune Complexes: Role in Inflammation and the Pathogenesis of Myositis. Arthritis Rheumatol 77, 190–201 (2025). 10.1002/art.42989

28 Eloranta, M. L. et al. A possible mechanism for endogenous activation of the type I interferon system in myositis patients with anti-Jo-1 or anti-Ro 52/anti-Ro 60 autoantibodies. Arthritis Rheum 56, 3112–3124 (2007). 10.1002/art.22860

29 Vollmer, J. et al. Immune stimulation mediated by autoantigen binding sites within small nuclear RNAs involves Toll-like receptors 7 and 8. J Exp Med 202, 1575–1585 (2005). 10.1084/jem.20051696

30 Hung, T. et al. The Ro60 autoantigen binds endogenous retroelements and regulates inflammatory gene expression. Science 350, 455–459 (2015).

31 Yazdi, A. S. et al. Nanoparticles activate the NLR pyrin domain containing 3 (Nlrp3) inflammasome and cause pulmonary inflammation through release of IL-1alpha and IL-1beta. Proc Natl Acad Sci U S A 107, 19449–19454 (2010). 10.1073/pnas.1008155107

32 Duewell, P. et al. NLRP3 inflammasomes are required for atherogenesis and activated by cholesterol crystals. Nature 464, 1357–1361 (2010). 10.1038/nature08938

33 Martinon, F., Petrilli, V., Mayor, A., Tardivel, A. & Tschopp, J. Gout-associated uric acid crystals activate the NALP3 inflammasome. Nature 440, 237–241 (2006). 10.1038/nature04516

34 Halle, A. et al. The NALP3 inflammasome is involved in the innate immune response to amyloid-beta. Nat Immunol 9, 857–865 (2008). 10.1038/ni.1636

35 Liu, T. et al. Neutrophil-to-lymphocyte ratio is a predictive marker for anti-MDA5 positive dermatomyositis. BMC Pulm Med 22, 316 (2022). 10.1186/s12890-022-02106-8

36 Peng, Y., Zhang, S., Zhao, Y., Liu, Y. & Yan, B. Neutrophil extracellular traps may contribute to interstitial lung disease associated with anti-MDA5 autoantibody positive dermatomyositis. Clin Rheumatol 37, 107–115 (2018). 10.1007/s10067-017-3799-y

37 Bottai, M. et al. EULAR/ACR classification criteria for adult and juvenile idiopathic inflammatory myopathies and their major subgroups: a methodology report. RMD Open 3, e000507 (2017). 10.1136/rmdopen-2017-000507

38 Aggarwal, R. et al. Trial of Intravenous Immune Globulin in Dermatomyositis. N Engl J Med 387, 1264–1278 (2022). 10.1056/NEJMoa2117912

39 Kovacsovics, M. et al. Overexpression of Helicard, a CARD-containing helicase cleaved during apoptosis, accelerates DNA degradation. Curr Biol 12, 838–843 (2002). 10.1016/s0960-9822(02)00842-4

40 Matsushita, T. et al. Antimelanoma differentiation-associated protein 5 antibody level is a novel tool for monitoring disease activity in rapidly progressive interstitial lung disease with dermatomyositis. Br J Dermatol 176, 395–402 (2017). 10.1111/bjd.14882

41 Cao, H. et al. Clinical manifestations of dermatomyositis and clinically amyopathic dermatomyositis patients with positive expression of anti-melanoma differentiation-associated gene 5 antibody. Arthritis Care Res (Hoboken) 64, 1602–1610 (2012). 10.1002/acr.21728

42 Mendez-Dorantes, C. & Burns, K. H. LINE-1 retrotransposition and its deregulation in cancers: implications for therapeutic opportunities. Genes Dev 37, 948–967 (2023). 10.1101/gad.351051.123

43 Richardson, S. R. et al. The Influence of LINE-1 and SINE Retrotransposons on Mammalian Genomes. Microbiol Spectr 3, MDNA3-0061-2014 (2015). 10.1128/microbiolspec.MDNA3-0061-2014

44 Peisley, A. et al. Kinetic mechanism for viral dsRNA length discrimination by MDA5 filaments. Proc Natl Acad Sci U S A 109, E3340–3349 (2012). 10.1073/pnas.1208618109

45 Mustelin, T., Lood, C. & Giltiay, N. V. Sources of Pathogenic Nucleic Acids in Systemic Lupus Erythematosus. Front Immunol 10, 1028 (2019). 10.3389/fimmu.2019.01028

46 Marshak-Rothstein, A. & Rifkin, I. R. Immunologically active autoantigens: the role of toll-like receptors in the development of chronic inflammatory disease. Annu Rev Immunol 25, 419–441 (2007). 10.1146/annurev.immunol.22.012703.104514

47 Wang, K. et al. RNA-Containing Immune Complexes Formed by Anti-Melanoma Differentiation Associated Gene 5 Autoantibody Are Potent Inducers of IFN-alpha. Front Immunol 12, 743704 (2021). 10.3389/fimmu.2021.743704

48 Leadbetter, E. A. et al. Chromatin-IgG complexes activate B cells by dual engagement of IgM and Toll-like receptors. Nature 416, 603–607 (2002). 10.1038/416603a

49 Burbelo, P. D., Iadarola, M. J., Keller, J. M. & Warner, B. M. Autoantibodies Targeting Intracellular and Extracellular Proteins in Autoimmunity. Front Immunol 12, 548469 (2021). 10.3389/fimmu.2021.548469

50 Suurmond, J. & Diamond, B. Autoantibodies in systemic autoimmune diseases: specificity and pathogenicity. J Clin Invest 125, 2194–2202 (2015). 10.1172/JCI78084

51 Zhang, X., Zhang, R. & Yu, J. New Understanding of the Relevant Role of LINE-1 Retrotransposition in Human Disease and Immune Modulation. Front Cell Dev Biol 8, 657 (2020). 10.3389/fcell.2020.00657

52 Ardeljan, D., Taylor, M. S., Ting, D. T. & Burns, K. H. The Human Long Interspersed Element-1 Retrotransposon: An Emerging Biomarker of Neoplasia. Clin Chem 63, 816–822 (2017). 10.1373/clinchem.2016.257444

53 Tokuyama, M. et al. ERVmap analysis reveals genome-wide transcription of human endogenous retroviruses. Proc Natl Acad Sci U S A 115, 12565–12572 (2018). 10.1073/pnas.1814589115

54 Le, E. et al. Type I interferons increase expression of endogenous retrovirus K102 and envelope protein in myeloid cells from patients with autoimmune disease. Mob DNA 16, 36 (2025). 10.1186/s13100-025-00371-y

55 Arteaga-Vazquez, L. J. et al. High interferon response signatures in SLE patient leukocytes are associated with increased transposable element expression in gene introns and intergenic regions. Mob DNA 16, 40 (2025). 10.1186/s13100-025-00377-6

1. Sanjana, N. E., Shalem, O. & Zhang, F. Improved vectors and genome-wide libraries for CRISPR screening. Nat. Methods 11, 783–784 (2014).

2. Peisley, A. et al. Cooperative assembly and dynamic disassembly of MDA5 filaments for viral dsRNA recognition. Proc. Natl. Acad. Sci. 108, 21010–21015 (2011).

3. Concha, J. S. S., Tarazi, M., Kushner, C. J., Gaffney, R. G. & Werth, V. P. The diagnosis and classification of amyopathic dermatomyositis: a historical review and assessment of existing criteria. Br. J. Dermatol. 180, 1001–1008 (2019).

4. Van Gompel, E. et al. Anti-MDA5 monoclonal antibodies from patients with dermatomyositis – B cell characteristics and differential targeting of the helicase domains. J. Autoimmun. 159, 103536 (2026).

5. Lundberg, I. E. et al. 2017 European League Against Rheumatism/American College of Rheumatology classification criteria for adult and juvenile idiopathic inflammatory myopathies and their major subgroups. Ann. Rheum. Dis. 76, 1955–1964 (2017).

6. Brinkmann, V., Laube, B., Abu Abed, U., Goosmann, C. & Zychlinsky, A. Neutrophil extracellular traps: how to generate and visualize them. J. Vis. Exp. JoVE 1724 (2010) doi:10.3791/1724.

7. Yoshimoto, N. et al. An automated system for high-throughput single cell-based breeding. Sci. Rep. 3, 1191 (2013).

8. Meyer, L. et al. A simplified workflow for monoclonal antibody sequencing. PloS One 14, e0218717 (2019).

9. Kurosawa, N., Yoshioka, M., Fujimoto, R., Yamagishi, F. & Isobe, M. Rapid production of antigen-specific monoclonal antibodies from a variety of animals. BMC Biol. 10, 80 (2012).

10. Ohi, M., Li, Y., Cheng, Y. & Walz, T. Negative Staining and Image Classification - Powerful Tools in Modern Electron Microscopy. Biol. Proced. Online 6, 23–34 (2004).

11. Punjani, A., Rubinstein, J. L., Fleet, D. J. & Brubaker, M. A. cryoSPARC: algorithms for rapid unsupervised cryo-EM structure determination. Nat. Methods 14, 290–296 (2017).

12. Bepler, T. et al. Positive-unlabeled convolutional neural networks for particle picking in cryo-electron micrographs. Nat. Methods 16, 1153–1160 (2019).

13. Punjani, A., Zhang, H. & Fleet, D. J. Non-uniform refinement: adaptive regularization improves single-particle cryo-EM reconstruction. Nat. Methods 17, 1214–1221 (2020).

14. Jamali, K. et al. Automated model building and protein identification in cryo-EM maps. Nature 628, 450–457 (2024).

15. Emsley, P., Lohkamp, B., Scott, W. G. & Cowtan, K. Features and development of Coot. Acta Crystallogr. D Biol. Crystallogr. 66, 486–501 (2010).

16. Sanchez-Garcia, R. et al. DeepEMhancer: a deep learning solution for cryo-EM volume post-processing. *Commun*. Biol. 4, 874 (2021).

17. Liebschner, D. et al. Macromolecular structure determination using X-rays, neutrons and electrons: recent developments in *Phenix*. Acta Crystallogr. Sect. Struct. Biol. 75, 861–877 (2019).

18. Williams, C. J. et al. MolProbity: More and better reference data for improved all-atom structure validation. Protein Sci. Publ. Protein Soc. 27, 293–315 (2018).

19. Pettersen, E. F. et al. UCSF ChimeraX: Structure visualization for researchers, educators, and developers. Protein Sci. Publ. Protein Soc. 30, 70–82 (2021).

20. Genin, M., Clement, F., Fattaccioli, A., Raes, M. & Michiels, C. M1 and M2 macrophages derived from THP-1 cells differentially modulate the response of cancer cells to etoposide. BMC Cancer 15, 577 (2015).

21. Bernaerts, E. & Vandooren, J. Protocol to generate human monocyte-derived macrophages in the absence of exogenous proteases and protease inhibitors to study proteolysis. STAR Protoc. 6, 103826 (2025).

22. Bolger, A. M., Lohse, M. & Usadel, B. Trimmomatic: a flexible trimmer for Illumina sequence data. Bioinformatics 30, 2114–2120 (2014).

23. Dobin, A. et al. STAR: ultrafast universal RNA-seq aligner. Bioinformatics 29, 15–21 (2013).

24. Li, H. et al. The Sequence Alignment/Map format and SAMtools. Bioinformatics 25, 2078–2079 (2009).

25. Liao, Y., Smyth, G. K. & Shi, W. The R package Rsubread is easier, faster, cheaper and better for alignment and quantification of RNA sequencing reads. Nucleic Acids Res. 47, e47 (2019).

26. Mudge, J. M. et al. GENCODE 2025: reference gene annotation for human and mouse. Nucleic Acids Res. 53, D966–D975 (2025).

27. Love, M. I., Huber, W. & Anders, S. Moderated estimation of fold change and dispersion for RNA-seq data with DESeq2. Genome Biol. 15, 550 (2014).

28. Xie, Z. et al. Gene Set Knowledge Discovery with Enrichr. Curr. Protoc. 1, e90 (2021).

29. Milacic, M. et al. The Reactome Pathway Knowledgebase 2024. Nucleic Acids Res. 52, D672–D678 (2024).

30. Bao, W., Kojima, K. K. & Kohany, O. Repbase Update, a database of repetitive elements in eukaryotic genomes. Mob. DNA 6, 11 (2015).

31. Lättekivi, F. et al. Transcriptional landscape of human endogenous retroviruses (HERVs) and other repetitive elements in psoriatic skin. Sci. Rep. 8, 4358 (2018).

32. Katoh, K. & Standley, D. M. MAFFT Multiple Sequence Alignment Software Version 7: Improvements in Performance and Usability. Mol. Biol. Evol. 30, 772–780 (2013).

33. Jin, Y., Tam, O. H., Paniagua, E. & Hammell, M. TEtranscripts: a package for including transposable elements in differential expression analysis of RNA-seq datasets. Bioinformatics 31, 3593–3599 (2015).

34. Garza, R. et al. LINE-1 retrotransposons drive human neuronal transcriptome complexity and functional diversification. Sci. Adv. 9, eadh9543 (2023).

35. Adami, A. et al. LINE-1 retrotransposons mediate cis-acting transcriptional control in human pluripotent stem cells and regulate early brain development. Cell Genomics 5, 100979 (2025).

36. Ivancevic, A. et al. Endogenous retroviruses mediate transcriptional rewiring in response to oncogenic signaling in colorectal cancer. Sci. Adv. 10, eado1218 (2024).

37. Nguyen, L. L. et al. Combining EHMT and PARP Inhibition: A Strategy to Diminish Therapy-Resistant Ovarian Cancer Tumor Growth while Stimulating Immune Activation. Mol. Cancer Ther. 23, 1332–1347 (2024).

38. Vandewege, M. W., Patt, R. N., Merriman, D. K., Ray, D. A. & Hoffmann, F. G. The PIWI/piRNA response is relaxed in a rodent that lacks mobilizing transposable elements. RNA 28, 609–621 (2022).

