## Supplementary Figures for "MDA5 multimerization on LINE RNA drives pathogenic extracellular immune complexes in autoimmunity"

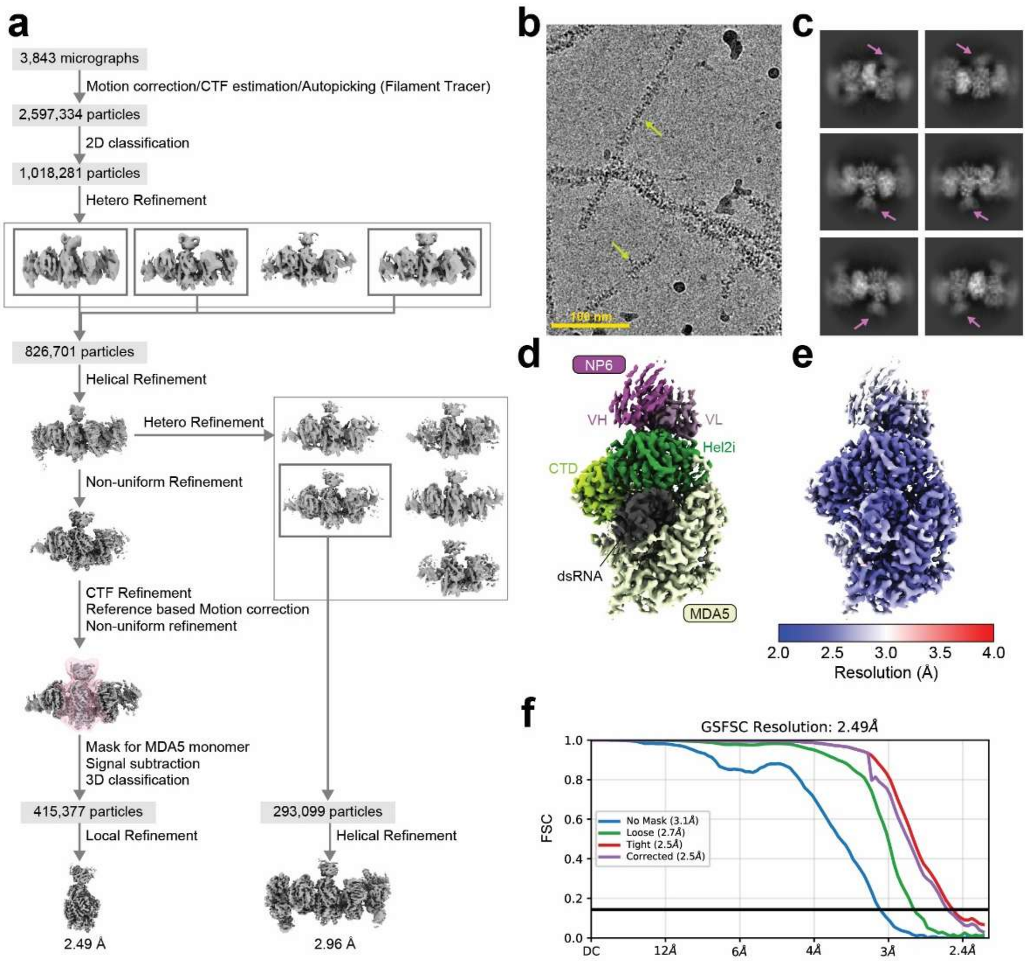


**Supplementary Figure 1. Cryo-EM analysis of the MDA5ΔN:NP6 complex**

a) Cryo-EM image processing workflow.
b) Representative cryo-EM micrograph at a magnification of 105,000x. The MDA5:dsRNA filaments are indicated by light green arrows.
c) Representative 2D class averaged images of particles used for the final reconstruction. The magenta arrows indicate NP6 scFv.
d) Cryo-EM density maps, colored according to the protein domains in Figure 1d.
e) Cryo-EM density maps, colored according to the local resolution.
f) Fourier shell correlation (FSC) curves calculated between the half-maps in the 3D reconstruction.

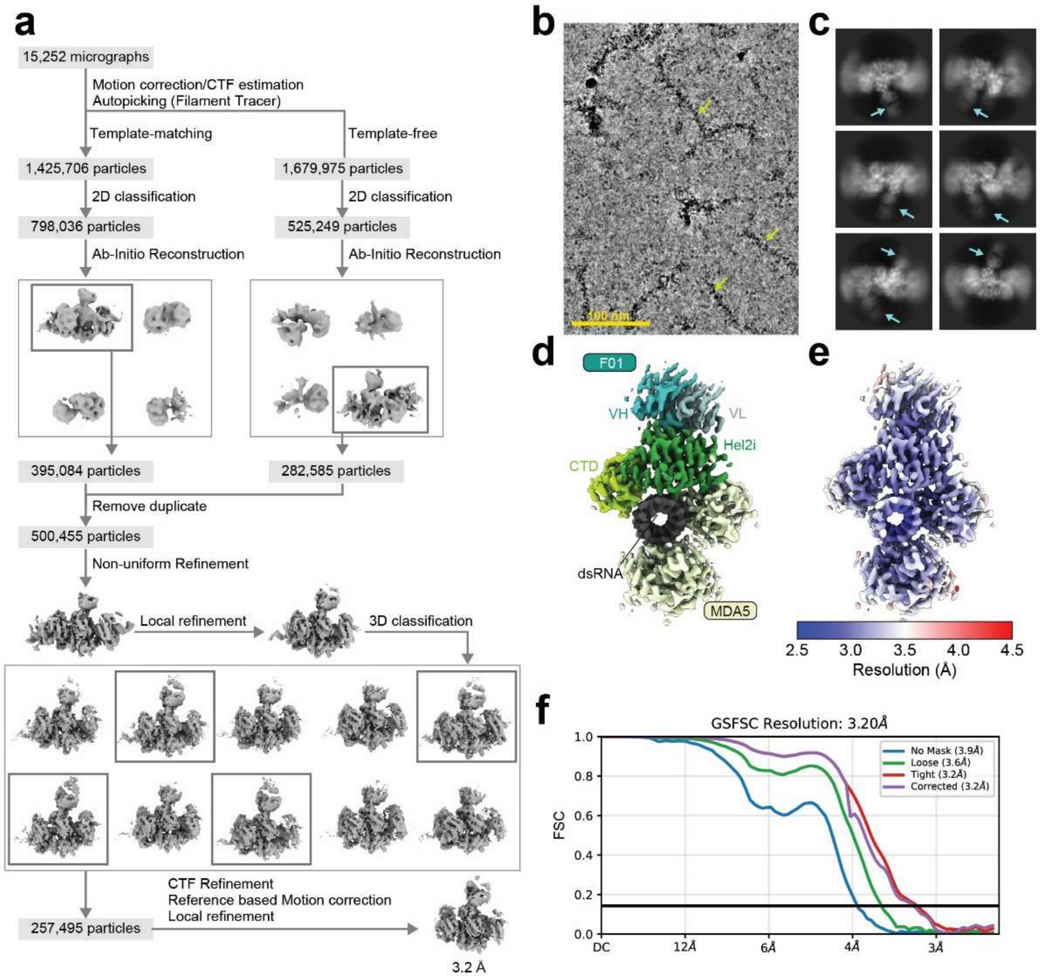


**Supplementary Figure 2. Cryo-EM analysis of the MDA5ΔN:F01 complex**

a) Cryo-EM image processing workflow.
b) Representative cryo-EM micrograph at a magnification of 105,000x. The MDA5:dsRNA filaments are indicated by light green arrows.
c) Representative 2D class averaged images of particles used for the final reconstruction. The cyan arrows indicate F01 Fab.
d) Cryo-EM density maps, colored according to the protein domains in Figure 1d.
e) Cryo-EM density maps, colored according to the local resolution.
f) Fourier shell correlation (FSC) curves calculated between the half-maps in the 3D reconstruction.

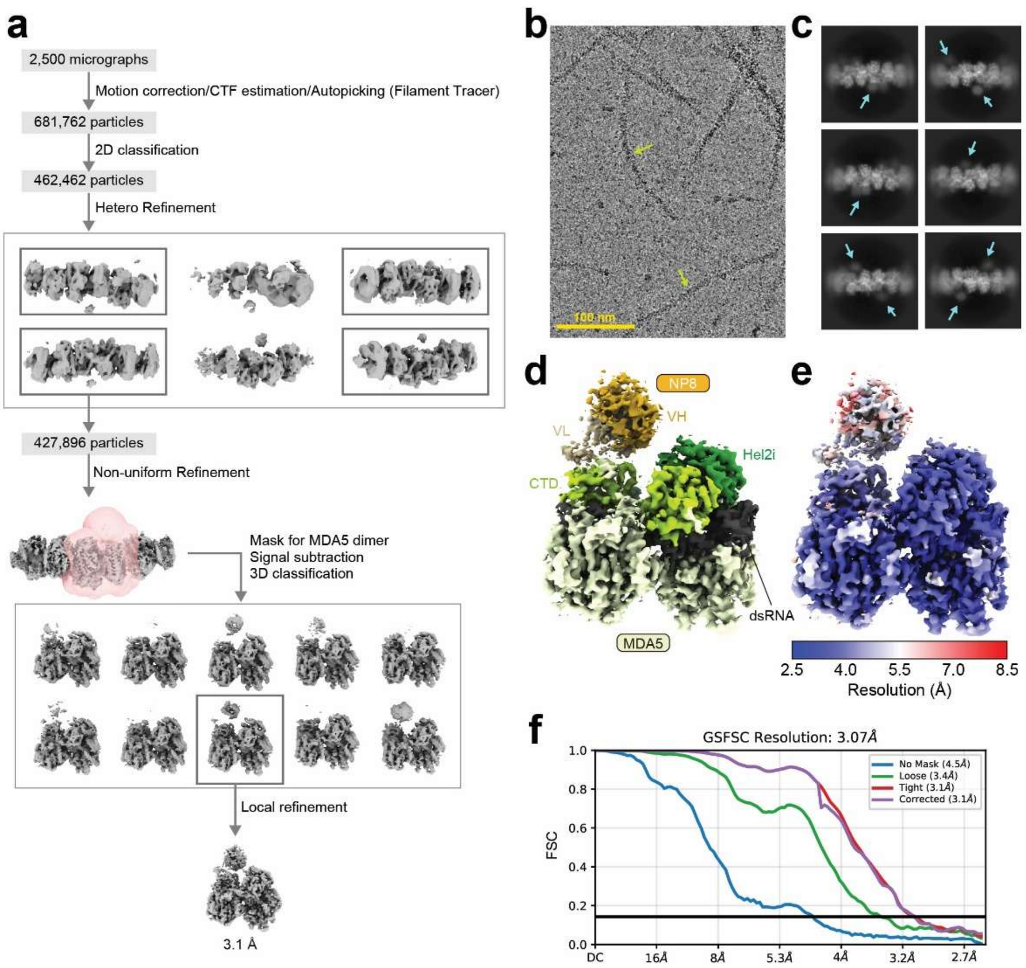


**Supplementary Figure 3. Cryo-EM analysis of the MDA5ΔN:NP8 complex**

a) Cryo-EM image processing workflow.
b) Representative cryo-EM micrograph at a magnification of 105,000x. The MDA5:dsRNA filaments are indicated by light green arrows.
c) Representative 2D class averaged images of particles used for the final reconstruction. The yellow arrows indicate NP8 scFv.
d) Cryo-EM density maps, colored according to the protein domains in Figure 1f.
e) Cryo-EM density maps, colored according to the local resolution.
f) Fourier shell correlation (FSC) curves calculated between the half-maps in the 3D reconstruction.

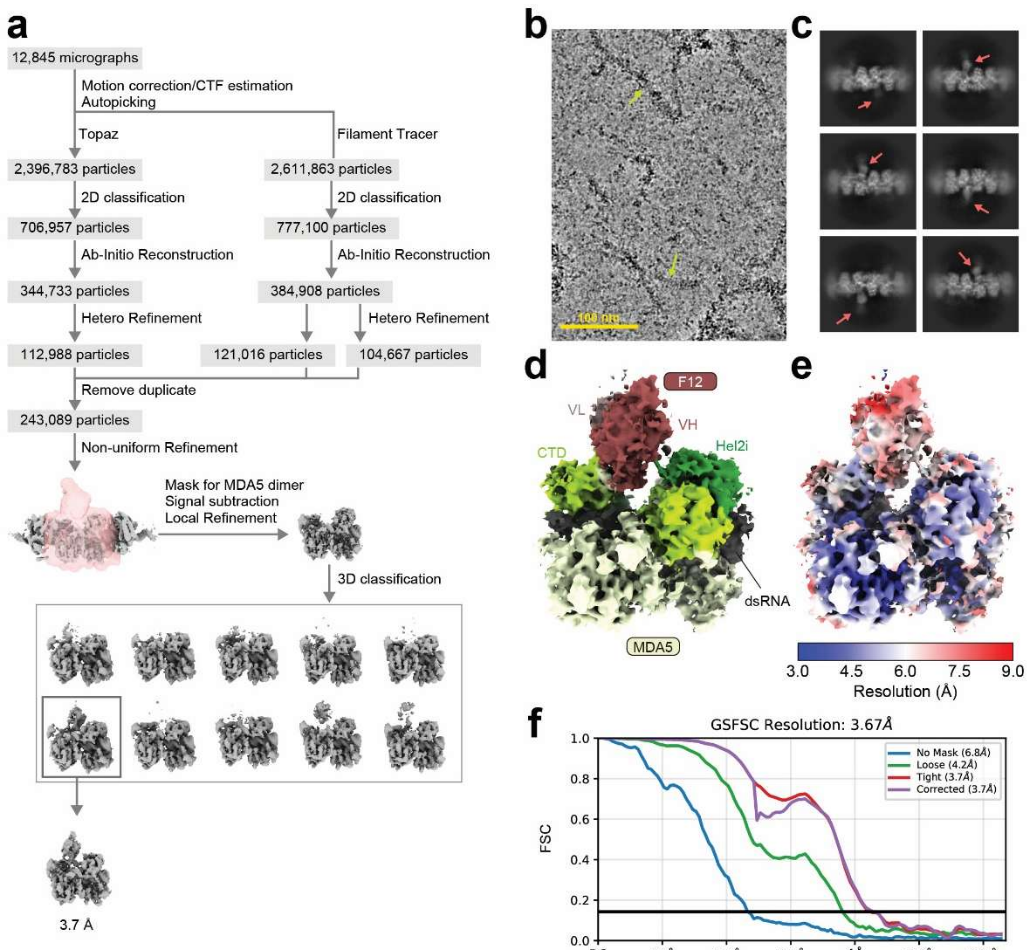


**Supplementary Figure 4. Cryo-EM analysis of the MDA5ΔN:F12 complex**

a) Cryo-EM image processing workflow.
b) Representative cryo-EM micrograph at a magnification of 105,000x. The MDA5:dsRNA filaments are indicated by light green arrows.
c) Representative 2D class averaged images of particles used for the final reconstruction. The red arrows indicate F12 Fab.
d) Cryo-EM density maps, colored according to the protein domains in Figure 1f.
e) Cryo-EM density maps, colored according to the local resolution.
f) Fourier shell correlation (FSC) curves calculated between the half-maps in the 3D reconstruction.

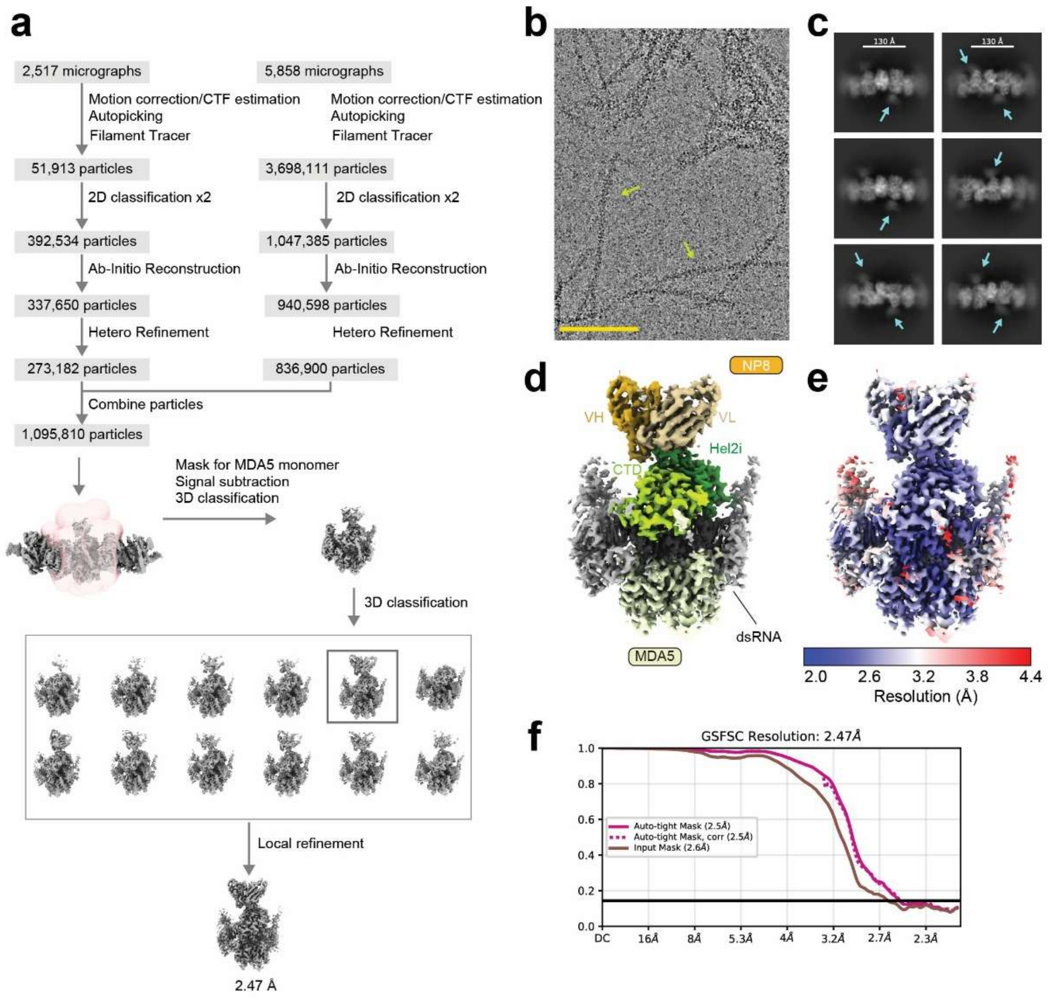


**Supplementary Figure 5. Cryo-EM analysis of the MDA5ΔNΔ640-644Δ657-667:NP8 complex**

a) Cryo-EM image processing workflow.
b) Representative cryo-EM micrograph at a magnification of 105,000x. The MDA5:dsRNA filaments are indicated by light green arrows.
c) Representative 2D class averaged images of particles used for the final reconstruction. The yellow arrows indicate NP8 scFv.
d) Cryo-EM density maps, colored according to the protein domains in Figure 1d.
e) Cryo-EM density maps, colored according to the local resolution.
f) Fourier shell correlation (FSC) curves calculated between the half-maps in the 3D reconstruction.

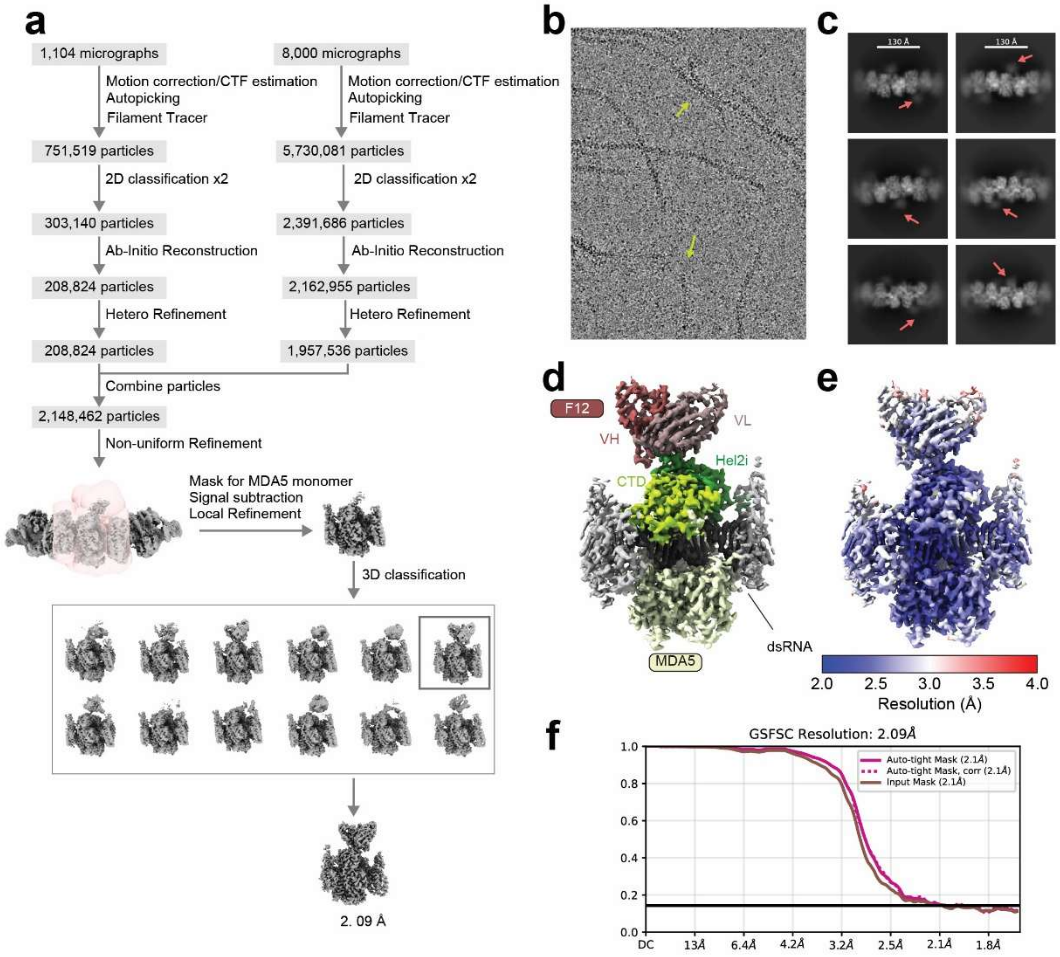


**Supplementary Figure 6. Cryo-EM analysis of the MDA5ΔNΔ640-644Δ657-667:F12 complex**

a) Cryo-EM image processing workflow.
b) Representative cryo-EM micrograph at a magnification of 105,000x. The MDA5:dsRNA filaments are indicated by light green arrows.
c) Representative 2D class averaged images of particles used for the final reconstruction. The red arrows indicate F12 scFv.
d) Cryo-EM density maps, colored according to the protein domains in Figure 1f.
e) Cryo-EM density maps, colored according to the local resolution.
f) Fourier shell correlation (FSC) curves calculated between the half-maps in the 3D reconstruction.
